# Prefrontal signals precede striatal signals for biased credit assignment to (in)actions

**DOI:** 10.1101/2021.10.03.462927

**Authors:** Johannes Algermissen, Jennifer C. Swart, René Scheeringa, Roshan Cools, Hanneke E.M. den Ouden

## Abstract

Actions are biased by the outcomes they can produce: Humans are more likely to show action under reward prospect, but hold back under punishment prospect. Such motivational biases derive not only from biased response selection, but also from biased learning: humans tend to attribute rewards to their own actions, but are reluctant to attribute punishments to having held back. The neural origin of these biases is unclear; in particular, it remains open whether motivational biases arise primarily from the architecture of subcortical regions or also reflect cortical influences, the latter being typically associated with increased behavioral flexibility and emancipation from stereotyped behaviors. Simultaneous EEG-fMRI allowed us to track which regions encoded biased prediction errors in which order. Biased prediction errors occurred in cortical regions (dACC, PCC) before subcortical regions (striatum). These results highlight that biased learning is not a mere feature of the basal ganglia, but arises through prefrontal cortical contributions, revealing motivational biases to be a potentially flexible, sophisticated mechanism.

## Introduction

Human action selection is biased by potential action outcomes: reward prospect drives us to invigorate action, while threat of punishment holds us back^1–3^. These motivational biases have been evoked to explain why humans are tempted by reward-related cues signaling the chance to gain food, drugs, or money, as they elicit automatic approach behavior. Conversely, punishment-related cues suppress action and lead to paralysis, which may even lie at the core of mental health problems such as phobias and mood disorders^4,5^. While such examples highlight the potential maladaptiveness of biases in some situations, they confer benefits in other situations: Biases could provide sensible “default” actions before context-specific knowledge is acquired^1,6^. They may also provide ready-made alternatives to more demanding action selection mechanisms, especially when speed has to be prioritized^7^.

Previous research has assumed that motivational biases arise because the valence of prospective outcomes influences action selection^8^. However, we have recently shown that not only action selection, but also the updating of action values based on obtained outcomes is subject to valence-dependent biases^3,9,10^: humans are more inclined to ascribe rewards to active responses, but have problems with attributing punishments to having held back. On the one hand, such biased learning might be adaptive in combining the flexibility of instrumental learning with somewhat rigid “priors” about typical action-outcome relationships. Exploiting lifetime (or evolutionary) experience might lead to learning that is faster and more robust to environmental “noise”. On the other hand, biases might be responsible for phenomena of “animal superstition” like negative auto-maintenance. Studies of this phenomenon used strict omission schedules in which reward were never delivered on trials on which animals showed an action (key peck, button press), but only when animals inhibited responding over a given time period. Still, animals showed continued key picking in such paradigms, which might either reflect a strong “prior belief” that any situation in which rewards were available requires active work to obtain those, or vice versa an inability to attribute rewards to having held back one’s actions^1,11,12^. While reward attainment can lead to an illusory sense of control over outcomes, control is underestimated under threat of punishment: Humans find it hard to comprehend how inactions can cause negative outcomes, which makes them more lenient in judging harms caused by others’ inactions^13,14^. Taken together, also credit assignment is subject to motivational biases, with enhanced credit for rewards given to actions, but diminished credit for punishments given to inactions.

While evident in behavior, the neural mechanisms subserving such biased credit assignment remain elusive. Previous fMRI studies have studied neural correlates of motivational biases in action selection at the time of cue presentation, finding that the striatal BOLD signal is dominated by the action rather than the cue valence^8,15,16^. More recently, we have reported evidence for cue valence signals in ventromedial prefrontal cortex (vmPFC) and anterior cingulate cortex (ACC), which putatively bias action selection processes in the striatum^17^. The same regions might be involved in motivational biases in learning during outcome processing, given the prominent role of the basal ganglia system not only in action selection, but also learning. Influential computational models of basal ganglia function^18,19^ (henceforth called “asymmetric pathways model”) predict such motivational learning biases: Positive prediction errors, elicited by rewards, lead to long-term potentiation in the striatal direct “Go” pathway (and long term depression in the indirect pathway), allowing for a particularly effective acquisition of Go responses after rewards. Conversely, negative prediction errors, elicited by punishments, lead to long term potentiation in the “NoGo” pathway, impairing the unlearning of NoGo responses after punishments. This account suggests that motivational biases arise within the same pathways involved in standard reinforcement learning (RL). An alternative candidate model is that biases arise through the modulation of these RL systems by external areas that also track past actions, putatively the prefrontal cortex (PFC). Past research has suggested that standard RL can be biased by information stored in PFC, such as explicit instructions^20,21^ or cognitive map-like models of the environment^22–24^. Most notably, the ACC has been found to reflect the impact of explicit instructions^21^ and of environmental changes^25,26^ on prediction errors.

Both candidate models predict that BOLD signal in striatum should be better described by biased compared with “standard” prediction errors. In addition, the model proposing a prefrontal influence on striatal processing makes a notable prediction about the timing of signals: information about the selected action and the obtained outcome should be present first in prefrontal circuits to then later affect processes in the striatum. While fMRI BOLD recordings allow for unequivocal access to striatal activity, the sluggish nature of the BOLD signal prevents clear inferences about temporal precedence of signals from different regions. We thus combined BOLD with simultaneous EEG recordings which allowed us to precisely characterize learning signals in both space and time.

The key question is whether biased credit assignment arises directly from biased RL through the asymmetric pathways in the striatum, or whether striatal RL mechanisms are biased by external prefrontal sources, with the dACC as likely candidate. To this end, participants performed a motivational Go/ NoGo learning task that is well-established to evoke motivational biases^3,9,27^. We expected to observe biased PEs in striatum and frontal cortical areas. By simultaneously recording fMRI and EEG and correlating trial-by-trial BOLD signal with EEG time-frequency power, we were able to time-lock the peaks of EEG-BOLD correlations for regions reflecting biased PEs and infer their relative temporal precedence. We focused on two well-established electrophysiological signatures of RL, namely theta and delta power^28–33^ as well as beta power^28,34^ over midfrontal electrodes.

## Results

Thirty-six participants performed a motivational Go/ NoGo learning task ^3,9^ in which required action (Go/ NoGo) and potential outcome (reward/ punishment) were orthogonalized (Fig. 1A-D). They learned by trial-and-error for each of eight cues whether to perform a left button press (Go_LEFT_), right button press (Go_RIGHT_), or no button press (NoGo), and whether a correct action increased the chance to win a reward (Win cues) or to avoid a punishment (Avoid cues). Correct actions led to 80% positive outcomes (reward, no punishment), with only 20% positive outcomes for incorrect actions. Participants performed two sessions of 320 trials with separate cue sets, which were counterbalanced across participants.

**Figure 1.**
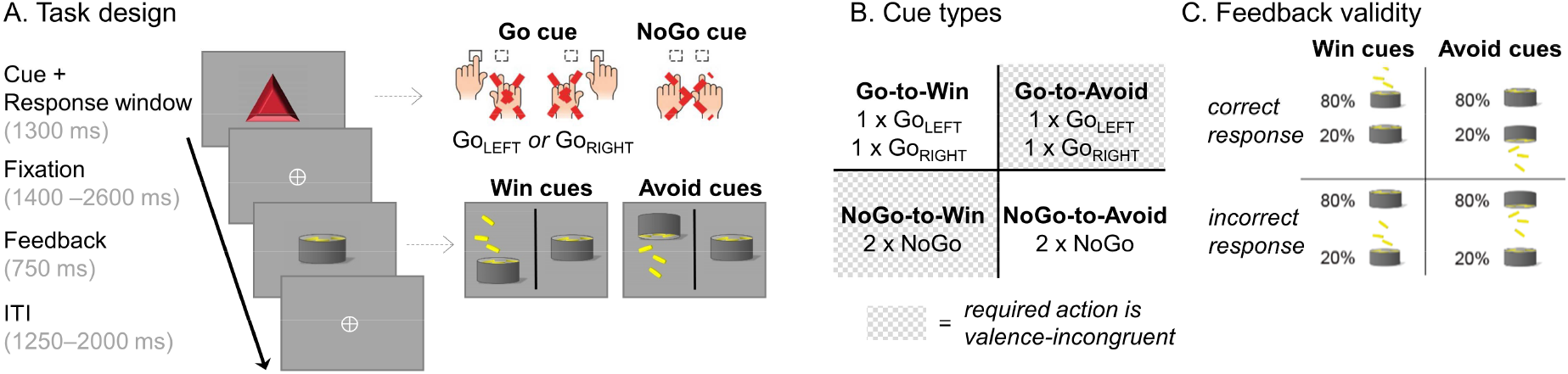
Motivational Go/ NoGo learning task design. **A**. On each trial, a Win or Avoid cue appeared; valence of the cue was not signaled but should be learned. Cue offset was also the response deadline. Response-dependent feedback followed after a jittered interval. Each cue had only one correct action (Go_LEFT_, Go_Right_, or NoGo), which was followed by the positive outcome 80% of the time. For Win cues, actions could lead to rewards or neutral outcomes; for Avoid cues, actions could lead to neutral outcomes or punishments. Rewards and punishments were represented by money falling into/ out of a can. **B.** There were eight different cues, orthogonalizing cue valence (Win versus Avoid) and required action (Go versus NoGo). The motivationally incongruent cues (for which the motivational action tendencies were incongruent with the instrumental requirements) are highlighted in gray. **C.** Feedback was probabilistic: Correct actions to Win cues led to rewards in 80% of cases, but neutral outcomes in 20% of cases. For Avoid cues, correct actions led to neutral outcomes in 80% of cases, but punishments in 20% of cases. For incorrect actions, these probabilities were reversed.

### Regression analyses of behavior

We performed regression analyses to test whether a) responses were biased by the valence of prospective outcomes (Win/ Avoid), reflecting biased responding and/ or learning, and b) whether response repetition after positive vs. negative outcomes was biased by whether a Go vs. NoGo response was performed, selectively reflecting biased learning.

For the first purpose, we analyzed choice data (Go/ NoGo) using mixed-effects logistic regression that included the factors required action (Go/ NoGo; note that this approach collapses across Go_LEFT_ and Go_RIGHT_ responses), cue valence (Win/ Avoid), and their interaction (also reported in)^17^. Participants learned the task, i.e., they performed more Go responses towards Go than NoGo cues (main effect of required action: *b* = 0.815, *SE* = 0.113, χ^2^(1) = 32.008, *p* < .001). In contrast to previous studies ^3,9^, learning did not asymptote (Fig. 2A), which provided greater dynamic range for the biased learning effects to surface. Furthermore, participants showed a motivational bias, i.e., they performed more Go responses to Win than Avoid cues (main effect of cue valence, *b* = 0.423, *SE* = 0.073, χ^2^(1) = 23.695, *p* < .001). Replicating other studies with this task, there was no significant interaction between required action and cue valence (*b* = 0.030, *SE* = 0.068, χ^2^(1) = 0.196, *p* = .658, Fig. 2A-B), i.e., there was no evidence for the effect of cue valence (motivational bias) differing in size between Go or NoGo cues.

**Figure 2.**
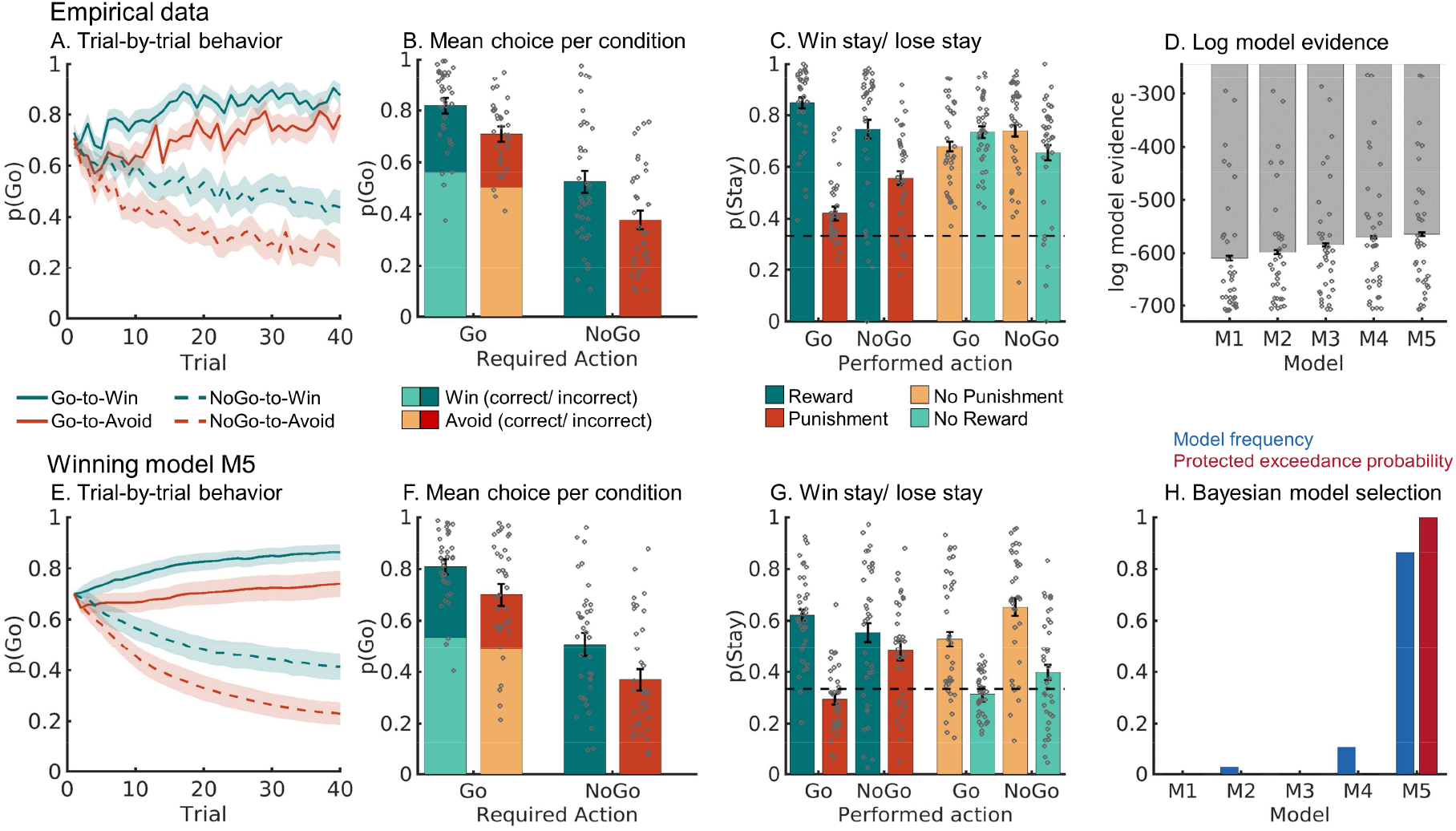
Behavioral performance. **A.** Trial-by-trial proportion of Go responses (±SEM across participants) for Go cues (solid lines) and NoGo cues (dashed lines). The motivational bias was already present from very early trials onwards, as participants made more Go responses to Win than Avoid cues (i.e., green lines are above red lines). Additionally, participants clearly learn whether to make a Go response or not (proportion of Go responses increases for Go cues and decreases for NoGo cues). **B**. Mean (±SEM across participants) proportion Go responses per cue condition (points are individual participants’ means). **C.** Probability to repeat a response (“stay”) on the next encounter of the same cue as a function of action and outcome. Learning was reflected in higher probability of staying after positive outcomes than after negative outcomes (main effect of outcome valence). Biased learning was evident in learning from salient outcomes, where this valence effect was stronger after Go responses than NoGo responses. Dashed line indicates chance level choice (p_Stay_ = 0.33). **D.** Log-model evidence favors the asymmetric pathways model (M5) over simpler models (M1-M4). **E-G.** Trial-by-trial proportion of Go responses, mean proportion Go responses, and probability of staying based on one-step-ahead predictions using parameters (hierarchical Bayesian inference) of the winning model (asymmetric pathways model, M5). **H.** Model frequency and protected exceedance probability indicate best fit for model M5 (asymmetric pathways model), in line with log model evidence.

Secondly, as a proxy of (biased) learning, we analyzed cue-based response repetition (i.e., the probability of repeating a response on the next encounter of the same cue) as a function of outcome valence (positive vs negative outcome), performed action (Go vs. NoGo), and outcome salience (salient: reward or punishment vs. neutral: no reward or no punishment). As expected, participants were more likely to repeat the same response following a positive outcome (main effect of outcome valence: *b* = 0.504, *SE* = 0.053, χ^2^(1) = 45.595, *p* < .001). Most importantly, after salient outcomes, participants adjusted their responses to a larger degree following Go responses than NoGo responses, revealing the presence of a learning bias (Fig. 2C; interaction of valence x action x salience: *b* = 0.248, *SE* = 0.048, χ^2^(1) = 19.732, *p* < .001). When selectively analyzing trials with salient outcomes only, rewards (compared to punishments) led to a higher proportion of choice repetitions following Go relative to NoGo responses (valence x response: *b* = 0.308, *SE* = 0.064, χ^2^(1) = 17.798, *p* < .001; valence effect for Go only: *b* = 1.276, *SE* = 0.115, χ^2^(1) = 53.932, *p* < .001; valence effect for NoGo only: *b* = 0.637, *SE* = 0.127, χ^2^(1) = 18.228, *p* < .001; see full results in Supplementary Table 1).

Taken together, these results suggested that behavioral adaptation following rewards and punishments was biased by the type of action that led to this outcome (Go or NoGo). However, this analysis only considered behavioral adaptation on the next trial, and could not pinpoint the precise algorithmic nature of this learning bias. More importantly, it did not provide trial-by-trial estimates of action values as required for model-based fMRI and EEG analyses to test for regions or time points that reflected biased learning. We thus analyzed the impact of past outcomes on participants’ choices using computational RL models.

### Computational modeling of behavior

In line with previous work^3,9^, we fitted a series of increasingly complex RL models. We started with a simple Rescorla Wagner model featuring learning rate and feedback sensitivity parameters (M1). We next added a Go bias, capturing participants’ overall propensity to make Go responses (M2), and a Pavlovian response bias (M3), reflecting participants’ propensity to adjust their likelihood of emitting a Go response in response to Win vs. Avoid cues^3^. Alternatively, we added a learning bias (M4), amplifying the learning rate after rewarded Go responses and dampening it after punished NoGo responses^3^, in line with the asymmetric pathways model. In the final model (M5), we added both the response bias and the learning bias. For the full model space (M1-M5) and model definitions, see the Methods section.

Model comparison showed clear evidence in favor of the full asymmetric pathways model featuring both response and learning biases (M5; model frequency: 86.43%, protected exceedance probability: 100%, see Fig. 2D, H; for model parameters and fit indices, see Supplementary Table 2; for parameter recovery analyses, see Supplementary Note 6 and Supplementary Fig. 5). Posterior predictive checks involving one-step-ahead predictions and model simulations showed that this model captured key behavioral features (Fig. 2E, F), including motivational biases and a greater behavioral adaptation after Go responses followed by salient outcomes than after NoGo responses followed by salient outcomes (Fig. 2G). This pattern could not be captured by an alternative learning bias model based on the idea that active responses generally enhance credit assignment^35^ (Supplementary Note 7 and Supplementary Fig. 6).

One feature of the behavioral data that was not well captured by the asymmetric pathways model was a high tendency of participants to repeat responses (“stay”) to the same cue irrespective of outcomes (see Fig. 2C and G). This tendency was stronger for Win than Avoid cues. We explored three additional models featuring supplementary mechanisms to account for this behavioral pattern (Supplementary Note 8 and Supplementary Fig. 7). All these models fitted the data well and captured the propensity of staying better than M5; however, these models overestimated the proportion of incorrect Go responses. Model-based fMRI analyses based on these models led to results largely identical to those obtained with M5 (Supplementary Note 9 and Supplementary Fig. 8). We thus focused on M5, which relied on only a single mechanism (i.e., biased learning from rewarded Go and punishment NoGo actions).

### fMRI: Basic quality control analyses

First, we performed a GLM as a quality-check to test which regions encoded positive (rewards, no punishments) vs. negative (no reward/ punishment) outcomes in a “model-free” way, independent of any model-based measure derived from a RL model (for full description of the GLM regressors and contrasts, see Supplementary Table 4). Positive outcomes elicited a higher BOLD response in regions including vmPFC, ventral striatum, and right hippocampus, while negative outcomes elicited higher BOLD in bilateral dorsolateral PFC (dlPFC), left ventrolateral PFC, and precuneus (Fig. 3A, see full report of significant clusters in Supplementary Table 6).

**Figure 3.**
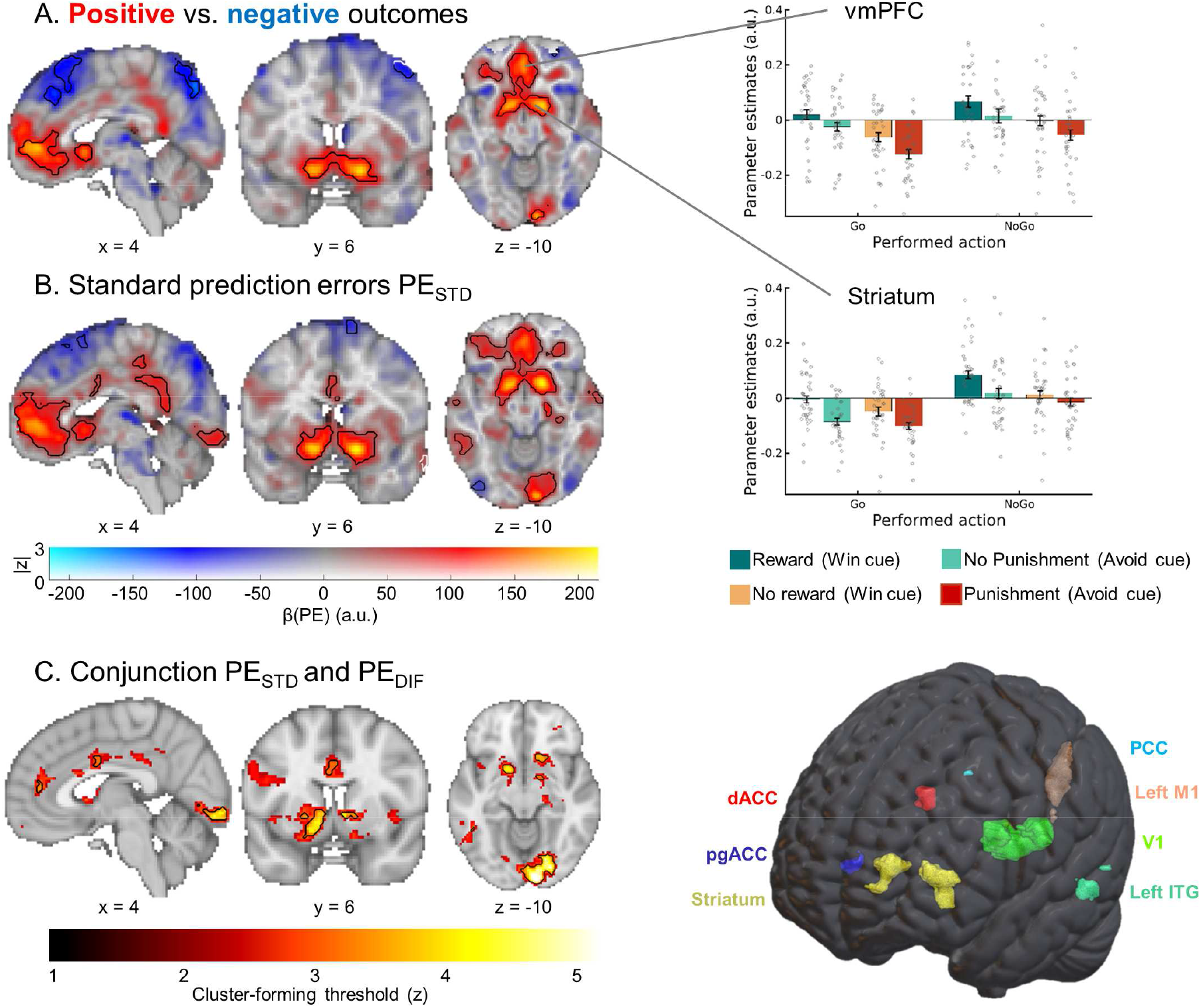
BOLD signal reflecting outcome processing. BOLD effects displayed using a dual-coding visualization: color indicates the parameter estimates and opacity the associated z-statistics. Significant clusters are surrounded by black edges. **A.** Significantly higher BOLD signal for positive outcomes (rewards, no punishments) compared with negative outcomes (no rewards, punishments) was present in a range of regions including bilateral ventral striatum and vmPFC. Bar plots show mean parameter estimates per condition (±SEM across participants; dots indicating individual participants) **B.** BOLD signals correlated positively to “standard” RL prediction errors in several regions, including the ventral striatum, dACC, vmPFC, and PCC. **C.** Left panel: Regions encoding both the standard PE term and the difference term to biased PEs (conjunction) at different cluster-forming thresholds (1 < z < 5, color coding; opacity constant). Clusters significant at a threshold of z > 3.1 are surrounded by black edges. In bilateral striatum, dACC, pgACC, PCC, left motor cortex, left inferior temporal gyrus, and primary visual cortex, BOLD was significantly better explained by biased learning than by standard learning. Right panel: 3D representation with all seven regions encoding biased learning (and used in fMRI-informed EEG analyses).

We also assessed which regions encoded Go vs. NoGo as well as Go_LEFT_ vs. Go_RIGHT_ responses. There was higher BOLD for Go than NoGo responses at the time of response in dorsal ACC (dACC), striatum, thalamus, motor cortices, and cerebellum, while BOLD was higher for NoGo than Go responses in right IFG (Fig. 6C left panel; Supplementary Table 6)^17^. For lateralized Go responses, there was higher BOLD signal in contralateral motor cortex and operculum as well as ipsilateral cerebellum when contrasting hand responses against each other (Fig. 6C, right panel). These results are in line with previous results on outcome processing and response selection and thus assure the general data quality.

**Figure 4.**
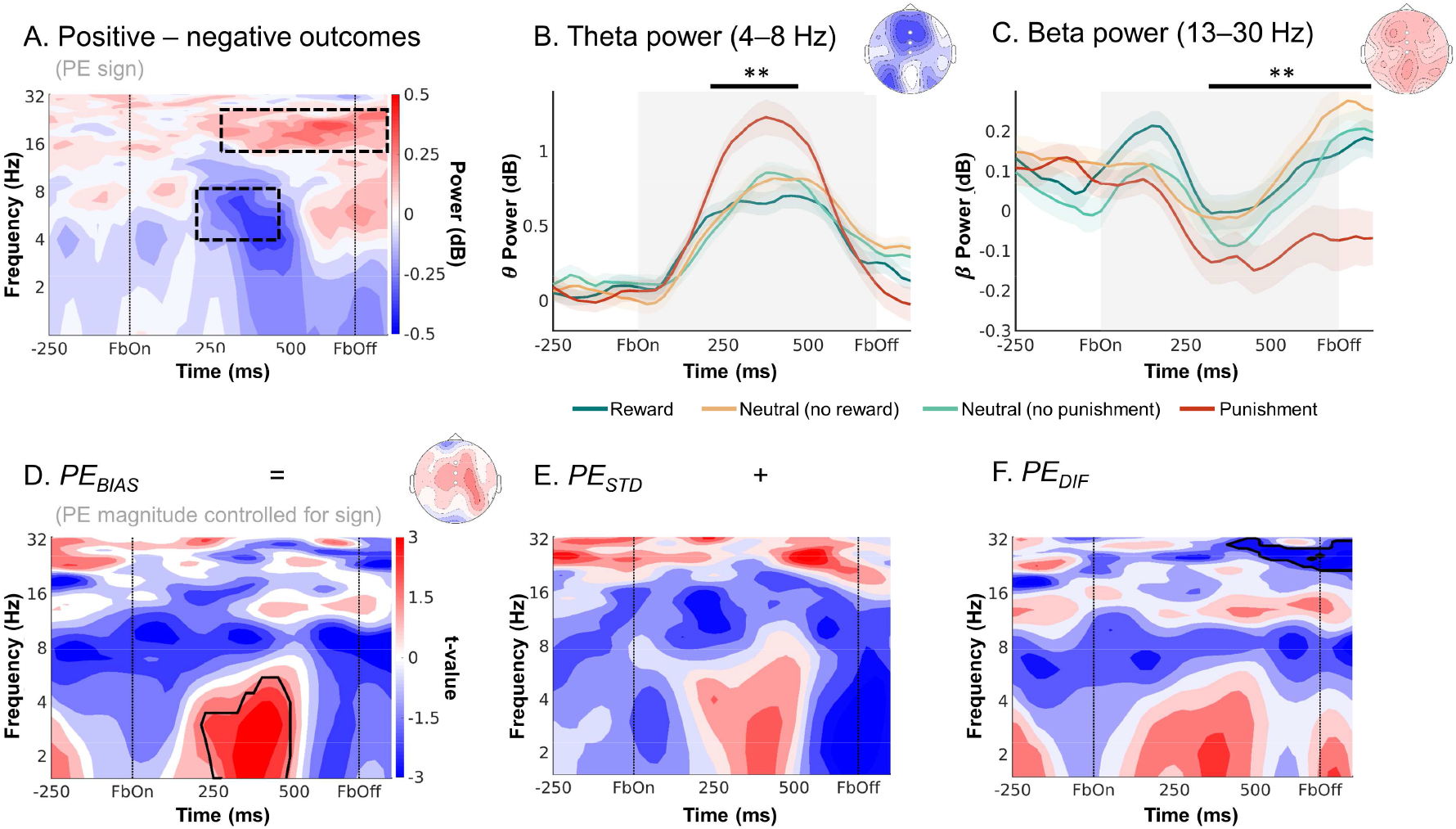
EEG time-frequency power over midfrontal electrodes (Fz/ FCz/ Cz). reflecting outcome processing. **A.** Time-frequency plot (logarithmic y-axis) displaying higher theta (4–8 Hz) power for negative (non-reward for Win cues and punishment for Avoid cues) outcomes and higher beta power (16–32 Hz) for positive (reward and non-punishment) outcomes. This contrast reflects EEG correlates of PE valence (better vs. worse than expected). Black square dot boxes indicate clusters above threshold that drive significance in a-priori defined frequency ranges. **B.** Theta power transiently increases for any outcome, but more so for negative outcomes (especially punishments) around 225–475 ms after feedback onset. Black horizontal lines indicate the time range for which the cluster driving significance was above threshold. (C) Beta power was higher for positive than negative outcomes over a long time period around 300– 1,250 ms after feedback onset. **D-F.** Correlations between midfrontal EEG power and model-based trial-by-trial PE magnitudes controlling for PE valence (thus effectively testing for correlates of “absolute” PEs). Panel **D** displays the correlates of biased prediction errors *PE_BIAS_*, which are decomposed into (**E**) *PE_STD_* based on the non-biased learning model M1, and (**F**) their difference *PE_DIF_.* Solid black lines indicate clusters above threshold. Biased PEs were significantly positively correlated with midfrontal delta power (**D**). The correlations of delta with the standard PEs (**E**) and the difference term to biased PEs (**F**) were positive as well, though not significant. Beta power only significantly encoded the difference term to biased PEs (F). ** *p* < 0.01.

### fMRI: Biased learning in prefrontal cortex and striatum

To test which brain regions were involved in biased learning, we performed a model-based GLM featuring the trial-by-trial PE update as a parametric regressor (see GLM notation in Supplementary Table 3). We used the group-level parameters of the best fitting computational model (M5) to compute trial-by-trial belief updates (i.e., prediction error * learning rate) for every trial for every participant. In assessing neural signatures of biased learning, we faced the complication that standard (Rescorla-Wagner learning in M1) and biased PEs (winning model M5) were highly correlated. A mean correlation of 0.92 across participants (range 0.88–0.95) made it difficult to neurally distinguish biased from standard learning. To circumvent this collinearity problem, we decomposed the biased PE (computed using model M5) into the standard PE (computed using model M1) plus a difference term^22,36^:

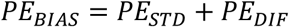

A neural signature of biased learning should significantly—and with the same sign—encode both components of this biased PE term. Standard PEs and the difference term were uncorrelated (mean correlation of −0.02 across participants; range −0.33–0.24; see Supplementary Fig. 9 and 10 for a graphical illustration of this procedure). We tested for biased prediction errors *PE_BIAS_* by testing which regions significantly encoded the conjunction of both its components, i.e., the significant encoding of both *PE_STD_* and *PE_DIF_*. Dissociating two alternative learning signals by decomposing one into the other plus a difference term is an established procedure to disentangle the contributions of two highly correlated signals^22,36^. It has an effect highly similar to orthogonalizing regressors^37^ while maintaining interpretability in that both regressors (*PE_STD_* and *PE_DIF_*) add up to the term of interest (*PE_BIAS_*). Significant encoding of both components (with the same sign) provides strong evidence for encoding of biased prediction errors *PE_BIAS_*. The *PE_DIF_* term itself has no substantive neural interpretation; it is merely an implicit model comparison of a null model (*PE_STD_*) against a full model (*PE_BIAS_*). Intuitively, for voxels for which both *PE_STD_* and *PE_DIF_* are significant, one can conclude that the BOLD signal correlates with the full biased prediction error term *PE_BIAS_*, and that this correlation is significantly stronger than for the baseline prediction error term *PE_STD_*.

While *PE_STD_* was encoded in a range of cortical and subcortical regions (Fig. 3B) previously reported in the literature^38^, significant encoding of both *PE_STD_* and PE_DIF_ (conjunction) occurred in striatum (caudate, nucleus accumbens), dACC (area 23/24), perigenual ACC (pgACC; area 32d bordering posterior vmPFC), posterior cingulate cortex (PCC), left motor cortex, left inferior temporal gyrus, and early visual regions (Fig. 3C; see full report of significant clusters in Supplementary Table 5). Thus, BOLD signal in these regions was better described (i.e., more variance explained) by biased learning than by standard prediction error learning.

### EEG: Biased learning in midfrontal delta, theta, and beta power

Similar to the fMRI analyses, we next tested whether midfrontal power encoded biased PEs rather than standard PEs. While fMRI provides spatial specificity of where PEs are encoded, EEG power provides temporal specificity of when signals encoding prediction errors occur^29,34^. In line with our fMRI analysis, we used the standard PE term *PE_STD_* and the difference to the biased PE term *PE_DIF_* as trial-by-trial regressors for EEG power at each channel-time-frequency bin for each participant and then performed cluster-based permutation tests across the *b*-maps of all participants. Note that differently from BOLD signal, EEG signatures of learning typically do not encode the full prediction error. Instead, PE valence (better vs. worse than expected) and PE magnitude (saliency, surprise) have been found encoded in the theta and delta band, respectively, but with opposite signs^31–33^. When testing for parametric correlates of PE magnitude, we therefore controlled for PE valence, thereby effectively testing for correlations with the absolute PE magnitude (i.e., degree of surprise). Note that PE valence was identical for standard and biased PEs. Thus, only PE magnitude could distinguish both learning models.

Both midfrontal theta and beta power reflected outcome (PE) valence: Theta power was higher for negative (non-reward and punishment) than for positive (reward and non-punishment) outcomes (225–475 ms, *p* = .006; Fig. 4A-B), while beta power was higher for positive than for negative outcomes (300–1,250 ms, *p* = .002; Fig. 4A, C). Differences in theta power were clearly strongest over frontal channels, while differences in the beta range were more diffuse, spreading over frontal and parietal channels (Fig. 4B-C). All results held when the condition-wise ERP was removed from the data (see Supplementary Note 10 and Supplementary Fig. 13), suggesting that differences between conditions were due to induced (rather than evoked) activity (for results in the time domain, see Supplementary Note 11 and Supplementary Fig. 14 and 15).

When testing for correlates of PE magnitude, we controlled for PE valence given that previous studies have reported TF correlates of both PE valence and PE magnitude in a similar time and frequency range, but with opposite signs^31–33^. Midfrontal delta power was indeed positively correlated with the *PE_BIAS_* term (225–475 ms; *p* = .017; Fig. 4D). Decomposition of the *PE_BIAS_* term into its constituent terms showed that this correlation was not significant for the *PE_STD_* term (*p* = 0.074, Fig. 4E) nor for the *PE_DIF_* term (*p* = 0.185; Fig. 4F). This result does not imply that the *PE_BIAS_* term explained delta power significantly better than the *PE_STD_* term; it only implies significant encoding of the *PE_BIAS_* term as suggested by the model that best fitted the behavioral data, with no significant evidence for a similar encoding of the conventional *PE_STD_* term. For a similar observation in the time-domain EEG signal, see Supplementary Note 12 and Supplementary Fig. 16. Beyond delta power, beta power correlated positively, though not significantly with *PE_STD_* (*p* = 0.110, Fig. 4E) and significantly negatively with *PE_DIF_* (*p* = .001, 425–850 ms). Given these oppositely-signed correlations of its constituents, the *PE_BIAS_* term did not significantly correlate with beta power (*p* = 0.550, Fig 4D).

In sum, both midfrontal theta power (negatively) and beta power (positively) encoded PE valence. In addition, delta power encoded PE magnitude (positively). This encoding was only significant for biased PEs, but not standard PEs. Taken together, as was the case for BOLD signal, midfrontal EEG power also reflected biased learning. As a next step, we tested whether the identified EEG phenomena were correlated with trial-by-trial BOLD signal in identified regions. Crucially, this allowed us to test whether EEG correlates of cortical learning precede EEG correlates of subcortical learning.

### Combined EEG-fMRI: Prefrontal cortex signals precede striatum during biased outcome processing

The observation that also cortical areas (dACC, pgACC, PCC) show biased PEs is consistent with the “external model” of cortical signals biasing learning processes in the striatum. However, this model makes the crucial prediction that these biased learning signals should be present first in cortical areas and only later in the striatum. Next, we used trial-by-trial BOLD signal from those regions encoding biased PE to predict midfrontal EEG power. By determining the time points at which different regions correlated with EEG power, we were able to infer the relative order of biased PE processing across cortical and subcortical regions, revealing whether cortical processing preceded striatal processing. We used trial-by-trial BOLD signal from the seven regions encoding biased PEs, i.e., striatum, dACC, pgACC, PCC, left motor cortex, left ITG, and primary visual cortex (see masks in Supplementary Fig. 11 and 12) as regressors on average EEG power over midfrontal electrodes (Fz/ FCz/ Cz; see Supplementary Fig. 17 for a graphical illustration of this approach). We performed analyses with and without PEs included in the model, which yielded identical results and suggested that EEG-fMRI correlations did not merely result from PE processing as a “common cause” driving signals in both modalities. Instead, EEG-fMRI correlations reflected incremental variance explained in EEG power by the BOLD signal in selected regions (even beyond variance explained by the model-based PE estimates), providing the strongest test for the hypothesis that BOLD and EEG signal reflect the same neural phenomenon. As the timeseries of all seven regions were included in one single regression, their regression weights reflected each region’s unique contribution, controlling for any shared variance. In line with the “external model”, BOLD signal from prefrontal cortical regions correlated with midfrontal EEG power earlier after outcome onset than did striatal BOLD signal:

First, dACC BOLD was significantly negatively correlated with alpha/ theta power early after outcome onset (100–575 ms, 2–17 Hz, *p* = .016; Fig. 5A). This cluster started in the alpha/ theta range and then spread into the theta/delta range (henceforth called “lower alpha band power”). It was not observed in the EEG-only analyses reported above.

**Figure 5.**
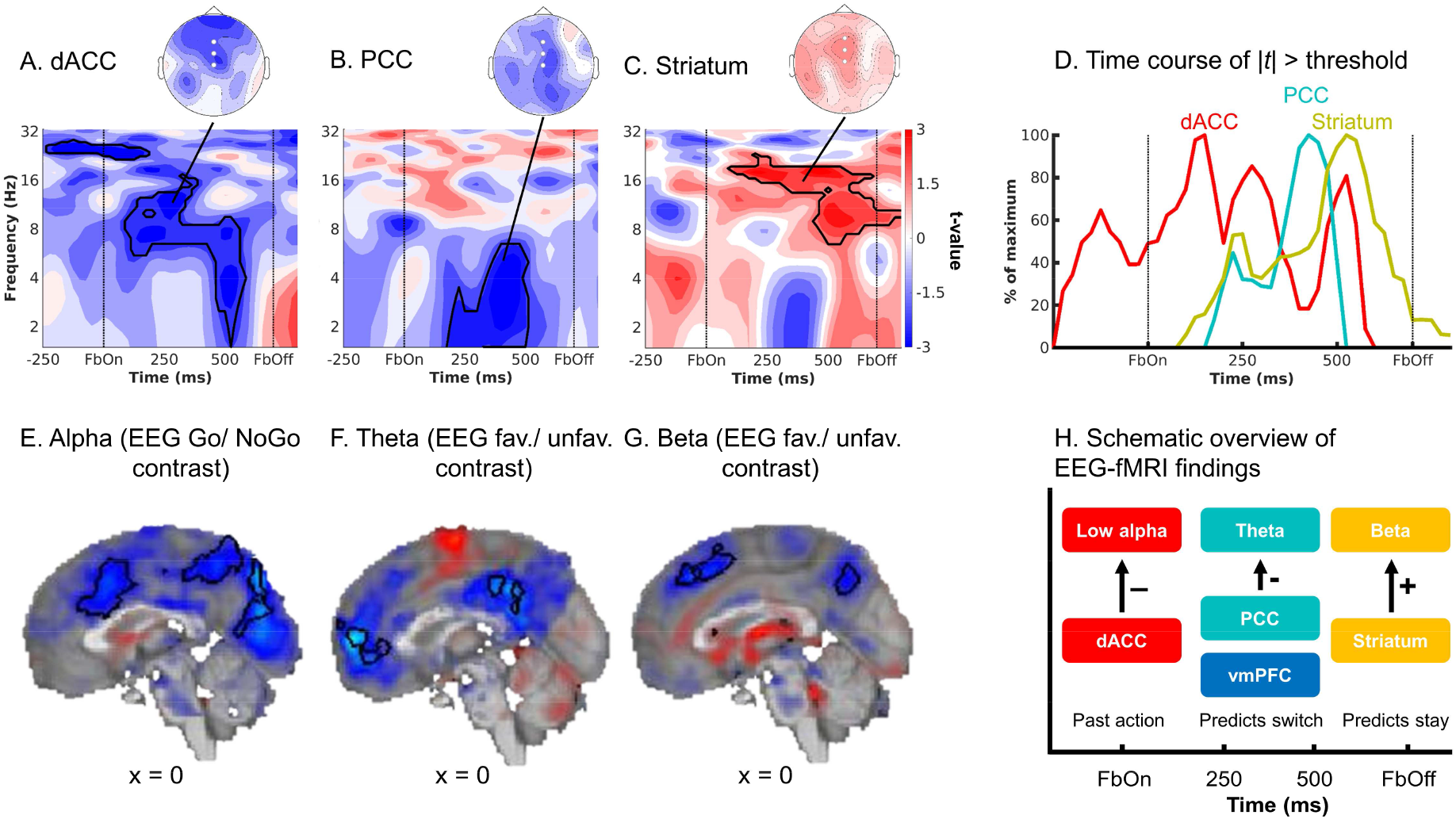
fMRI-informed EEG analyses. Unique temporal contributions of BOLD signal in (**A**) dACC, (**B**) PCC, and (**C**) striatum to average EEG power over midfrontal electrodes (Fz/ FCz/ Cz). Group-level *t*-maps display the modulation of the EEG power by trial-by-trial BOLD signal in the selected ROIs. dACC BOLD correlated negatively with early alpha/ theta power, PCC BOLD negatively with theta/ delta power, and striatal BOLD positively with beta/ alpha power. Areas surrounded by a black edge indicate clusters of |*t*| > 2 with *p* < .05 (cluster-corrected). Topoplots indicate the topography of the respective cluster. **D**. Time course of dACC, PCC, and striatal BOLD correlations, normalized to the peak of the time course of each region. dACC-lower alpha band correlations emerged first, followed by (negative) PCC-theta correlations and finally positive striatum-beta correlations. The reverse approach using lower alpha (**E**), theta (**F**) and beta (**G**) power as trial-by-trial regressors in fMRI GLMs corroborated the fMRI-informed EEG analyses: Lower alpha band power correlated negatively with the dACC BOLD, theta power negatively with vmPFC and PCC BOLD, and beta power positively with striatal BOLD. **H.** Schematic overview of the main EEG-fMRI results: dACC encoded the previously performed response and correlated with early midfrontal lower alpha band power. vmPFC/ PCC (correlated with theta power) and striatum (correlated with beta power) both encoded outcome valence, but had opposite effects on subsequent behavior. Note that activity in these regions temporally overlaps; boxes are ordered in temporal precedence of peak activity.

**Figure 6.**
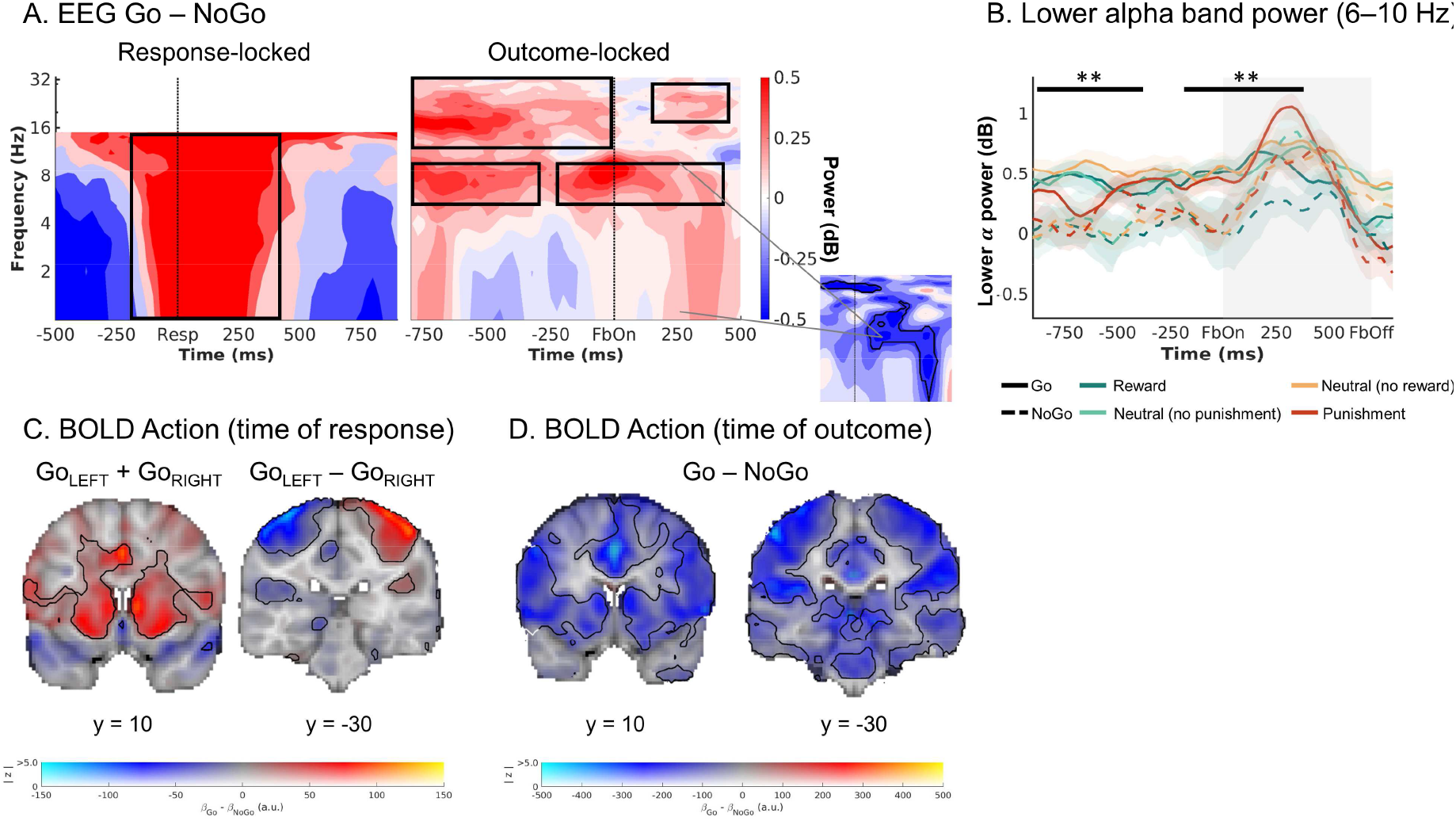
Exploratory follow-up analyses on dACC BOLD signal and midfrontal lower alpha band power. **A.** Midfrontal time-frequency response-locked (left panel) and outcome-locked (right panel). Before and shortly after outcome onset, power in the lower alpha band was higher on trials with Go actions than on trials with NoGo actions. The shape of this difference resembles the shape of dACC BOLD-EEG TF correlations (small plot; note that this plot depicts BOLD-EEG correlations, which were negative). Note that differences between Go and NoGo trials occurred already before outcome onset in the alpha and beta range, reminiscent of delay activity, but were not fully sustained throughout the delay between response and outcome. **B.** Midfrontal power in the lower alpha band per action x outcome condition. Lower alpha band power was consistently higher on trials with Go actions than on trials with NoGo actions, starting already before outcome onset. **C.** BOLD signal differences between Go and NoGo actions (activation by either left or right Go actions compared to the implicit baseline in the GLM, which contains the NoGo actions; left panel) and left vs. right hand responses (right panel) at the time or responses. Response-locked dACC BOLD signal was significantly higher for Go than NoGo actions. **D.** BOLD signal differences between Go and NoGo actions at the time of outcomes. Outcome-locked dACC BOLD signal (and BOLD signal in other parts of cortex) was significantly lower on trials with Go than on trials with NoGo actions.

Second, while pgACC BOLD did not correlate significantly with midfrontal EEG power (p = .184), BOLD in PCC was negatively correlated with theta/ delta power (Fig. 5B; 175–500 ms, 1–6 Hz, *p* = .014). This finding bore resemblance in terms of time-frequency space to the cluster of (negative) PE valence encoding in the theta band and (positive) PE magnitude encoding in the delta band identified in the EEG-only analyses (Fig. 4A). Complementary to the fMRI-informed EEG analyses, we also performed independent EEG-informed fMRI analyses, which showed the robustness of this EEG-fMRI correlation. We used the trial-by-trial EEG signal in the cluster identified in the EEG-only analyses (see Fig. 4 A, B) to predict BOLD signal across the brain (see Supplementary Fig. 18 for a graphical illustration of this approach). The EEG time-frequency-mask used to create the EEG regressor was defined based on the EEG-only analyses (Fig. 4A, B) and thus blind to the result of the fMRI-informed EEG analysis. We observed significant clusters of negative EEG-BOLD correlation in vmPFC and PCC (Fig. 5F; Supplementary Table 7). We thus discuss vmPFC and PCC together in the following.

Third, there was a significant positive correlation between striatal BOLD and midfrontal beta/ alpha power (driven by a cluster at 100–800 ms, 7–23 Hz, *p* = .010; Fig. 5C). This finding bore resemblance in time-frequency space to the cluster of positive PE valence encoding in beta power identified in the EEG-only analyses (Fig. 4A, C), but extended into the alpha range. Again, to support the robustness of this finding, we used trial-by-trial midfrontal beta power in the cluster identified in the EEG-only analyses (see Fig. 4A, C) to predict BOLD signal across the brain. Clusters of positive EEG-BOLD correlations in right dorsal caudate (and left parahippocampal gyrus) as well as clusters of negative correlations in bilateral dorsolateral PFC (dlPFC) and supramarginal gyrus (SMG; Fig. 5G; Supplementary Table 7) confirmed the positive striatal BOLD-beta power association. Given that the striatum is far away from the scalp and thus unlikely to be the source of midfrontal beta power over the scalp, and given the assumption that trial-by-trial variation in an oscillatory signal should correlate with BOLD signal in its source^39,40^, we speculate that dlPFC and SMG (identified in the EEG-informed fMRI analyses) are the sources of beta power over the scalp and act as an “antenna” for striatal signals. In line with this idea, previous studies have localized feedback-related beta power in lateral frontal and parietal regions, both using simultaneous EEG-fMRI^41–43^ and source-localization ^44,45^.

Finally, regarding the other three regions that showed a significant BOLD signature of biased PEs, BOLD in left motor cortex was significantly negatively correlated with midfrontal beta power (*p* = .002; around 0–625 ms; Supplementary Note 13 and Supplementary Fig. 19). There were no significant correlations between midfrontal EEG power and left inferior temporal gyrus or primary visual cortex BOLD (Supplementary Fig. 19). All results were robust to different analysis approaches including shorter trial windows, different GLM specifications, inclusion of task-condition and fMRI motion realignment regressors, and individual modelling of each region. TF results were not reducible to phenomena in the time domain (Supplementary Note 14 and Supplementary Fig. 20).

In sum, there were negative correlations between dACC BOLD and midfrontal lower alpha band power early after outcome onset, negative correlations between PCC BOLD and midfrontal theta/ delta power at intermediate time points, and positive correlations between striatal BOLD and midfrontal beta power at late time points. This temporal dissociation was especially clear in the time courses of the test statistics for each region (thresholded at |t| > 2 and summed across frequencies), for which the peaks of the cortical regions preceded the peak of the striatum (Fig. 5D, H). Note that time-frequency power is estimated over temporally extended windows (400 ms in our case), which renders any interpretation of the “onset” or “offset” of such correlations more difficult. In sum, these results are consistent with an “external model” of motivational biases arising from early cortical processes biasing later learning processes in the striatum.

### dACC BOLD and midfrontal lower alpha band power encode the previously performed action during outcome presentation

While the clusters of EEG-fMRI correlation in the theta/ delta and beta range matched the clusters identified in EEG-only analyses, the cluster of negative correlations between dACC BOLD and early midfrontal lower alpha band power was novel and did not match our expectations. Given that these correlations arose very soon after outcome onset, we hypothesized that dACC BOLD and midfrontal lower alpha band power might reflect a process occurring even before outcome onset, such as the maintenance (“memory trace”) of the previously performed response to which credit may later be assigned. We therefore assessed whether information of the previous response was present in dACC BOLD and in the lower alpha band around the time of outcome onset.

First, we tested for BOLD correlates of the previous response at the time of *outcomes* (eight outcome-locked regressors for every Go/ NoGo x reward/ no reward/ no punishment/ punishment combination) while controlling for motor-related signals at the time of the *response* (response-locked regressors for left-hand and right-hand button presses). At the time of outcomes, there was higher BOLD signal for NoGo than Go responses across several cortical and subcortical regions, peaking in both the dACC and striatum (Fig. 6D). This inversion of effects—higher BOLD for Go than NoGo responses at the time of response (see quality checks), but the reverse at the time of outcome—was also observed in the upsampled raw BOLD and was independent of the response of the next trial (Supplementary Note 15 and Supplementary Fig. 21). In sum, large parts of cortex, including the dACC, encoded the previously performed response at the moment outcomes were presented, in line with the idea that the dACC maintains a “memory trace” of the previously performed response.

Second, we tested for differences between Go and NoGo responses at the time of outcomes in midfrontal broadband EEG power. Power was significantly higher on trials with Go than on trials with NoGo responses, driven by clusters in the lower alpha band (spreading into the theta band; around 0.000–0.425 sec., 1–11 Hz, *p* = .012) and in the beta band (around 0.200–0.450 sec., 18–27 Hz, *p* = .022; Fig. 6A, B). The first cluster matched the time-frequency pattern of dACC BOLD-alpha power correlations (Fig. 5A).

If this activity cluster contained a signature of the previously performed response, it might have been present throughout the delay between cue offset and outcome onset. When repeating the above permutation test including the last second before outcome onset, there were significant differences again, driven by a sustained cluster in the beta band (−1–0 sec., 13–33 Hz, *p* = .002) and two clusters in the alpha/ theta band (Cluster 1: −1.000– −0.275 sec., 1–10 Hz, *p* = 0.014; Cluster 2: −0.225–0.425 sec., 1– 11 Hz, *p* = .022; Fig. 6B). These findings suggest that lower alpha band power might reflect a sustained memory of the previously performed response. Additional analyses (Supplementary Note 15 and Supplementary Fig. 21) yielded that this Go-NoGo trace during outcome processing did not change over the time course of the experiment, suggesting that it did not reflect typical fatigue/ time-on task effects often observed in the alpha band.

Again, we performed the reverse EEG-fMRI analysis using trial-by-trial power in the identified lower alpha band cluster (Fig. 6B) as an additional regressor in the quality-check fMRI GLM. Clusters of negative EEG-BOLD occurred correlation in a range of cortical regions, including dACC and precuneus (Fig. 5E; Supplementary Table 7). In sum, both dACC BOLD signal and midfrontal lower alpha band power contained information about the previously performed response, consistent with the idea that both signals reflect a “memory trace” of the response to which credit is assigned once an outcome is obtained.

### Striatal and vmPFC/ PCC BOLD differentially relate to action policy updating

EEG correlates of PCC BOLD and striatal BOLD occurred later than for the dACC BOLD and overlapped with classical feedback-related midfrontal theta and beta power responses. We hypothesized that those neural signals might be more closely related to the updating of action policies (i.e., which action to perform for each cue) and predict the next response to the same cue^30,46^. We thus used the trial-by-trial BOLD responses in dACC, PCC/ vmPFC, and striatum to predict whether participants would repeat the same response on the next trial with the same cue (“stay”) or switch to another response (“shift”). Mixed-effects logistic regression yielded that dACC BOLD did not significantly predict response repetition (*b* = −0.019, *SE* = 0.016, χ^2^(1) = 1.294, *p* = .255). In contrast, BOLD in PCC/ vmPFC and striatum did predict response repetition, though in opposite directions: Participants were significantly more likely to *repeat* the same response when striatal BOLD was high (*b* = 0.067, *SE* = 0.024, χ^2^(1) = 9.051, *p* = .003), but more likely to *switch* to another response when vmPFC BOLD (*b* = −0.065, *SE* = 0.020, χ^2^(1) = 8.765, *p* = .003) or PCC BOLD (*b* = −0.036, *SE* = 0.016, χ^2^(1) = 3.691, *p* = .030; Fig. 5H) was high (Supplementary Fig. 22). Similarly, high pgACC BOLD predicted a higher likelihood of switching, likening it with the circuits formed by vmPFC and PCC (*b* = −0.076, *SE* = 0.017, χ^2^(1) = 15.559, *p* < .001). We also inspected the raw upsampled HRF shapes per region per condition, confirming that differential relationships were not driven by differences in HRF shapes across regions.

We also tested whether trial-by-trial midfrontal lower alpha band, theta, or beta power (within the clusters identified in the EEG-only analyses) predicted action policy updating. Participants were significantly more likely to repeat the same response when beta power was high (*b* = 0.145, *SE* = 0.041, χ^2^(1) = 11.886, *p* < .001), but more likely to switch when theta power was high (*b* = −0.099, *SE* = 0.047, χ^2^(1) = 4.179, *p* = .041). Notably, unlike its BOLD correlate in ACC, lower alpha band power did predict response repetition, with more repetition when alpha power was high (*b* = .0.179, SE = 0.052, χ^2^(1) = 10.711, *p* = .001; Supplementary Fig. 22).

In sum, high striatal BOLD and midfrontal beta power predicted that the same response would be repeated on the next encounter of a cue, while high vmPFC and PCC BOLD and high theta power predicted that participants would switch to another response. Thus, although both striatal and vmPFC/ PCC BOLD positively encoded biased prediction errors, these two sets of regions had opposite roles in learning: while the striatum reinforced previous responses, vmPFC/ PCC triggered the shift to another response strategy (Fig. 5H).

## Discussion

We investigated neural correlates of biased learning for Go and NoGo responses. In line with previous research^3,9^, participants’ behavior was best described by a computational model featuring faster learning from rewarded Go responses and slower learning from punished NoGo responses. Neural correlates of biased PEs were present in BOLD signals in several regions, including ACC, PCC, and striatum. These regions exhibited distinct midfrontal EEG power correlates. Most importantly, correlates of prefrontal cortical BOLD preceded correlates of striatal BOLD: Trial-by-trial dACC BOLD correlated with lower alpha band power immediately after outcome onset, followed by PCC (and vmPFC) BOLD correlated with theta power, and finally, striatal BOLD correlated with beta power. These results suggest that the architecture of the asymmetric striatal pathways might not be the only neural structure that gives rise to motivational learning biases; instead, the PFC might critically contribute to these biases.

The observation that both PFC and striatal BOLD signal reflected biased PEs might be explained by three different models. One model assumes that both PFC and striatal processes arrive at biased learning independently of each other, which is highly unlikely given strong recurrent connections between both regions^18,19,47^. Another model incorporates such interconnections, but assumes that striatum leads the PFC. While such a model is in line with past animal studies^48^ and modeling work^49^, it would predict EEG correlates of the PFC to trail after EEG correlates of the striatum—or at least to occur with considerable delay after outcome onset. This model is not supported by our findings, which showed EEG correlates of PFC regions soon after outcome onset, preceding striatal EEG correlates. These early EEG correlates of PFC BOLD are in line with single cell recordings in PFC which show responses confined to the first 500 ms following outcome onset^50,51^, corroborating that PFC outcome processing occurs before the time of EEG correlates of striatal BOLD. The only model consistent with our data assumes recurrent connections between PFC and striatum, but with the PFC leading the striatum. Hence, these results are in line with a model of PFC biasing striatal outcome processing, giving rise to motivational learning biases in behavior.

The dominant idea about the origin of motivational biases has been that these biases are an emergent feature of the asymmetric direct/ indirect pathway architecture in the basal ganglia ^2,19^. We find that these biases are present first in prefrontal cortical areas, notably dACC and PCC, which argues against biases being purely driven by subcortical circuits. Rather, motivational learning biases might be an instance of sophisticated, even “model-based” learning processes in the striatum instructed by the prefrontal cortex^52,53^. An influence of PFC on striatal RL has prominently been observed in the case of model-based vs. model-free learning^23,24^ and has been stipulated as a mechanism of how instructions can impact RL^20,21^. Although there are reports of striatal processes preceding prefrontal processes within learning tasks^48,54^, the opposite pattern of PFC preceding striatum has been observed as well^55^ and a causal impact of PFC on striatal learning is well established^56,57^. In particular, we have previously observed that motivational biases in action selection might arise from early prefrontal inputs to the striatum, as well^17^. Prefrontal influences on striatal processes might thus be a common signature of both motivational response and learning biases.

The particular subregion of PFC showing the earliest EEG correlates was the dACC. This observation is in line with an earlier EEG-fMRI study reporting dACC to be part of an early valuation system preceding a later system comprising vmPFC and striatum^58^. The dACC has been suggested to encode models of agents’ environment^59,60^ that are relevant for interpreting outcomes, with BOLD in this region scaling with the size of PEs^25,26^ and indexing how much should be learned from new outcomes. We hypothesize that, at the moment of outcome, dACC maintains a “memory trace” of the previously performed response^61^ which might modulate the processing of outcomes as soon as they become available^62,63^. Notably, dACC exhibited stronger BOLD signal for Go than NoGo responses at the time of participants’ response, but this pattern reversed at the time of outcomes. This reversal rules out the possibility that response-locked BOLD signal simply spilled over into the time of outcomes. Future research will be necessary to corroborate such a motor “memory trace” in dACC. In sum, the dACC might be in a designated position to inform subsequent outcome processing in downstream regions by modulating the learning rate as a function of the previously performed response and the obtained outcome. Rather than striatal circuits being sufficient for the emergence of motivational biases, the more “flexible” PFC seems to play an important role in instructing downstream striatal learning processes.

Striatal, dACC and PCC BOLD encoded biased PEs. In line with previous research, striatal BOLD positively linked to midfrontal beta power^41,42^, which positively encoded PE valence^28,34,64^, with correlations extending into alpha power. PCC and vmPFC BOLD negatively linked to midfrontal theta/ delta power^17,65,66^, which encoded PE valence negatively, but PE magnitude positively. Notably, theta/ delta power correlates of vmPFC/ PCC BOLD preceded beta power correlates of striatal BOLD in time, which aligns with previous findings of motivational response biases being first visible in the vmPFC BOLD before they impact striatal action selection^17^. Notably, EEG correlates of striatal BOLD during outcome processing were in the beta band—in contrast to previously observed correlates of striatal BOLD during action selection in the theta band^17^. This dissociation suggests important differences in the role of the striatum in these two processes. The frequency-specific nature of these EEG-fMRI correlations further suggests that they are signatures of task-induced events that are specific to the trial phase and unlikely to reflect general anatomical connectivity. In sum, while these EEG-fMRI findings on outcome processing resemble our previous EEG-fMRI findings on action selection in that prefrontal signals precede striatal signals, they are dissociated in terms of the frequency specificity, highlighting the distinct roles of the striatum in these processes.

Positive encoding of prediction errors in striatal BOLD signal is a well-established phenomenon^38,67^. Striatal BOLD was better described by biased PEs than by standard PEs, corroborating the presence of motivational learning biases also in striatal learning processes. Notably, EEG correlates of striatal BOLD peaked rather late, suggesting that these processes are informed by early sources in PFC which are connected to the striatum via recurrent feedback loops^18,47^. Positive prediction errors increase the value of a performed action and thus strengthen action policies. Hence, it is not surprising that high striatal BOLD signal and midfrontal beta power predicted action repetition^68,69^.

In contrast to striatal learning signals, the PCC and vmPFC BOLD as well as midfrontal theta and delta power signals were more complicated: Theta encoded PE valence, delta encoded PE magnitude. Both correlates showed opposite polarities. This observation is in line with previous literature suggesting that midfrontal theta and delta power might reflect the “saliency” or “surprise” aspect of PEs^31,32,70^. Surprises have the potential to disrupt an ongoing action policy^71^ and motivate a shift to another policy, which might explain why these signals predicted switching to another response^72,73^. Notably, this EEG surprise signal was only significantly correlated with the biased (but not the standard) PE term, corroborating that the surprise attributed to outcomes depends on the previously performed response in line with motivational learning biases. In sum, both vmPFC and striatum encode biased PEs, though with different consequences for future action policies.

Taken together, distinct brain regions processed outcomes in a biased fashion at distinct time points with distinct EEG power correlates. Simultaneous EEG-fMRI recordings allowed us to infer when those regions reached their peak activity^74^. However, the correlational nature of BOLD-EEG links precludes strong statements about these regions actually generating the respective power phenomena. Alternatively, activity in those regions might merely modulate the amplitude of time-frequency responses originating from other sources. Furthermore, while the observed associations align with previous literature ^17,41,42,65,66^, the considerable distance of the striatum to the scalp raises the question whether scalp EEG could in principle reflect striatal activity, at all^75,76^. Intracranial recordings have observed beta oscillations during outcome processing in the striatum before^69,77–79^. Also, our analysis controlled for BOLD signal in motor cortex, an alternative candidate source for beta power, suggesting that late midfrontal beta power did not merely reflect motor cortex beta. Even if the striatum is not the generator of the beta oscillations over the scalp, their true (cortical) generator might be tightly coupled to the striatum and thus act as a “transmitter” of striatal beta oscillations. In fact, the analyses using trial-by-trial beta power to predict BOLD yielded significant clusters in dlPFC and SMG, two candidate regions for such a “transmitter”.

We observed EEG correlates of striatal BOLD at a rather late time point after outcome onset. While we conclude that biased outcome processing occurs much earlier in cortical regions than the striatum, it is possible that the modulating influence of the striatum on cortical sources of beta synchronization over the scalp (possibly dlPFC and SMG, corroborating previous EEG-fMRI^41–43^ and source-reconstruction findings^44,45^) takes time to surface. However, speaking against any delay, some single studies have reported maximal correlations between striatal LFPs and scalp EEG at a time lag of 0^80^. Regardless, even in the presence of a non-zero lag, our main conclusion would hold: Biased learning is present in cortical regions early after outcome onset, which cannot be a consequence of striatal input, but must constitute an independent origin of motivational learning biases.

In order to make inferences about the relative order of PE processing in different brain regions, we must assume that the regressor in our EEG-fMRI analysis approach—the trial-by-trial BOLD amplitude in selected regions—mostly reflects the PE signal rather than learning-unrelated processes occurring in parallel. In support of this assumption, animal recordings have indeed found that neural activity in ACC, PCC, and striatum is dominated by reward processing during outcome receipt^81–85^ and meta-analyses on human BOLD signal have found strong effect sizes for PE processing in these regions^38,67^. Importantly, we observe transient EEG-fMRI correlations that are likely event-related rather than reflecting resting-state like correlations. We thus favor the conclusion that the observed EEG-fMRI correlations reflect differences in the timing of PE processing in these regions, although we cannot fully exclude the possibility that parallel processes unrelated to (biased) learning contribute to these correlations. Note that, while outcome processing in these regions is better described by biased than by standard PEs, each region might encode PEs in an idiosyncratic way (potentially reflecting noise in the value representations^86^) and these residual idiosyncrasies drive the EEG-fMRI correlations even when controlling for biased PEs predicted by the winning computational model.

The correlational nature of the study prevents strong statements over any causal interactions between the observed regions. We assume here that a region showing an earlier midfrontal EEG correlate influences other regions showing later midfrontal EEG correlates, and such an influence is plausible given findings of feedback loops between prefrontal regions and the striatum^47^. Future studies targeting those regions via selective causal manipulations will be necessary to test for the causal role of PFC in informing striatal learning. Furthermore, while parameter recovery for most parameters in the winning computational model (including the effective learning rates incorporating the learning bias) was excellent, parameter recovery for the learning bias term itself was positive, but weaker (see Supplementary Note 6). Supplementary models tested incorporating a perseveration parameter (see Supplementary Note 8) yielded higher model recovery, but failed to capture crucial aspects of the biased learning under investigation. Future studies comprising larger samples of participants should explore alternative implementations to reliably quantify individual differences in these learning biases.

In conclusion, biased learning—increased credit assignment to rewarded action, decreased credit assignment to punished inaction—was visible both in behavior and in BOLD signal in a range of regions. EEG correlates of prefrontal cortical regions, notably dACC and PCC, *preceded* correlates of the striatum, consistent with a model of the PFC biasing RL in the striatum. The dACC appeared to hold a “motor memory trace” of the past response, biasing early outcome processing. Subsequently, biased learning was also present in vmPFC/ PCC and striatum, with opposite roles in adjusting vs. maintaining action policies. These results refine previous views on the neural origin of these learning biases, suggesting they might not only rely on subcortical parts of the brain typically associated with rigid, habit-like responding, but rather incorporate frontal inputs that are associated with counterfactual reasoning and increased behavioral flexibility^87,88^. The PFC is typically believed to facilitate goal-directed over instinctive processes. Hence, PFC involvement into biased learning suggests that these biases are not necessarily agents’ inescapable “fate”, but rather likely act as global “priors” that facilitate learning of more local relationships. They allow for combining “the best of both worlds”—long-term experience with consequences of actions and inactions together with flexible learning from rewards and punishments.

## Materials and methods

### Participants

Thirty-six participants (*M_age_* = 23.6, SD_age_ = 3.4, range 19–32; 25 women; all right-handed; all normal or corrected-to-normal vision) took part in a single 3-h data collection session, for which they received €30 flat fee plus a performance-dependent bonus (range €0–5, *M_bonus_* = €1.28, *SD*_bonus_ = 1.54). The study was approved by the local ethics committee (CMO2014/288; Commissie Mensengeboden Onderzoek Arnhem-Nijmegen) and all participants provided written informed consent. Exclusion criteria comprised claustrophobia, allergy to gels used for EEG electrode application, hearing aids, impaired vision, colorblindness, history of neurological or psychiatric diseases (including heavy concussions and brain surgery), epilepsy and metal parts in the body, or heart problems. Sample size was based on previous EEG studies with a comparable paradigm^9,89^.

Behavioral and modeling results include all 36 participants. The following participants were excluded from analyses of neural data: For two participants, fMRI functional-to-standard image registration failed; hence, all fMRI-only results are based on 34 participants (*M_age_* = 23.47, 25 women). Four participants exhibited excessive residual noise in their EEG data (> 33% rejected trials) and were thus excluded from all EEG analyses; hence, all EEG-only analyses are based on 32 participants (*M_age_* = 23.09, 23 women). For combined EEG-fMRI analyses, we excluded the above-mentioned six participants plus one more participant whose regression weights for every regressor were about ten times larger than for other participants, leaving 29 participants (*M_age_* = 23.00, 22 women). Exclusions were in line with a previous analysis of this data set^17^. fMRI- and EEG-only results held when analyzing only those 29 participants (see Supplementary Notes 1–5 and Supplementary Figures 1–4).

### Task

Participants performed a motivational Go/ NoGo learning task^3,9^ administered via MATLAB 2014b (MathWorks, Natick, MA, United States) and Psychtoolbox-3.0.13. On each trial, participants saw a gem-shaped cue for 1300 ms which signaled whether they could potentially win a reward (Win cues) or avoid a punishment (Avoid cues) and whether they had to perform a Go (Go cue) or NoGo response (NoGo cue). They could press a left (Go_LEFT_), right (Go_RIGHT_), or no (NoGo) button while the cue was presented. Only one response option was correct per cue. Participants had to learn both cue valence and required action from trial-and-error. After a variable inter-stimulus-interval of 1,400–1,600 ms, the outcome was presented for 750 ms. Potential outcomes were a reward (symbolized by coins falling into a can) or neutral outcome (can without money) for Win cues, and a neutral outcome or punishment (symbolized by money falling out of a can) for Avoid cues. Feedback validity was 80%, i.e., correct responses were followed by positive outcomes (rewards/ no punishments) on only 80% of trials, while incorrect responses were still followed by positive outcomes on 20% of trials. Trials ended with a jittered inter-trial interval of 1250–2000 ms, yielding total trial lengths of 4700–6650 ms.

Participants gave left and right Go responses via two button boxes positioned lateral to their body. Each box featured four buttons, but only one button per box was required in this task. When participants accidentally pressed a non-instructed button, they received the message “Please press one of the correct keys” instead of an outcome. In the analyses, these responses were recoded into the instructed button on the respective button box. In the fMRI GLMs, such trials were modeled with a separate regressor.

Before the task, participants were instructed that each cue could be followed by either reward or punishment, that each cue had one optimal response, that feedback was probabilistic, and that the rewards and punishments were converted into a monetary bonus upon completion of the study. They performed an elaborate practice session in which they got familiarized first with each condition separately (using practice stimuli) and finally practiced all conditions together. They then performed 640 trials of the main task, separated into two sessions of 320 trials with separate cue sets. Introducing a new set of cues allowed us to prevent ceiling effects in performance and investigate continuous learning throughout the task. Each session featured eight cues that were presented 40 times. After every 100–110 trials (∼ 6 min.), participants could take a self-paced break. The assignment of the gems to cue conditions was counterbalanced across participants, and trial order was pseudo-random (preventing that the same cue occurred on more than two consecutive trials).

### Behavior analyses

We used mixed-effects logistic regression (as implemented in the R package *lme4*) to analyze behavioral responses (Go vs. NoGo) as a function of required action (Go/ NoGo), cue valence (Win/ Avoid), and their interaction. We included a random intercept and all possible random slopes and correlations per participant to achieve a maximal random-effects structure^90^. Sum-to-zero coding was employed for the factors. Type 3 *p*-values were based on likelihood ratio tests (implemented in the R package *afex*). We used a significance criterion of α = .05 for all the analyses.

Furthermore, we used mixed-effects logistic regression to analyze “stay behavior”, i.e., whether participants repeated an action on the next encounter of the same cue, as a function of outcome valence (positive: reward or no punishment/ negative: no reward or punishment), outcome salience (salient: reward or punishment/ neutral: no reward or no punishment), and performed action (Go/ NoGo). We again included all possible random intercepts, slopes, and correlations.

### Computational modeling

We fit a series of increasingly complex RL models to participants’ choices to decide between different algorithmic explanations for the emergence of motivational biases in behavior. We employed the same set of nested models as in previous studies using this task^3,9^. For tests of alternative biases specifications, see Supplementary Notes 7–9 and Supplementary Fig. 6–8.

#### Model space

To determine whether a Pavlovian response bias, a learning bias, or both biases jointly predicted behavior best, we fitted a series of increasing complex computational models. In each trial (t), choice probabilities for all three response options (a) given the displayed cue (s) were computed from their action weights (modified Q-values) using a softmax function:

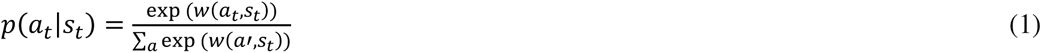

After each response, action values were updated with the prediction error based on the obtained outcome *r* ∈ {−1; 0; 1}. As the starting model (M1), we fitted an standard delta-learning model ^91^ in which action values were updated with prediction errors, i.e., the deviation between the experienced outcome and expected outcome. This model contained two free parameters: the learning rate (ε) scaling the updating term and the feedback sensitivity (ρ) scaling the received outcome (i.e., higher feedback sensitivity led to choices more strongly guided by value difference, akin to the role of the inverse temperature parameter frequency used in reinforcement learning models):

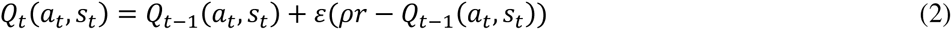

In this model, choice probabilities were fully determined by action values, without any bias. We initialized action values Q_0_ such that they reflected a “neutral” expected value for each action. Win cues could lead to reward (+1) or neutral (0) outcomes and Avoid cues to neutral (0) or punishment (−1) outcomes. A neutral expected value would assign equal probability to either possible outcome, leading to expectations of +1/2 and −1/2, respectively. In addition, because participants’ feedback sensitivity parameter ρ reflected how participants weighed the outcomes they received, also the initial values had to be multiplied with the feedback sensitivity to stay neutral between 0 and participants’ re-weighted positive/ negative outcome of +/−1*ρ. Thus, initial action values Q_0_ were set to 1/2*ρ (Win cues) and - 1/2*ρ (Avoid cues).

Unlike previous versions of the task^3^, cue valences were not instructed, but had to be learned from outcomes, as well^9^. Thus, until experiencing the first non-neutral outcome (reward or punishment) for a cue, participants could not know its valence and thus not learn from neutral feedback. Hence, for these early trials, action values were multiplied with zero when computing choice probabilities ^9^. After the first encounter of a valenced outcome, action values were “unmuted” and started to influence choices probabilities, retrospectively considering all previous outcomes^9^.

In M2, we added the Go bias parameter *b*, which accounted for individual differences in participants’ overall propensity to make Go responses, to the action values Q, resulting in action weights w:

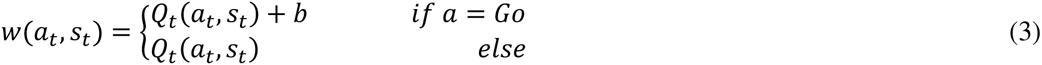

In M3, we added a Pavlovian response bias π, scaling how positive/ negative cue valence (Pavlovian values) increased/ decreased the weights of Go responses:

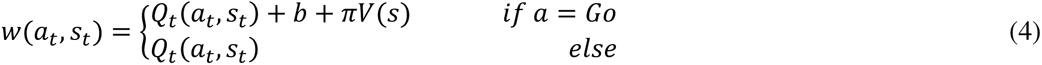

Participants were instructed that a cue was either a Win cue (affording rewards or neutral outcomes) or an Avoid cue (affording neutral outcomes or punishments). Hence, cue valence (Win/ Avoid) did not have to be learned instrumentally; instead, it could be inferred as soon participants experienced a non-neutral outcome. Until that moment, cue valence *V(s)* was set to zero. Afterwards, *V(s)* was set to +0.5 for Win cues and −0.5 for Avoid cues. Note that choosing different values than 0.5 would merely rescale the bias parameter π (e.g., halving π with cue valences of +1 and −1) without any changes in the model’s predictions. The Pavlovian response bias affected left-hand and right-hand Go responses similarly and thus reflected generalized activation/ inactivation by the cue valence.

In M4, we added a learning bias κ, increasing the learning rate for rewards after Go responses and decreasing it for punishments after NoGo responses. The learning bias was specific to the response shown, thus reflecting a specific enhancement in action learning/ impairment in unlearning for that particular response. Conceptually, learning rates differed between response-outcome conditions in the following way:

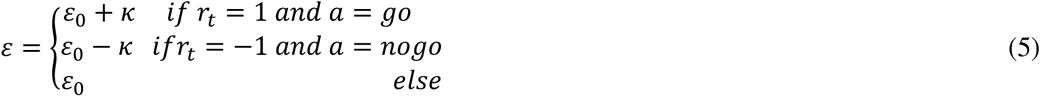

In the technical implementation of this model, learning rates were sampled in continuous space and then inverse-logit transformed to constrain them to the range [0 1]^3,9^. However, after this transformation, the impact of adding vs. subtracting the learning bias κ would no longer be symmetric. Hence, for baseline learning rates ε_0_ < 0.5, we first computed the difference between the baseline learning rate and the learning rates for punished NoGo responses and used this difference to compute the learning rate for rewarded Go responses:

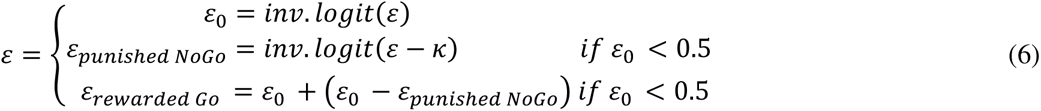

Notably, this procedure is only guaranteed to work when ε_0_ < 0.5. For ε_0_ > 0.5, the difference term could become > 0.5 and the learning rate for rewarded Go responses would become > 1, which is impractical. Hence, for ε_0_ > 0.5, we first computed the learning rate for rewarded Go responses and used the difference to the baseline learning rate ε_0_ to compute the learning rate for punished NoGo responses:

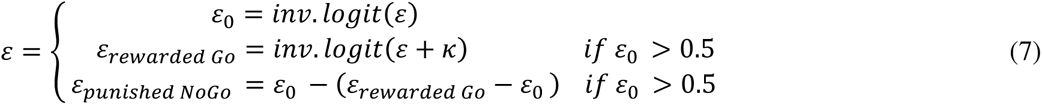

In the model M5, we included both the Pavlovian response bias and the learning bias.

The weakly informative hyperpriors were set to *X_ρ_∼N(2,3)*, *X_ε_∼N(0,2)*, *X_b,_*_π_*_,κ_∼N(0,3)*, in line with previous implementations of this model^3,9^. The same priors (for the same parameters) were used across different model implementations to not bias model comparison. Alternative hyperpriors did not change the results. For computing the participant-level parameters, ρ was exponentiated to constrain it to positive values, and the inverse-logit transformation was applied to ε.

#### Model fitting and comparison

For model fitting and comparison, we used hierarchical Bayesian inference as implemented in the CBM toolbox in MATLAB^92^. This approach combines hierarchical Bayesian parameter estimation with random-effects model comparison^93^. The fitting procedure involves two steps, starting with the Laplace approximation of the model evidence to compute the group evidence, which quantifies how well each model fits the data while penalizing for model complexity. Both group-level and individual-level parameters are estimated using an iterative algorithm. We used wide Gaussian priors (see hyperpriors above) and exponential and sigmoid transforms to constrain parameter spaces. Subsequent random-effects model selection allows for the possibility that different models generated the data for different participants. Participants contribute to the group-level parameter estimation in proportion to how well a given model fits their data, quantified via a responsibility measure (i.e., the probability that the model at hand is responsible for generating data of the respective participant). This model-comparison approach has been shown to be less susceptible to the influence of outliers^92^. We selected the “winning” model based on the protected exceedance probability.

#### Model validation

We assured that the winning model was able to reproduce the data, using the sampled combinations of participant-level parameter estimates to create 3600 agents that “played” the task. We employed two approaches to simulate the task: *posterior predictive model simulations* and *one-step-ahead model predictions*. In the posterior predictive model simulations, agents’ choices were sampled probabilistically based on their action values, and outcomes probabilistically sampled based on their choices. This method ignores participant-specific choice histories and can thus yield choice/ outcome sequences that diverge considerably from participants’ actual experiences. In contrast, one-step-ahead predictions use participants’ actual choices and experienced outcomes in each trial to update action values. We simulated choices for each participant using both methods, which confirmed that the winning model M5 (“asymmetric pathways model”) was able to qualitatively reproduce the data, while an alternative implementation of biased learning (“action priming model”) failed to do so (see Supplementary Note 7 and Supplementary Fig. 6).

### fMRI data acquisition

fMRI data were collected on a 3T Siemens Magnetom Prisma fit MRI scanner with a 64-channel head coil. During scanning, participants’ heads were restricted using foam pillows and strips of adhesive tape were applied to participants’ forehead to provide active motion feedback and minimize head movement ^94^. After two localizer scans to position slices, we collected functional scans with a whole-brain T2*-weighted sequence (68 axial-oblique slices, TR = 1400 ms, TE = 32 ms, voxel size 2.0 mm isotropic, interslice gap 0 mm, interleaved multiband slice acquisition with acceleration factor 4, FOV 210 mm, flip angle 75°, A/ P phase encoding direction). The first seven volumes of each run were automatically discarded. This sequence was chosen because of its balance between a short TR and relatively high spatial resolution, which was required to disentangle cue and outcome-related neural activity. Pilots using different sequences yielded that this sequence performed best in reducing signal loss in striatum.

Furthermore, after task completion, we removed the EEG cap and collected a high-resolution anatomical image using a T1-weighted MP-RAGE sequence (192 sagittal slices per slab, GRAPPA acceleration factor = 2, TI = 1100 ms, TR = 2300 ms, TE = 3.03 ms, FOV 256 mm, voxel size 1.0 mm isotropic, flip angle 8°) which was used to aid image registration, and a gradient fieldmap (GRE; TR = 614 ms, TE1 = 4.92 ms, voxel size 2.4 mm isotropic, flip angle 60°) for distortion correction. For one participant, no fieldmap was collected due to time constraints. At the end of each session, an additional DTI data collection took place; results will be reported elsewhere.

### fMRI preprocessing

All fMRI pre-processing was performed in FSL 6.0.0. After cleaning images from non-brain tissue (brain-extraction with BET), we performed motion correction (MC-FLIRT), spatial smoothing (FWHM 3 mm), and used fieldmaps for B0 unwarping and distortion correction in orbitofrontal areas. We used ICA-AROMA^95^ to automatically detect and reject independent components associated with head motion. Finally, images were high-pass filtered at 100 s and pre-whitened. After the first-level GLM analyses, we computed and applied co-registration of EPI images to high-resolution images (linearly with FLIRT using boundary-based registration) and to MNI152 2mm isotropic standard space (non-linearly with FNIRT using 12 DOF and 10 mm warp resolution).

### ROI selection

For fMRI-informed EEG analyses, we first created a functional mask as the conjunction of the PE_STD_ and PE_DIF_ contrasts by thresholding both z-maps at z > 3.1, binarizing, and multiplying them (see Supplementary Figures 9 and 10). After visual inspection of the respective clusters, we created seven anatomical masks based on the probabilistic Harvard-Oxford Atlas (thresholded at 10%): striatum and ACC (see above), vmPFC (combined frontal pole, frontal medial cortex, and paracingulate gyrus), motor cortex (combined precentral and postcentral gyrus), PCC (Cingulate Gyrus, posterior division), ITG (Inferior Temporal Gyrus, posterior division, and Inferior Temporal Gyrus, temporooccipital part) and primary visual cortex (Lingual Gyrus, Occipital Fusiform Gyrus, Occipital Pole). We then multiplied this functional mask with each of the seven anatomical masks, returning seven masks focused on the respective significant clusters, which were then used for signal extraction. For the dACC mask, we manually excluded voxels in pgACC belonging to a distinct cluster. Masks were back-transformed to each participant’s native space.

For bar plots in Fig. 3A, we multiplied the anatomical masks of vmPFC and striatum specified above with the binarized outcome valence contrast.

### fMRI analyses

For each participant, data were modelled using two event-related GLMs. First, we performed a model-based GLM in which we used trial-by-trial estimates of biased PEs as regressors. Second, we used another model-free GLM in which we modeled all possible action x outcome combinations via outcome-locked categorical regressors while at the same time modeling response-locked left- and right-hand response regressors. This model free GLM also contained the outcome valence contrast reported as an initial manipulation check.

In the model-based GLM, we used two model-based regressors that reflected the trial-by-trial prediction error (PE) update term. The update term was computed by multiplying the prediction-error with the condition-specific learning rate. As described above, in the winning model M5, the learning bias term κ leads to altered learning from “congruent” action-outcome pairs, with faster learning of Go actions followed by rewards, but slower unlearning of NoGo actions followed by punishments. To compute trial-by-trial updates, we extracted the group-level parameters of the best fitting computational model M5 (asymmetric pathways model) and used those parameters to compute the prediction error on every trial for every participant. Using the same parameter for each participant is warranted when testing for the same qualitative learning pattern across participants^96^. Given that both standard (base model M1) and biased (winning model M5) PEs were highly correlated (mean correlation of 0.921 across participants, range 0.884–0.952), it appeared difficult to distinguish standard learning from biased learning. As a remedy, we decomposed the biased PE into the standard PE plus a difference term as *PE_BIAS_ = PE_STD_ + PE_DIF_* ^22,36^. Any region displaying truly biased learning should significantly encode *both* the standard PE term and the difference term. The standard PE and difference term were much less correlated (mean correlation of −0.020, range −0.326–0.237). To control for cue-related activation, we furthermore added four regressors spanned by crossing cue valence and performed action (Go response to Win cue, Go response to Avoid cue, NoGo response to Win cue, NoGo response to Avoid cue).

The model-free GLM included a separate regressor for each of the eight conditions obtained when crossing performed action (Go/ NoGo) and obtained outcome (reward/ no reward/ no punishment/ punishment). We fitted four contrasts: 1) one contrast comparing conditions with positive (reward/ no punishment) and negative (no reward/ punishment) outcomes, used as a quality check to identify regions that encoded outcome valence; 2) one contrast comparing Go vs. NoGo responses at the time of the outcome; 3) one contrast summing of left- and right-hand responses, reflecting Go vs. NoGo responses at the time of the response; and 4) one contrast subtracting right-from left-handed responses, reflecting lateralized motor activation. As this GLM resulted in empty regressors for several participants when fitted on a block level, making it impossible to use the data of the respective blocks on a higher level, we instead concatenated blocks and performed a single GLM per participant. We therefore registered the data from all blocks to the middle image of the first block (default reference volume in FSL) using MCFLIRT. The first and last 20 seconds of each block did not feature any task-related events, such that carry-over effects of task events in the design matrix from one block to another were not possible.

In both GLMs, we added four regressors of no interest: one for the motor response (left = +1, right = −1, NoGo = 0), one for error trials, one for outcome onset, and one for trials with invalid motor response (and no outcome respectively). We also added nine or more nuisance regressors: the six realignment parameters from motion correction, mean cerebrospinal fluid (CSF) signal, mean out-of-brain (OBO) signal, and a separate spike regressor for each volume with a relative displacement of more than 2 mm (occurred in 10 participants; in those participants: M = 7.40, range 1–29). For the model-free GLM, nuisance regressors were added separately for each block as well as an overall intercept per block. We convolved task regressors with double-gamma haemodynamic response function (HRF) and high-pass filtered the design matrix at 100 s.

First-level contrasts were fit in native space. Afterwards, co-registration and reslicing was applied to participants’ contrast maps, which were then combined on a (participant and) group level using FSL’s mixed effects models tool FLAME with a cluster-forming threshold of z > 3.1 and cluster-level error control at α < .05 (i.e., two one-sided tests with α < .025).

### EEG data acquisition

We recorded EEG data with 64 channels (BrainCap-MR-3-0 64Ch-Standard; Easycap GmbH; Herrsching, Germany; international 10-20 layout, reference electrode at FCz) plus channels for electrocardiogram, heart rate, and respiration (used for MR artifact correction) at a sampling rate of 1000 Hz. We placed MRI-compatible EEG amplifiers (BrainAmp MR plus; Brain Products GmbH, Gilching, Germany) behind the MR scanner and attached cables to the participants once they were located in final position in the scanner. Furthermore, we fixated cables using sand-filled pillows to reduce artifacts induced through cable movement in the magnetic field. During functional scans, the MR helium pump was switched off to reduce EEG artifacts. After the scanning, we recorded the exact EEG electrode locations on participants’ heads relative to three fiducial points using a Polhemus FASTRAK device. For four participants, no such data were available due to time constraints/ technical errors, in which case we used the average electrode locations of the remaining 32 participants.

### EEG pre-processing

First, raw EEG data were cleaned from MR scanner and cardioballistic artifacts using BrainVisionAnalyzer^97^. The rest of the pre-processing was performed in Fieldtrip^98^. After rejecting channels with high residual MR noise (mean 4.8 channels per participant, range 1–13), we epoched trials into time windows of −1,400–2,000 ms relative to the onset of outcomes. Timing of this epochs was determined by the minimal inter-stimulus interval beforehand until the minimal inter-trial interval afterwards. Data was re-referenced to the grand average, which allowed us to recover the reference as channel FCz, and then band-pass filtered using a two-pass 4th order Butterworth IIR filter (Fieldtrip default) in the range of 0.5–35 Hz. These filter settings allowed us to distinguish the delta, theta, alpha, and beta band, while filtering out residual high-frequency MR noise. This low-pass filter cut-off was different from a previous analysis of this data in which we set it at 15 Hz^17^ because, in this analysis, we had a hypothesis on outcome valence encoding in the beta range. We then applied linear baseline correction based on the 200 ms prior to cue onset and used ICA to detect and reject independent components related to eye-blinks, saccades, head motion, and residual MR artifacts (mean number of rejected components per participant: 32.694, range 24–45). Afterwards, we manually rejected trials with residual motion (for all 36 participants: *M* = 117.722, range 11–499). Based on trial rejection, four participants for which more than 211 (33%) of trials were rejected were excluded from any further analyses (rejected trials after excluding those participants: *M* = 81.875, range 11–194). Finally, we computed a Laplacian filter with the spherical spline method to remove global noise (using the exact electrode positions recorded with Polhemus FASTRAK), which we also used to interpolate previously rejected channels. This filter attenuates more global signals (e.g., signal from deep sources or global noise) and noise (heart-beat and muscle artifacts) while accentuating more local effects (e.g., superficial sources).

### EEG TF decomposition

We decomposed the trial-by-trial EEG time series into their time-frequency representations using 33 Hanning tapers between 1 and 33 Hz in steps of 1 Hz, every 25 ms from −1000 until 1,300 ms relative to outcome onset. We first zero-padded trials to a length of 8 sec. and then performed time-frequency decomposition in steps of 1 Hz by multiplying the Fourier transform of the trial with the Fourier transform of a Hanning taper of 400 ms width, centered around the time point of interest. This procedure results in an effective resolution of 2.5 Hz (Rayleigh frequency), interpolated in 1 Hz steps, which was more robust to the choice of exact frequency bins. To exclude the possibility of slow drifts in power over the time course of the experiment, we performed baseline correction across participants and trials by fitting a linear model for each channel/ frequency combination with trial number as predictor and the average power 250–50 ms before outcome onset as outcome, and subtracting the power predicted by this model from the data. This procedure is able to remove slow linear drifts in power over time from the data. In absence of such drifts, it is equivalent to correcting all trials by the grand mean across trials per frequency in the selected baseline time window. Afterwards, we averaged power over trials within each condition spanned by performed action (Go/ NoGo) and outcome (reward/ no reward/ no punishment/ punishment). We finally converted the average time-frequency data per condition to decibel to ensure that data across frequencies, time points, electrodes, and participants were on same scale.

### EEG analyses

All analyses were performed on the average signal of a-priori selected channels Fz, FCz, and Cz based on previous literature^9,17^. We again performed model-free and model-based analyses. For the model-free analyses, we sorted trials based on the performed action (Go/ NoGo) and obtained outcome (reward/ no reward/ no punishment/ punishment) and computed the mean TF power across trials for each of the resultant eight conditions for each participant. We tested whether theta power (average power 4–8 Hz) and beta power (average power 13–30 Hz) encoded outcome valence by contrasting positive (reward/ no punishment) and negative (no reward/ punishment) conditions (irrespective of the performed action). We also tested for differences between Go and NoGo responses in the lower alpha band (6–10 Hz). For all contrasts, we employed two-sided cluster-based permutation tests in a window from 0– 1,000 ms relative to outcome onset. For beta power, results were driven by a cluster that was at the edge of 1,000 ms; to more accurately report the time span during which this cluster exceeded the threshold, we extended the time window to 1,300 ms in this particular analysis. Such tests are able to reject the null hypothesis of exchangeability of two experimental conditions, but they are not suited to precisely locate clusters in time-frequency space. Hence, interpretations were mostly based on the visual inspection of plots of the signal time courses.

For model-based analyses, similar to fMRI analyses, we used the group-level parameters from the best fitting computational model M5 to compute the trial-by-trial biased PE term and decomposed it into the standard PE term and the difference to the biased PE term. We used both terms as predictors in a multiple linear regression for each channel-time-frequency bin for each participant, and then performed one-sample cluster-based permutation-tests across the resultant *b*-maps of all participants^99^. For further details on this procedure, see fMRI-inspired EEG analyses.

### fMRI-informed EEG analyses

The BOLD signal is sluggish. It is thus hard to determine when different brain regions become active. In contrast, EEG provides much higher temporal resolution. A fruitful approach can be to identify distinct EEG correlates of the BOLD signal in different regions, allowing to test hypotheses about the temporal order in which regions might become active and modulated EEG power ^17,74^. Furthermore, by using the BOLD signal from different regions in a multiple linear regression, one can control for variance shared among regions (e.g., changes in global signal; variance due to task regressors) and test which region is the best unique predictor of a certain EEG signal. In such an analysis, any correlation between EEG and BOLD signal from a certain region reflects an association above and beyond those induced by task conditions.

We used the trial-by-trial BOLD signal in selected regions in a multiple linear regression to predict EEG signal over the scalp^17,74^ (building on existing code from https://github.com/tuhauser/TAfT; see Supplementary Fig. 17 for a graphical illustration). As a first step, we extracted the volume-by-volume signal (first eigenvariate) from each of the seven regions identified to encode biased PEs (conjunction of PE_STD_ and PE_DIF_: striatum, dACC, pgACC, left motor cortex, PCC, left ITG, and primary visual cortex). We applied a highpass-filter at 128 s and regressed out nuisance regressors (6 realignment parameters, CSF, OOB, single volumes with strong motion, same as in the fMRI GLM). We then upsampled the signal by a factor 10, epoched it into trials of 8 s duration, and fitted a separate HRF (based on the SPM template) to each trial (58 upsampled data points), resulting in trial-by-trial regression weights reflecting the respective BOLD response. We then combined the regression weights of all trials and regions of a certain participant into a design matrix with trials as rows and the seven ROIs as columns, which we then used to predict power at each time-frequency-channel bin. As further control variables, we added the behavioral PE_STD_ and PE_DIF_ regressors to the design matrix. All results were identical with and without the inclusion of PEs as covariates in the regression, suggesting that EEG-fMRI correlations did not merely arise from both modalities encoded PEs as a “common cause” that induced correlations. Instead, these correlations reflected the incremental variance explained in EEG power that was afforded by the BOLD signal even beyond the PEs. All predictors and outcomes were demeaned such that the intercept became zero. Such a multiple linear regression was performed for each participant, resulting in a time-frequency-channel-ROI *b*-map reflecting the association between trial-by-trial BOLD signal and TF power at each time-frequency-channel bin. *B*-maps were Fisher-*z* transformed, which makes the sampling distribution of correlation coefficients approximately normal and allows for combining them across participants. Finally, we tested for fMRI-EEG associations with a cluster-based one-sample permutation *t*-test ^99^ on the mean regression weights over channels Fz, FCz, and Cz across participants in the range of 0–1000 ms, 1–33 Hz. We first obtained a null distribution of maximal cluster mass statistics from 10000 permutations. For each permutation, we flipped the sign of the *b*-map of a random subset of participants, computed a separate *t*-test at each time-frequency bin (bins of 25 ms, 1 Hz) across participants (results in *t*-map), thresholded these maps at |t| > 2, and finally computed the maximal cluster mask statistic (sum of all *t*-values) for any cluster (adjacent voxels above threshold). Afterwards, we computed the same *t*-map for the real data, identified the cluster with the biggest cluster-mass statistic, and computed the corresponding *p*-value as number of permutations in the null distribution that were larger than the maximal cluster mass statistic in the real data.

### EEG-informed fMRI analyses

For the EEG-informed fMRI analyses, we fit three additional GLMs for which we entered the trial-by-trial theta/ delta power (1–8 Hz), beta power (13–30 Hz), and lower alpha band power (6–10 Hz) as parametric regressors on top of the task regressors of the model-free GLM. These measures were created by using the 3-D (time-frequency-channel) *t*-map obtained when contrasting positive vs. negative outcomes (theta/ delta and beta; Fig. 4 A, B) and Go vs. NoGo conditions (lower alpha band) as a linear filter (Fig. 4; see Supplementary Fig. 18 for a graphical illustration of this approach). Note that these signals were selected based on the EEG-only results and not informed by the fMRI-informed EEG analyses. We enforced strict frequency cut-offs. For lower alpha band and beta, we used midfrontal channels (Fz/ FCz/ Cz). For theta/ delta power, given the topography that reached far beyond midfrontal channels and over the entire frontal scalp, we used a much wider ROI (AF3/ AF4/ AF7/ AF8/ F1/ F2/ F3/ F4/ F5/ F6/ F7/ F8/ FC1/ FC2/ FC3/ FC4/ FC5/ FC6/ FCz/ Fp1/ Fp2/ Fpz/ Fz). We extracted those maps and retained all voxels with t > 2. These masks were applied to the trial-by-trial time-frequency data to create weighted summary measures of the average power in the identified clusters in each trial. For trials for which EEG data was rejected, we imputed the participant mean value of the respective action (Go/ NoGo) x outcome (reward/ no reward/ no punishment/ punishment) condition. Note that this approach accentuates differences between conditions, which were already captured by the task regressors in the GLM, but decreases trial-by-trial variability within each condition, which is of interest in this analysis. This imputation approach is thus conservative. While trial-by-trial beta and theta power were largely uncorrelated, mean *r* = 0.104, range −0.118–0.283 across participants, and so were beta and alpha, mean *r* = 0.097, range −0.162–0.284 across participants, theta and alpha power moderately correlate, mean *r* = 0.412, range 0.121–0.836 across participants, warranting the use of a separate channel ROI for theta and using separate GLMs for each frequency band.

### Analyses of behavior as a function of BOLD signal and EEG power

We used mixed-effects logistic regression to analyze “stay behavior”, i.e., whether participants repeated an action on the next encounter of the same cue, as a function of BOLD signal and EEG power in selected regions. For analyses featuring BOLD signal, we used the trial-by-trial HRF amplitude also used for fMRI-informed EEG analyses. For analyses featuring EEG, we used the trial-by-trial EEG power also used in the EEG-informed fMRI analyses.

## Acknowledgments

We thank Emma van Dijk for assistance with data collection, Michael J. Frank for helpful discussions, and the weekly Donders M/EEG meeting for discussions of these results and many helpful suggestions.

## Author contributions

Conceptualization: JA, JCS, RC, HEMDO

Data curation: JA

Formal analysis: JA

Funding acquisition: JCS, RC, HEMDO

Investigation: JA, JCS

Methodology: JA, HEMDO

Project administration: JA, JCS, HEMDO

Resources: RC, HEMDO

Software: JA, JCS, HEMDO

Supervision: JCS, RS, RC, HEMDO

Validation: JA, JCS, RS, RC, HEMDO

Visualization: JA

Writing – original draft: JA, HEMDO

Writing – review & editing: JA, JCS, RS, RC, HEMDO

## Data and code availability statement

All raw data is available under: https://doi.org/10.34973/pezs-pw62. All code required to achieve the reported results as well as preprocessed data and fMRI results are available under: [All data and code will be made available upon 19 manuscript acceptance]. In line with requirements of the Ethics Committee and the Radboud University security officer, potentially identifying data (such as imaging data) can only be shared to identifiable researchers. Hence, researchers requesting access to the data have to register and accept a data user agreement; access will then automatically be granted via a “click-through” procedure (without involvement of authors or data stewards).

Group-level unthresholded fMRI z-maps are available on Neurovault (https://neurovault.org/collections/11184/).

## Supplementary Information

### Supplementary Note 1: Behavioral results with only the 29 participants included in EEG-fMRI analyses

We repeated the behavioral analyses reported in the main text while excluding the seven participants that were also not included in the fMRI-inspired EEG analyses in the main text: (a) two participants due to fMRI co-registration failure, which were also not included in the fMRI-only analyses; (b) four further participants who exhibited excessive residual noise in their EEG data (> 33% rejected trials) and were thus also not included in the EEG-only analyses, and finally (c) one more participant who (together with four other participants already excluded) exhibited regression weights for every regressor about ten times larger than for other participants.

Participants in this subgroup learned the task, reflected in a significant main effect of required action on responses, *b* = 0.896, *SE* = 0.129, χ^2^(1) = 28.398, *p* < .001, and exhibited motivational biases, reflected in a significant main effect of cue valence on responses, *b* = 0.439, *SE* = 0.084, χ^2^(1) = 19.308, *p* < .001. The interaction between required action and cue valence was not significant, *b* = 0.025, *SE* = 0.085, χ^2^(1) = 0.111, *p* = .739 (Supplementary Fig. 1A-B).

Participants in this subgroup also showed biased learning: They were more likely to repeat an action after a positive outcome (main effect of outcome valence: *b* = .0553, *SE* = 0.059, χ^2^(1) = 40.920, *p* < .001. After salient outcomes, they adjusted their responses more strongly after feedback on Go than on NoGo responses, in line with our model of biased learning and as reflected in a significant three-way interaction between action, salience, and valence, *b* = 0.266, *SE* = 0.055, χ^2^(1) = 16.862, *p* < .001. When only analyzing trials with salient outcomes, outcome valence was more likely to affect response repetition following Go relative to NoGo responses, *b* = 0.324, *SE* = 0.079, χ^2^(1) = 13.266, *p* < .001, with a stronger effect of outcome valence after Go responses, *b* = 1.342, *SE* = 0.120, χ^2^(1) = 49.003, *p* = .001, than NoGo responses, *b* = 0.693, *SE* = 0.129, χ^2^(1) = 18.988, *p* < .001 (Supplementary Fig. 1C).

In this subgroup of participants, Bayesian model selection clearly favored the full asymmetric pathways models featuring response and learning biases (M5, model frequency: 81.81%, protected exceedance probability: 100%; Supplementary Fig. 1D-H). In sum, behavioral results were qualitatively identical when analyzing only this subgroup of only 29 participants.

### Supplementary Note 2: Behavioral fMRI results with only the 29 participants included in EEG-fMRI analyses

We repeated the fMRI analyses reported in the main text while excluding the seven participants that were also not included in the fMRI-inspired EEG analyses in the main text: (a) two participants due to fMRI co-registration failure, which were also not included in the fMRI-only analyses; (b) four further participants who exhibited excessive residual noise in their EEG data (> 33% rejected trials) and were thus also not included in the EEG-only analyses, and finally (c) one more participant who (together with four other participants already excluded) exhibited regression weights for every regressor about ten times larger than for other participants.

We first repeated the model-free GLM just contrasting positive and negative outcomes. BOLD signal was higher for positive than negative outcomes in five clusters, namely in vmPFC, striatum, amygdala, and hippocampus (*z*_max_ = 5.65, *p* = 2.24e-25, 6110 voxels, MNI coordinates xyz = [6 30 - 12]), left superior lateral occipital cortex (*z*_max_ = 4.40, *p* = .00144, 367 voxels, xyz = [−46 −68 46]), right occipital pole (*z*_max_ = 4.45, *p* = .00154, 363 voxels, xyz = [12 −92 −12]), posterior cingulate cortex (*z*_max_ = 4.36, *p* = .00181, 353 voxels, xyz = [−2 −48 28]), and left middle temporal gyrus (*z*_max_ = 4.63, *p* = .00548, 289 voxels, xyz = [−60 −10 −16]; Supplementary Fig. 2A). The clusters in left slOCC, PCC, and left MTG emerged anew compared to the original analysis comprising 34 participants. Also, compared to the original analysis, clusters in left orbitofrontal cortex and left superior frontal gyrus were merged with the cluster in vmPFC. In sum, all clusters from the original analysis were found back, plus some additional clusters.

There was also one cluster in right orbitofrontal cortex (*z*_max_ = 4.37, *p* = .0209, 217 voxels, xyz = [30 62 −2]) in which BOLD signal was higher for negative than positive outcomes. Compared to the original analysis comprising 34 participants, clusters in precuneus and right superior frontal gyrus were not significant.

In the model-based GLM featuring regressors for standard PEs and the difference term towards biased PEs, BOLD signal correlated with standard PEs in ten clusters, namely in vmPFC, striatum, bilateral amygdala and hippocampus (*z*_max_ = 6.04, *p* = .4.78e-44, 8848 voxels, xyz = [12 14 −6]), left superior frontal gyrus (*z*_max_ = 5.58, *p* = 3.5e-10, 1043 voxels, xyz = [−18 34 52]), left occipital pole and lingual gyrus (*z*_max_ = 6.23, *p* = 7.18e-10, 998 voxels, xyz = [10 −92 −10]), posterior cingulate cortex (*z*_max_ = 5.12, *p* = 8.57e-10, 987 voxels, xyz = [4 −36 48]), left inferior temporal gyrus (*z*_max_ = 5.03, *p* = 7.07e-09, 859 voxels, xyz = [−52 −46 −10]), right anterior middle temporal gyrus (*z*_max_ = 5.32, *p* = .000292, 314 voxels, xyz = [62 −4 −16]), right cerebellum (*z*_max_ = 5.32, *p* = .002228, 231 voxels, xyz = [44 −72 −40]), left superior lateral occipital cortex (*z*_max_ = 4.69, *p* = .00322, 218 voxels, xyz = [−46 −74 −38]), right caudate (*z*_max_ = 4.33, *p* = .00538, 199 voxels, xyz = [20 12 22]), and right middle temporal gyrus (*z*_max_ = 4.09, *p* = .0129, 189 voxels, xyz = [54 −38 −12]; Supplementary Fig. 2B). The clusters in left superior lateral occipital cortex, right caudate, and right posterior middle temporal gyrus emerged anew by splitting from larger clusters visible in the original analysis based on 34 participants. Vice versa, the cluster in left middle temporal gyrus reported for the original analysis was merged with a bigger cluster in the analysis of only 29 participants. The clusters in postcentral gyrus and ACC observed in the original analysis based on 34 participants were not significant anymore; however, they were still visible at a level of *z* > 3.1 uncorrected.

BOLD signal correlated significantly negatively with standard PEs in a single cluster in right superior frontal gyrus (*z*_max_ = 5.04, *p* = .00771, 186 voxels, xyz = [6 26 64]), similar to the respective cluster reported in the original analysis. In contrast, the clusters in right occipital pole, intracalcarine cortex, and left inferior lateral occipital cortex were not significant any more, though visible at a level of *z* > 3.1 uncorrected.

BOLD signal in six clusters correlated significantly positively with the difference term towards biased PEs, namely in large parts of cortex and subcortex including striatum (*z*_max_ = 6-54, *p* = 0, 29428 voxels, xyz = [34 −84 20]), dorsomedial prefrontal cortex (*z*_max_ = 5.94, *p* = 2.69e-40, 7001 voxels, xyz = [6 22 34]), right insula (*z*_max_ = 5.76, *p* = 7.84e-27, 3847 voxels, xyz = [34 20 −8]), thalamus and brainstem (*z*_max_ = 5.10, *p* = 4.06e-18, 2169 voxels, xyz = [4 −30 0]), left caudate (*z*_max_ = 4.71, *p* = .000188, 305 voxels, xyz = [−12 8 6]) and another cluster in brainstem (*z*_max_ = 4.05, *p* = .0151, 160 voxels, xyz = [4 − 30 −30]). Clusters in dmPFC, right insula, and left caudate split from larger clusters reported in the original analysis. Vice versa, the cluster in left insula reported in the original analysis merged with the largest cluster. The clusters in right middle temporal gyrus and right insula were missing in the analysis of only 29 participants, but visible at a level of *z* > 3.1 uncorrected.

BOLD signal in three clusters correlated significantly negatively with the difference term towards biased PEs, namely in vmPFC (*z*_max_ = 4.23, *p* = .0051, 185 voxels, xyz = [−12 48 −6]), left hippocampus (*z*_max_ = 4.58, *p* = .00857, 168 voxels, xyz = [−26 −14 −22]), and left medial temporal gyrus (*z*_max_ = 4.30, *p* = .0172, 146 voxels, xyz = [−62 −4 −16]). Compared to the original analysis, the cluster in vmPFC emerged anew.

When computing the conjunction between both (positive) contrasts, BOLD signal encoded both the standard and the difference in four clusters, namely in vmPFC, bilateral striatum, bilateral ITG, and V1 (Supplementary Fig. 2C). Clusters in ACC, left motor cortex, and PCC were not significant any more (because they were z > 3.1, but not significant after cluster correction in the standard PE contrast). However, new (though rather small) clusters of biased PE encoding emerged in right insula, left amygdala, and left OFC. In sum, results when analyzing only this subgroup of only 29 participants were largely similar to results based on the full sample; however, clusters of biased PE encoding in left motor cortex, ACC, and PCC were small and thus did not survive cluster correction in this subgroup.

### Supplementary Note 3: EEG results with only the 29 participants included in EEG-fMRI analyses

We repeated the EEG analyses reported in the main text while excluding the seven participants that were also not included in the fMRI-inspired EEG analyses in the main text: (a) two participants due to fMRI co-registration failure, which were also not included in the fMRI-only analyses; (b) four further participants who exhibited excessive residual noise in their EEG data (> 33% rejected trials) and were thus also not included in the EEG-only analyses, and finally (c) one more participant who (together with four other participants already excluded) exhibited regression weights for every regressor about ten times larger than for other participants.

In participants in this subgroup, both midfrontal theta and beta power reflected outcome valence: Theta power was higher for negative than positive outcomes (driven by a cluster around 225–500 ms, *p* = .002; Supplementary Fig. 3A, B), while beta power was higher for positive than negative outcomes (driven by a cluster around 325–1000 ms, *p* = .002; Supplementary Fig. 3A, C). When using PE terms as regressor for midfrontal EEG power while controlling for PE valence, delta power did not encode *PE_STD_* positively, though not significant (p = .056), and also the positive encoding of *PE_DIF_* was non-significant (*p* = .053; Supplementary Fig. 3D-F). The positive correlation of beta power with *PE_STD_* was not significant anymore (*p* = .059), while the negative correlation with *PE_DIF_* remained (p = .001, 450– 950 ms). When adding *PE_STD_* and *PE_DIF_* together to achieve *PE_BIAS_*, theta/delta power indeed significantly encoded *PE_BIAS_*, first positively (*p* = .032, 224–475 ms) and then negatively (*p* = .019, 600 – 1,000 ms; around 8 Hz and thus rather in the alpha band). Also, beta power was significantly negatively correlated with *PE_BIAS_* (*p* = .008, 450 – 975 ms).

In sum, all findings reported in the main text also held when analyzing only this subgroup of only 29 participants. In addition, also late beta power and theta/alpha power appeared to negatively encode the *PE_BIAS_* term.

### Supplementary Note 4: EEG and fMRI correlates of past action with only the 29 participants included in EEG-fMRI analyses

We repeated the behavioral analyses reported in the main text while excluding the seven participants that were also not included in the fMRI-inspired EEG analyses in the main text: (a) two participants due to fMRI co-registration failure, which were also not included in the fMRI-only analyses; (b) four further participants who exhibited excessive residual noise in their EEG data (> 33% rejected trials) and were thus also not included in the EEG-only analyses, and finally (c) one more participant who (together with four other participants already excluded) exhibited regression weights for every regressor about ten times larger than for other participants.

Regarding fMRI correlates of the past action, similar to the original analysis comprising 34 participants, there were no clusters with higher BOLD after Go than NoGo actions at the time of outcomes, but vice versa, large parts of cortex and subcortex showed higher BOLD after NoGo than Go actions, highly similar to the original analysis (*z*_max_ = 7.65, *p* = 0, 124629 voxels, xyz = [−58 18 22]; Supplementary Fig. 4D).

Furthermore, there were four clusters with higher BOLD for Go than NoGo actions at the time of the response, namely one large cluster across lateral prefrontal cortex, anterior cingulate cortex, striatum, thalamus, angular gyrus, cerebellum, left operculum and motor cortex, intracalcarine cortex, and occipital pole (*z*_max_ = 7.45, *p* = 0, 61057 voxels, xyz = [32 −4 −4]), one in right middle temporal gyrus (*z*_max_ = 4.90, *p* = 8.66e-05, 493 voxels, xyz = [66 −32 −12]), one in left inferior temporal gyrus (*z*_max_ = 4.43, *p* = .00294, 293 voxels, xyz = [−60 −44 −18]), and one in precuneous (*z*_max_ = 2.39, *p* = .0041, 276 voxels, xyz = [−8 −70 38]; Supplementary Fig. 4C). All these regions were also found in the original analysis comprising 34 participants. Vice versa, BOLD signal was higher NoGo than Go actions at the time of the response in two clusters in vmPFC and subcallosal cortex (*z*_max_ = 4.23, *p* = .00864, 239 voxels, xyz = [−2 18 −6]) and right anterior temporal gyrus/ temporal pole (*z*_max_ = .4.14, *p* = .0193, 201 voxels, xyz = [48 −6 −8]), identical to the original analysis comprising 34 participants.

Finally, there was higher BOLD signal for left hand compared to right hand responses at the time of response in two clusters in right precentral and postcentral gyrus, superior parietal lobule, and operculum (*z*_max_ = 6.66, *p* = 0, 11597 voxels, xyz = [46 −24 64]) and left cerebellum (*z*_max_ = 6.76, *p* = 1.05e-18, 2672 voxels, xyz = [−18 −54 −16]; Supplementary Fig. 4C), identical to the original analysis comprising 34 participants. Vice versa, there was higher BOLD signal for right hand than left hand responses at the time of responses in five clusters in left precentral and postcentral gyrus, superior parietal lobule, operculum, and thalamus (*z*_max_ = 6.4, *p* = 0, 12372 voxels, xyz = [−36 −20 66]), right cerebellum (*z*_max_ = 7.17, *p* = 3.41e-21, 3206 voxels, xyz = [20 −54 −20]), right superior lateral occipital cortex (*z*_max_ = 4.84, *p* = 2.28e-09, 988 voxels, xyz = [48 −86 −4]), right angular gyrus (*z*_max_ = 4.11, *p* = 7.68e-05, 396 voxels, xyz = [66 −50 28]), and left superior lateral occipital cortex (*z*_max_ = 5.03, *p* = .019, 164 voxels, xyz = [−18 −82 48]). The clusters in right occipital pole/ intracalcarine cortex and in right posterior cerebellum observed in the original analysis comprising 34 participants were not observed in this analysis. In sum, all major findings also held when analyzing only this subgroup of only 29 participants.

Regarding EEG time-frequency correlates of the past action, when testing for differences in broadband after outcome onset, there was no significant difference after Go and NoGo responses, *p* = .283. When restricting analyses to the low alpha range, the permutation test was marginally significant, *p* = .056, driven by a cluster around 0–100 ms around 7–10 Hz; Supplementary Fig. 4A, B). When repeating the permutation test for the broadband signal including the last second before outcome onset, there was a significant difference after Go and NoGo responses, driven by clusters in the beta band. *p* = 0.002, −1000 – −275 ms, 13–32 Hz, and in the theta/ low alpha band, *p* = 0.020, −1000 – −525 ms, 4–10 Hz.

## Supplementary Note 5: Stay behavior as a function of EEG and fMRI with only the 29 participants included in EEG-fMRI analyses

When linking trial-by-trial BOLD signal in selected ROIs as well as midfrontal EEG TF power to response repetition on the next trial with the same cue, dACC BOLD signal did not significantly predict the response repetition, *b* = −0.013, *SE* = 0.018, χ^2^(1) = 0.524, *p* = .469, and neither did PCC BOLD signal, *b* = −0.037, *SE* = 0.018, χ^2^(1) = 2.079, *p* = .149. However, participants in this subgroup were significantly more likely to repeat the sample action when striatal BOLD signal was high, *b* = 0.097, *SE* = 0.025, χ^2^(1) = 12.043, *p* < .001, but more likely to switch when vmPFC BOLD was high, *b* = −0.075, *SE* = 0.019, χ^2^(1) = 13.170, *p* < .001.

When linking trial-by-trial midfrontal EEG TF power to response repetition on the next trial with the same cue, participants in this subgroup were more likely to repeat the same response when beta power was high, *b* = 0.124, *SE* = 0.036, χ^2^(1) = 3.502, *p* < .001, or when low alpha power was high, *b* = 0.135, *SE* = 0.044, χ^2^(1) = 8.789, *p* = .003, but more likely to switch to another response when theta power was high, *b* = −0.090, *SE* = 0.040, χ^2^(1) = 4.812, *p* = .028.

## Supplementary Note 6: Parameter recovery analyses for model M5

We performed parameter recovery analyses to assess the identifiability of the model parameters in the winning “asymmetric pathways” model M5. We simulated 100 new data sets based on the best fitting parameters of each participant, fitted a separate model to each simulated data set (using first Laplace approximation and then hierarchical Bayesian inference), and finally averaged parameters across the 100 fitted models.

Parameter recovery was excellent for the feedback sensitivity ρ (*r* = .91), the baseline learning rate ε_0_ (*r* = .98), the Go bias *b* (*r* > .99), and the Pavlovian response bias π (*r* > .99), with between-participant differences in ground-truth parameters correlating at high levels (all *r* > .90; Supplementary Fig. 5) with between-participant differences in the recovered parameters. Note that, due to shrinkage to the mean as a consequence of hierarchical Bayesian inference, extreme parameter values tended to be shrunk to the overall group-level mean in the recovered parameters. Correlations for the learning bias parameter κ were considerably lower, though still strongly positive (*r* = 0.50; *r* = 0.51 when removing one outlier participant; Supplementary Fig. 5E). Note however that the effect of κ on learning depended on participants’ baseline learning rate ε_0_. When computing increased learning rates for rewarded Go actions and decreased learning rates for punished NoGo actions—the parameters that determine the effective degree of trial-by-trial learning—these learning rates were again highly correlated with the ground truth parameters (*E_rewarded Go_* : *r* = 0.96; *E_punished NoGo_* : *r* = 0.85 resp. *r* = 0.86 when removing one outlier participant; Supplementary Fig. 5F-G).

Further parameter recovery analyses on the models explored in Supplementary Note 8 yielded that the recovery of κ was improved (*r* = 0.78) when adding perseveration parameters (which themselves had recovery performances of *r*’s > 0.99). This observation suggested that models featuring such perseveration parameters might be better suited for quantifying individual differences in the learning bias.

In sum, parameter recovery was excellent for all parameters but the learning bias κ. More relevant than recovery of κ, however, was that we could recover the effective learning rate well (combining baseline learning rate ε_0_ and the learning bias κ). However, when combining the baseline learning rate ε_0_ and the learning bias κ, recovery was high, as well. Note that the ability to accurately capture individual differences in biased learning is not of interest in this study, nor relevant to the imaging analyses. In fact, we used a single set of parameters (the group-level parameters) to compute trial-by-trial regressors for the EEG and fMRI analyses. This is a standard approach in model-based fMRI for two main reasons. First, it has been shown that the exact parameter values for relatively simple RL models like the ones used here have little impact on the results of fMRI analyses ^1^. For the current study, of most relevance is the qualitatively differential pattern of learning updates after Go and NoGo responses ^2–4^, as embodied by the algorithmic specification of the model. This pattern drives the EEG and fMRI results and indeed, using a different set of parameter values, we obtain essentially identical fMRI results (see Supplementary Note 9 and Supplementary Fig. 8).

## Supplementary Note 7: Simulations for asymmetric pathways and action priming model

Motivational learning biases are predicted by the *asymmetric pathways model* ^5,6^: Positive PEs, elicited by rewards, lead to long-term potentiation in the striatal direct “Go” pathway (and long term depression in the indirect pathway), allowing for a particularly effective acquisition of Go actions to obtain rewards. Conversely, negative PEs, elicited by punishments, lead to long term potentiation in the NoGo pathway, impairing the unlearning of NoGo actions in face of punishments.

An alternative account has recently suggested that self-generated (Go) actions lead to preferential learning (relative to non-self-generated actions, including inaction), more generally (henceforth called “action priming model”)^7^. A self-generated action could “prime” basal ganglia circuits and lead to subsequently larger PEs and thus faster learning. The main differential prediction between these two models is how they account for the failure to learn “Go” actions to avoid punishment: In the first model, this is due to a failure to unlearn punished “NoGo” actions, while in the second model, this is due to increased unlearning of punished “Go” actions.

Here, we directly tested both models against each other. We specified an alternative model M6 ^7^ with two separate learning rates, one learning rate for trials where self-generated (Go) action selection should prime the processing of any following salient outcome (i.e., Go actions followed by rewards/ punishments), and one learning rate for any other action-outcome combination. In this model, equation (6) was substituted by equation (7):

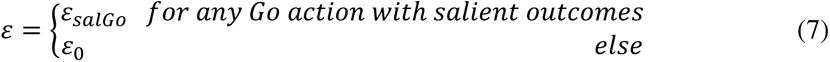

When comparing all models M1–M6 using Bayesian model selection, M5 (the asymmetric pathways model) received highest support (model frequency: 68.15%; protected exceedance probability: 99.70%), also compared to M6 (the action priming model; model frequency: 24.19%; protected exceedance probability: 0.30%; Supplementary Fig. 6D, H). In fact, as visible in Supplementary Fig. 6E-G, the action priming did not reproduce the motivational biases in learning curves and bar plots, which constitutes a case of qualitative model falsification ^2,3^. If anything, it seemed that the action priming model traded off both biases, leading to negative response biases for a majority of participants. In contrast, the asymmetric pathways model (M5) was well able to capture the qualitative patterns observed in the data (Supplementary Fig. 6A-C). We conclude that only the asymmetric pathways model is able to qualitatively reproduce core characteristics of our data.

## Supplementary Note 8: Behavioral results for the perseveration model (M7), cue valence-based perseveration model (M8), and neutral outcomes reinterpretation model (M9)

While the winning model M5 reported in the main text captured learning curves and the proportion of (correct/ incorrect) Go and NoGo responses well, it did not fully capture the propensity to stay (i.e., repeat the same response to the subsequent presentation of the same cue) following different action-outcome combinations (see Fig. 2G in the main text). Specifically, M5 underestimated the overall propensity to stay and predicted a higher probability of repeating a Go response after a positive (neutral) outcome for Avoid cues, relative to the negative (neutral) outcome for Win cues. In contrast, in the data, there was no such significant difference. We thus explored three extensions of M5 that had the potential to capture this behavioral pattern. Specifically, we considered mechanisms that would make the model more likely to repeat a given response. Furthermore, any such mechanism should boost repetition of Go responses to non-rewarded Win cues particular. We hypothesized that two potential mechanisms could account for these data features, and present three new models to test these mechanisms.

As a first mechanism, we considered overall “response stickiness” or “perseveration”^8^, a process that leads participants to repeat a previous response independent of the obtained outcome. This mechanism could explain participants’ overall higher propensity to stay, which we tested in model M7. ***Model M7***, called “***single perseveration model***”, featured the same parameters as M5 plus a perseveration parameter *φ* that was added as a “bonus” to the action weight *w(a_i_, s_t_)* of the specific action shown on the last occurrence of the respective cue^8^:

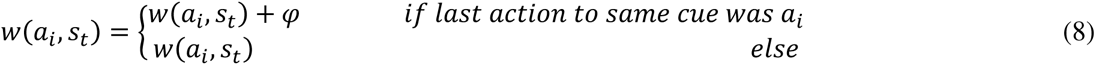

**In M7** equation 7 in the main manuscript was replaced by equation 8 above, such that parameter *φ* captured the propensity to repeat the action from the last time this cue was presented.

However, to account for the fact that staying was not different and numerically even higher for a non-rewarded Go response (to a Win cue), relative to a non-punished Go response to an Avoid cue, we tested whether separate perseveration parameters for Win and Avoid cues could capture this behavioral difference (M8), as such a pattern of results could result from an overall higher propensity to stay for Win cues. **This “cue valence-dependent perseveration model” (M8)**, contained two separate perseveration parameters, one for Win cues *φ_WIN_*, and one for Avoid cues *φ_AVOID_*. The respective perseveration parameter was added to the action weight *w(a_i_, s_t_)* of the specific action shown on the last occurrence of respective the cue:

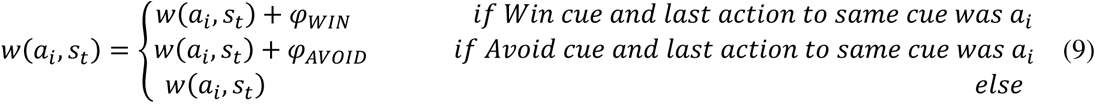

**In M8,** equation 7 in the main manuscript was replaced by equation 9 above, such that parameter *φ_WIN_* and *φ_AVOID_* captured the propensity to repeat the action from the last time this cue was presented, separately for Win and Avoid cues.

As an alternative mechanism that could potentially capture the p(stay) pattern in the data, we considered the possibility that participants might “re-interpret” neutral outcomes in line with the cue valence: although a non-reward after a Win cue constitutes negative feedback, the positive cue valence might “overshadow” this feedback and give participants the impression that they received a reward. Similarly, a non-punishment after an Avoid cue constitutes positive feedback, but the negative cue valence might overshadow this feedback and give participants the impression that they received a punishment.

Following this idea, lastly, we considered **M9,** called the “**neutral outcome reinterpretation model**”, which featured a single perseveration parameter φ as in equation (8), but in addition replaced neutral outcomes (coded as zero) with what we term the “effective reward” *r_EFF_*, which allows the neutral outcome to take on a value in the direction of the cue valence *V(s)*. The degree to which this happens is scaled by the parameter η:

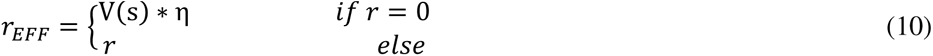

We subsequently used *r_EFF_* for computing prediction errors. Thus **M9** adds equation 10 to model **M7**. Note that for *η = 0*, neutral outcomes stay at zero and M9 becomes equivalent to M7.

Bayesian model comparison across the winning original model M5 and these three new models yielded highest model evidence for M8, followed by M9 (model frequency: M5: 3%, M7: 0%, M8: 62%, M9: 35%; protected exceedance probability: M5: 0%, M7: 0% M8: 95%, M9: 5%). All three models performed better than the original winning model M5 (Supplementary Fig. 7, bottom row). Simulations showed that the best fitting model M8 (with separate perseveration rates for Win and Avoid cues) indeed better captured the propensity to stay on neutral trials, though this came at the cost of a general overestimation of staying after punished responses (which hold similarly for M7 and M9; see Supplementary Fig. 7, third row). More importantly, however, this model drastically underestimated the crucial pattern of behavior under study here, namely the propensity of incorrect, bias-driven Go responses to Win cues (see Supplementary Fig. 7, second row, dark green part of bars).

In sum, the three additional models provided a better quantitative fit to the data compared to the winning model M5 reported in the main text. Also, these additional models predicted the propensity more accurately than the base models did. However, their qualitative fit (i.e. the ability to capture relevant aspects of the data) was worse: These additional models systematically underestimated the proportion of incorrect Go responses (Supplementary Fig. 7). Furthermore, although the predicted patterns of the propensity to stay matched the data more closely than M5, these predicted patterns still mis-matched some aspects of the data, particularly now over-estimating the tendency to stay following a punishment. Taken together, these models could capture certain qualitative patterns in the data, but not others, which is a core feature of computational modelling, which by definition constitutes a data reduction procedure that necessarily loses some details of the data. In terms of qualitative model validation/ falsification^2,3^, M5 and M8/M9 capture different qualitative features of the data, but no model captured all features well.

### Supplementary Note 9: Neural results based on prediction-errors from the cue valence-based perseveration model (M8) and neutral outcomes reinterpretation model (M9)

To confirm that neural correlates of biased prediction-error updating were not altered under these alternative model specifications, we repeated the model-based fMRI analyses for both the cue valence-dependent perseveration model M8 and the neutral outcomes interpretation model M9. In summary, the results are effectively unchanged, as we present in more detail below.

Notably, M8 does not make different predictions about trial-by-trial learning updates; the only difference to M5 consisted in slightly different best fitting parameter estimates for ε and κ (leading a slightly different BOLD regressors. Neural correlates of learning typically reflect the qualitative learning pattern, which is the same for M5 and M8, but are hardly sensitive to the exact parameter values ^1^. Indeed, when repeating the fMRI analyses with those different parameter values, we found almost identical results, with significant encoding of both PE_STD_ and PE_DIF_ in striatum, dACC, pgACC, PCC, left motor cortex, left ITG, and V1 (Supplementary Fig. 8A, B). The only exception was the cluster in dACC, which under M8 was not significant at a whole-brain level, but significant when using small-volume correction with an anatomical ACC mask (from the Harvard-Oxford Atlas), warranted by our a-priori hypotheses based on previous literature^9^.

When we repeated our fMRI analyses with learning updates predicted by M9, we again found significant encoding of both PE_STD_ and PE_DIF_ in striatum, dACC, pgACC, PCC, left motor cortex, left ITG, and V1 (Supplementary Fig. 8C). However, the pgACC cluster was much larger and extended into the vmPFC. Similarly, the PCC cluster was much larger. In addition, BOLD signal in left inferior frontal gyrus and in multiple clusters in superior and inferior lateral occipital cortex encoded both PE_STD_ and PE_DIF_ significantly. Using trial-by-trial BOLD signal from the extended vmPFC and PCC clusters identified with M9 regressors to predict midfrontal EEG power, we obtained results that were highly similar to the results for the pgACC and PCC clusters identified with M5 regressors.

In sum, model-based fMRI analyses based on PEs derived from M8 and M9 replicated the findings based on M5 reported in the main text. In addition, M9 led to larger clusters in vmPFC and PCC, tentatively suggesting that these regions might potentially contribute to “reinterpreting” neutral outcomes in light of the previously presented cue valence (see also Fig. 2 in the main text).

### Supplementary Note 10: EEG time-frequency results after ERPs were removed

Given that differences in theta power between positive and negative outcomes as well as differences in lower alpha band power after Go and NoGo responses occurred quite soon after cue onset, we aimed to test whether these effects reflected differences in evoked rather than induced activity. For this purpose, we removed evoked components from our data by computing the ERP for each of the eight conditions (action x outcome) for each participant and then subtracting the condition-specific ERP from the trial-by-trial data ^10^. Only afterwards, we performed time-frequency decomposition.

In line with the results reported in the main text, power was higher for negative compared to positive outcomes in the theta band (*p* =.018, driven by cluster at 225–475 ms; Supplementary Fig. 11A, B), but higher for positive than negative outcomes in the beta band (*p* < .001, driven by cluster at 0– 1250 ms; Supplementary Fig. 11A, C). Notably, unlike the results reported in the main text (Fig. 4A), the cluster of high power for negative compared to positive outcomes was constrained to the theta range, and did not extend further into the delta range (Supplementary Fig. 11A).

When using the trial-by-trial PEs (both the standard PE and the difference term to a biased PE) as predictors in a multiple linear regression at each time-frequency-channel bin while controlling for PE valence, delta power encoded *PE_STD_* positively, though not significantly (*p* = .198). However, at a later time point around outcome offset, delta (and theta) power in fact correlated negatively with *PE_STD_* (575– 800 ms, *p* = .002; Supplementary Fig. 11E). The correlation between delta and the *PE_DIF_* term was still positive, but not significant (p = .228; Supplementary Fig. 11F). Similarly, the correlation of the *PE_BIAS_* term with delta power was positive, but not significant (*p* = .084; Supplementary Fig. 11D).

Regarding beta power, there was a positive, though non-significant correlation of beta power with *PE_STD_* (*p* = .096; Supplementary Fig. 11E). There was again a significantly negative correlation of beta power with *PE_DIF_* (425–875 ms, *p* < .001; Supplementary Fig. 11F). Likewise, beta power correlated significantly negatively with *PE_BIAS_* (450–800 ms, *p* = .018; Supplementary Fig. 11D), driven by the correlation with *PE_DIF_*.

In sum, after subtracting the condition-wise ERP from each trial before time-frequency decomposition, supposedly removing the phase-locked aspect of power, both beta and theta still encoded PE valence. However, the encoding of PE magnitude by delta power was attenuated and not significant any more.

This reduction in magnitude encoding might occur of several reasons. Firstly, it might be that this correlation in the delta range was in fact (partly) reflecting correlations with phase-locked, i.e., evoked activity (ERPs), especially in the N2 (FPN)/ P3 (RewP) time range (see Supplementary Note 11 and Supplementary Fig. 12)^11–20^. Nonetheless, a positively correlation between delta power and biased PEs was still visible in Supplementary Fig. 11D, suggesting that at least part of the signal encoding biased PEs was not phase-locked. Secondly, it might be that the removal of the condition-wise ERPs has introduced additional noise in the data, attenuating any true correlation. Thirdly, there was a negative correlation between *PE_STD_* and theta/ delta power at later time points which was visible, though not significant in the results reported in the main text (Fig. 4D). Subtraction of an ERP-like template acts like a high-pass filter. High-pass filtering at relatively high cut-offs (> 0.5 Hz) can artificially postpone or induce effects at later points ^21^. It is possible that in this case, ERP subtraction attenuated a positive correlation in the theta/ delta range, but enhanced a later negative correlation.

Taken together, it is possible that part of the PE magnitude encoding in the theta/ delta range is due to correlations with the phase-locked (ERP) signal. However, this finding did not compromise the conclusion that overall, theta/delta power seemed to be more strongly associated with the *PE_BIAS_* term than the *PE_STD_* term. Our primary goal was not to pinpoint the precise nature of electrophysiological correlates of biased learning, but rather test the relative temporal order of when different regions exhibiting biased learning signals become active.

Finally, we tested whether after ERP subtraction, low alpha (and beta power) still encoded the previously performed action. When testing for differences in broadband power after Go and NoGo responses, power was indeed significantly different between conditions, driven by clusters in beta band (*p* = 0.002, 0.125–625 ms; *p* = 0.052, 700–1000 ms, 23–29 Hz) and theta/ low alpha band (*p* = 0.024, 575–1000 ms, 5–9 Hz; *p* = 0.056, 0–225 ms, 6–11 Hz). For power before outcome onset, there were again broadband differences between Go and NoGo (*p* = 0.002, −1000 – +225 ms, 1–33 Hz), but note that there was no ERP subtracted before outcome onset. We thus conclude that the differences between Go and NoGo responses were attributable to differences in induced rather than evoked activity.

### Supplementary Note 11: ERPs as a function of action and outcome

In addition to the induced activity in time-frequency power reported in the main text, we also analyzed the data in the time domain to test for differences in evoked activity. These analyses were particularly motivated given that differences in time-frequency power between positive and negative outcomes (theta/delta range) and after Go and NoGo responses (lower alpha/ theta range) occurred soon after outcome onset, warranting the assumption that differences might also occur in evoked activity. A large range of previous research has reported a modulation of evoked potentials by outcome valence in form of the feedback-reduced negativity ^14–20,22^, i.e., a stronger N2 component for negative compared to positive outcomes around ∼ 250 post-cue over midfrontal electrodes, recently also characterized as rather constituting a reward positivity (RewP) ^14^. Also, some studies have reported a modulation of the P3 by outcome valence, which has been attributed to outcome magnitude or salience rather than valence^17,18,20,23^.

Similar to the analysis of time frequency power, we sorted trials into the eight conditions spanned by the performed action (Go/ NoGo) and the obtained outcome (reward/ no reward/ no punishment/ punishment), computed the average ERP for each condition per participant, and tested for differences between positive (reward/ no punishment) and negative (no reward/ punishment) outcomes as well as conditions of relative stronger (rewarded Go and punished Go) vs. relatively weaker learning (rewarded NoGo and punished NoGo). We used cluster-based permutation tests on the average signal over midfrontal electrodes (Fz/ FCz/ Cz) in the time range of 0–700 ms after outcome onset (where evoked potentials visible in condition-averaged plot).

First, midfrontal ERPs were significantly different between positive and negative outcomes, driven by two separate clusters of differences above threshold (Cluster 1: around 246 – 294 ms, *p* = .034; Cluster 2: around 344 – 414 ms, *p* =.004; Supplementary Fig. 12A, C). The first cluster the classical feedback-related negativity, i.e., a stronger N2 component for negative compared to positive outcomes. The second cluster reflected weaker P3 component for negative compared to positive outcomes, similar the reward positivity reported before. In fact, the N3 was rather absent for negative outcomes (Supplementary Fig. 13). Both effects were clearly focused on midfrontal electrodes. These findings replicate previous findings of outcome valence modulating N2 (feedback-related negativity) and P3 components, and complement our time-frequency findings of theta and beta power reflecting outcome valence.

Second, when contrasting trials with Go vs. NoGo responses, no significant difference was observed (*p* = .358; Supplementary Fig. 12D). Visual inspection of the topoplot yielded that, if anything, differences emerged over right occipital electrodes. If one performed a test over those right occipital electrodes (O2, 04, PO4; Supplementary Fig. 12F; note that this procedure constitutes double-dipping because the test was informed by first looking at the data), this test would have yielded significant results (*p* = .016) driven by cluster around 423–466 ms, reflecting a slightly larger P3 after Go than NoGo responses (Supplementary Fig. 12E). This finding appears to be the strongest (if any) difference in amplitude after outcome onset between Go and NoGo actions. Given that this difference was not hypothesized and occurred far away from our a-priori selected channels of interest, we are careful not to over-interpret those differences.

Third, contrasting trials with positive and negative at the same right occipital electrodes yielded a significant difference, driven by clusters around 46–103 ms (*p* = 0.034), 141–255 ms (*p* = .002), and 519 – 580 ms (*p* = .034). Most notably, the P1 amplitude was much larger for positive than negative outcomes (Supplementary Fig. 12B). However, given that these differences were not hypothesized and occurred far away from our a-priori selected channels of interest, we are careful not to over-interpret those differences.

Taken together, we found a bigger midfrontal N2/ FRN for negative compared to positive outcomes, and a bigger midfrontal P3/ RewP for positive compared to negative outcomes, in line with a vast literature of previous findings ^14–20,22,23^. Midfrontal voltage did not significantly differ after Go or NoGo responses. If anything, differences after Go and NoGo responses were maximal over right occipital electrodes, with a larger P3 after Go than after NoGo responses. Signal at these channels also differed between positive and negative outcomes, most notably with a bigger P1 after positive than negative outcomes. In sum, we replicate classical reward learning ERP effects, which shows that the motivational Go/NoGo learning task taps into reward learning processes reported before, but these processes appeared to be unaffected by the previously performed action.

### Supplementary Note 12: Model-based EEG analyses in the time domain

In addition to testing whether midfrontal time-frequency power reflected signatures of biased learning (see main text), we also tested whether the midfrontal time domain signal reflected biased learning. Again, we used the standard PE term and the difference term to biased PEs as regressors in a multiple linear regression on each channel-time bin.

Focusing on midfrontal electrodes, and controlling for outcomes valence, first, the *PE_STD_* term was negatively correlated with midfrontal voltage around 529–575 ms (p = .039; Supplementary Fig. 14B). Note that so late after outcome onset, signal was not part of any “classical” ERP component any more. Second, the *PE_DIF_* correlated negatively with midfrontal voltage around 123–166 ms (*p* = .029) in the time range of the N1 and later positively around 365–443 ms (*p* < .001; Supplementary Fig. 14C) in the time range of the P3/ RewP. Third, a similar pattern of correlations occurred for the *PE_BIAS_* term (Cluster 1: negative, 111–184 ms, *p* = .004; Cluster 2: positive, 346–449 ms, *p* < .001; Supplementary Fig. 14A). Fourth, around these same time windows, midfrontal voltage also encoded outcome valence itself, but with opposite sign (Cluster 1: positive, 99–184 ms, *p* < .001; Cluster 2: negative, 308–448 ms, *p* < .001; see Supplementary Note 11 and Supplementary Fig. 12A).

In sum, similar to analyses of midfrontal power reported in the main text, PE sign and magnitude were encoded in midfrontal voltage around the same time, but with opposite polarity: Signal around the time of the N1 encoded PE sign positively, but PE magnitude negatively. Vice versa, signal around the time of the P3/ RewP encoded PE sign negatively, but PE magnitude positively. The same phenomenon of separate valence and magnitude encoding in midfrontal EEG signal has been reported before ^12,13,19^. Notably, magnitude encoding in midfrontal voltage emerged for the *PE_BIAS_* term, but not the *PE_STD_*, indicating that this correlation was driven by the *PE_DIF_* term and that biased learning described midfrontal voltage better than standard learning. These results complement our findings of theta/delta power encoding outcome valence and magnitude with opposite polarities (see main text).

## Supplementary Note 13: fMRI-informed EEG results in time-frequency space

Besides the results for striatum, ACC, and PCC reported in the main text, there were also significant EEG correlates over midfrontal electrodes for trial-by-trial BOLD signal from left motor cortex (*p* = .002, around 0–625 ms, 16–27 Hz; Supplementary Fig. 17A). There were however no significant EEG correlates over midfrontal electrodes for BOLD signal from pgACC (*p* = .174; Fig. Supplementary Fig. 17B), left inferior temporal gyrus (*p* = .097; Supplementary Fig. 17C), and primary visual cortex (*p* = .170; Supplementary Fig. 17D).

As quality checks, we checked whether visual cortex BOLD correlated negatively with alpha over occipital electrodes ^24,25^ and whether motor cortex BOLD correlated negatively with beta power over central electrodes ^26,27^. Both was the case (see Supplementary Fig. 17E, F), showing that our data was of sufficient quality to detect these well-established associations.

### Supplementary Note 14: fMRI-informed EEG results in the time domain

For fMRI-inspired analysis of the EEG signal in the time domain (voltage), we applied the same approach as reported in main text, but with voltage signal (time-domain) instead of time-frequency power as dependent variable. As independent variables, we entered the trial-by-trial BOLD signal from all seven regions encoding biased PEs plus the trial-by-trial standard PE and the different term towards the biased PE (exact same procedure as for EEG TF analyses), all in one single multiple linear regression. On a group-level, we again focused on the mean signal over midfrontal electrodes (Fz/ FCz/ Cz) in a time range of 0–700 ms, for which ERPs had been visible in the condition-averaged plots (see Supplementary Note 11 and Supplementary Fig. 12 and 13).

First, trial-by-trial striatal BOLD correlated significantly with midfrontal voltage at two time points, namely positively around 152–196 ms (*p* = .017) in the time range of the N1 and again negatively around 316–383 ms (*p* < .001, Supplementary Fig. 18A) in the time range of the N2/ FRN and P3/RewP. Second, trial-by-trial pgACC BOLD correlated significantly positively with midfrontal voltage around 347–412 ms (*p* = .006, Supplementary Fig. 18A) in the time range of the N2/ FRN and P3/RewP. Third, trial-by-trial BOLD from primary visual cortex correlated significantly positively with midfrontal voltage around 307–367 ms (*p* = .011, Supplementary Fig. 18B), overlapping with (but slightly earlier than) correlations from pgACC BOLD, i.e., in the time range of the N2/ FRN and P3/RewP. For midfrontal voltage split up per high vs. low BOLD signal (revealing which ERP components were respectively modulated), see Supplementary Fig. 18C-E. There were no significantly correlations between midfrontal voltage and trial-by-trial BOLD from dACC (*p* = .927, Supplementary Fig. 18A), left motor cortex (*p* = .649, Supplementary Fig. 18B), PCC (*p* = .796, Supplementary Fig. 18A), or left inferior temporal gyrus (*p* = .649, Supplementary Fig. 18B). For further details on BOLD-EEG voltage correlations in the time domain, see Supplementary Fig. 18F–L.

Taken together, trial-by-trial BOLD signal in striatum, pgACC, and V1 all correlated with FRN/ RewP amplitude, which was the dominant phenomenon over midfrontal electrodes reflecting outcome valence (see Supplementary Note 11 and Supplementary Fig. 12, 13). Notably, correlations with striatal and pgACC BOLD were of opposite signs, which aligns with the finding that striatal and pgACC BOLD predicted opposite behavioral tendencies on future trials (see main text; see Supplementary Fig. 20). However, crucially, the time domain signal did not allow for a temporal dissociation of these different regions. Possibly, the midfrontal evoked signal (i.e., the part of the signal that was phase-locked to outcome onset) was so stereotyped that only the FRN/ RewP complex showed enough variation across trials to allow for substantial correlations with trial-by-trial BOLD signal. This finding demonstrates that the time-frequency domain signal (i.e., the part of the signal that is not necessarily phase-locked to outcome onset) might be more suited for dissociating the activity of different regions in time.

### Supplementary Note 15: Go/NoGo differences over time in BOLD signal, choices, alpha, and beta power

We observed differences between trials with Go responses and trials with NoGo responses in the low alpha power before and shortly after outcome onset (Fig. 6A, B main text). Alpha typically increases over the time course of an experiment, potentially related to fatigue and decreasing arousal ^28^. If the ratio of Go and NoGo responses changed over time, as well, such an increase over time could spuriously lead to a difference between Go and NoGo responses (though note that this ratio did not noticeably change over time; Supplementary Fig. 19D). To exclude this possibility, we extracted trial-by-trial time-frequency power from the three significant clusters report in the main text in which power differed between Go and NoGo responses: i) lower alpha band power after outcome onset, ii) lower alpha band power before and after outcome onset, iii) beta band power before outcome onset. We log10-transformed this data to decibel and analyzed it as a function of the performed response (factor), block number (1–6; z-standardized), and the interaction between both. We reasoned that if power differences occurred merely due to fatigue effects, the main effect of performed response should not be significant when accounting for time on task (i.e., block number).

For lower alpha band power after outcome onset, there was a significant main effect of performed response, *b* = 0.035, *SE* = 0.015, χ^2^(1) = 5.350, *p* = .021, with higher power for Go than NoGo responses, a significant main effect of block number with lower alpha band power increasing over time, *b* = 0.052, *SE* = 0.019, χ^2^(1) = 6.645, *p* = .010, but no significant interaction, *b* = 0.003, *SE* = 0.008, χ^2^(1) = 0.156, *p* = .693. As Supplementary Fig. 19A reveals, lower alpha band power was consistently higher after Go than after NoGo responses for every block of the task, suggesting that differences in lower alpha band power were not merely due to time on task.

For lower alpha band power before and after outcome onset, as well, there was a significant main effect of performed response, *b* = 0.068, *SE* = 0.030, χ^2^(1) = 5.010, *p* = .025, with higher power after Go than NoGo responses, a significant main effect of block number with lower alpha band power increasing over time, *b* = 0.072, *SE* = 0.029, χ^2^(1) = 6.757, *p* = .016, but no significant interaction, *b* = 0.010, *SE* = 0.009, χ^2^(1) = 1.184, *p* = .277 (Supplementary Fig. 19B), leading to identical conclusions.

For beta band power before and after outcome onset, there was a significant main effect of performed response, *b* = 0.083, *SE* = 0.032, χ^2^(1) = 6.301, *p* = .012, with higher power after Go than NoGo responses, a significant main effect of block number with beta power decreasing over time, *b* = − 0.042, *SE* = 0.021, χ^2^(1) = 4.007, *p* = .045, but no significant interaction, *b* = 0.001, *SE* = 0.007, χ^2^(1) = 0.030, *p* = .864 (Supplementary Fig. 19C). In sum, even in presence of changes in power over the time course of the task, lower alpha band and beta band power were consistently higher after Go responses than after NoGo responses, suggesting that these effects were not due to time on task.

Furthermore, we asked whether differences in dACC BOLD between trials with Go and trials with NoGo response at the time of the outcome were due to outcome-related activity or might rather the reflect action on the next trial. We thus plotted the “raw” BOLD signal per action x outcome condition. We used the first eigenvariate of the BOLD in signal in the dACC cluster that reflected biased learning, upsampled the BOLD signal, epoched it into trials relative to outcome onset (same procedure as for fMRI-informed EEG analyses), and averaged the signal across trials and participants separately per performed action (Go/NoGo) and outcome valence (positive/ negative). This plot yielded higher dACC BOLD signal on trials with NoGo responses than on trials with Go responses at the time of outcomes (Supplementary Fig. 19E). However, this difference could potentially be driven by the response on the following task. Hence, we further split the data according to whether the action on the following trial was a Go or a NoGo response. Irrespective of the action on the following trial, dACC BOLD signal was higher when the action on the current trial was a NoGo response compared to a Go response (Supplementary Fig. 20F). In sum, these analyses corroborate that dACC BOLD signal was indeed higher after NoGo than Go responses at the time of outcomes.

**Supplementary Figure 1.**
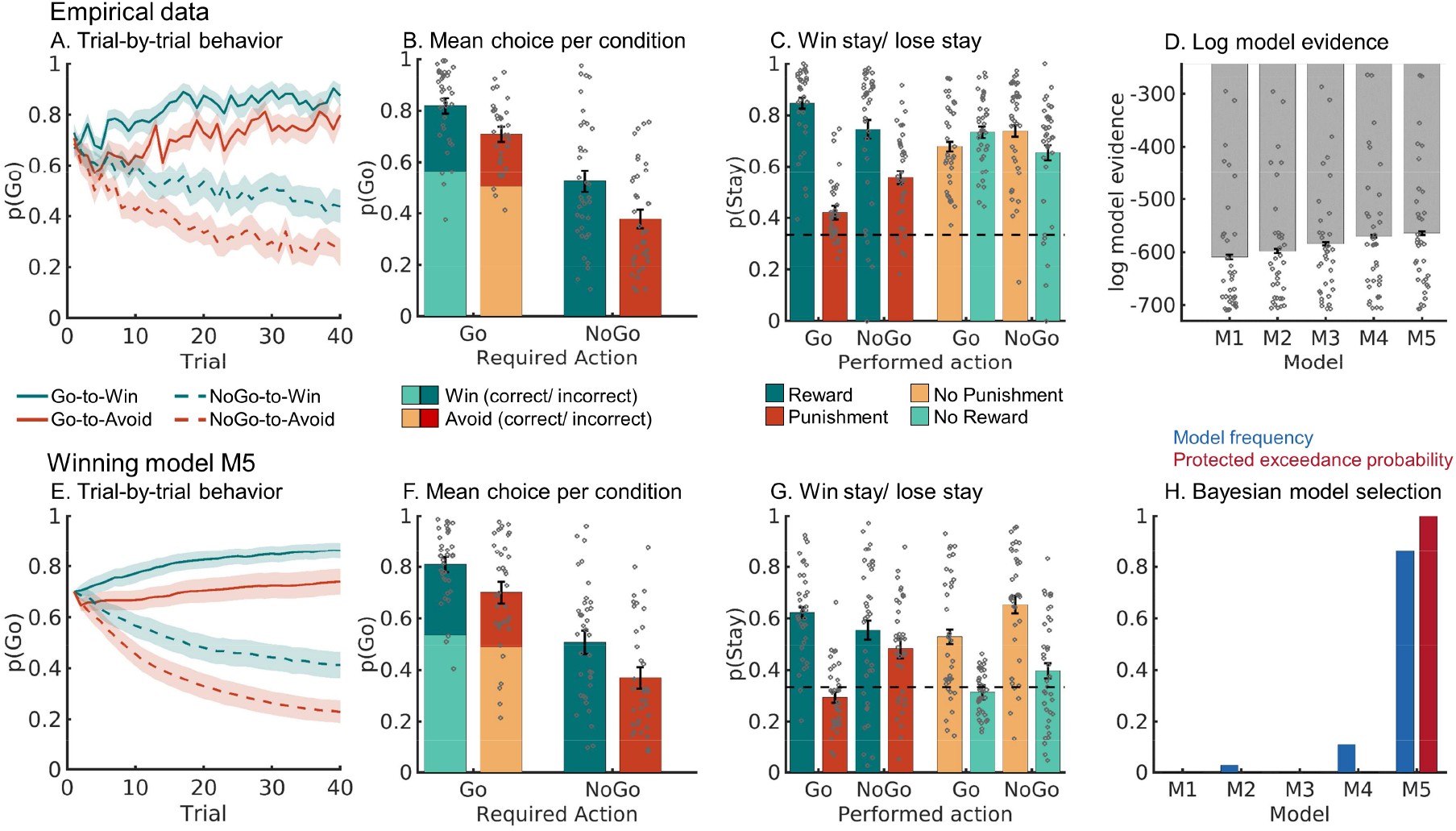
Behavioral performance in the subgroup of 29 participants included in the fMRI-inspired EEG analyses. **A.** Trial-by-trial proportion of Go responses (±SEM across participants) for Go cues (solid lines) and NoGo cues (dashed lines). The motivational bias was already present from very early trials onwards, as participants made more Go responses to Win than Avoid cues (i.e., green lines are above red lines). Additionally, participants clearly learn whether to make a Go response or not (proportion of Go responses increases for Go cues and decreases for NoGo cues). **B.** Mean (±SEM across participants) proportion Go responses per cue condition (points are individual participants’ means). **C.** Probability of repeating a response (“stay”) on the next encounter of the same cue as a function of action and outcome. Learning was reflected in higher probability of staying after positive outcomes than after negative outcomes (main effect of outcome valence). Biased learning was evident in learning from salient outcomes, where this valence effect was stronger after Go responses than NoGo responses. Dashed line indicates chance level choice (p_Stay_ = 0.33). **D.** Log-model evidence favors the asymmetric pathways model (M5 over simpler models (M1-M4). **E-G.** Trial-by-trial proportion of Go responses, mean proportion Go responses, and probability of staying based on one-step-ahead predictions using parameters (hierarchical Bayesian inference) of the winning model (asymmetric pathways model, M5). **H.** Model frequency and protected exceedance probability indicate best fit for model M5 (asymmetric pathways model), in line with log model evidence.

**Supplementary Figure 2.**
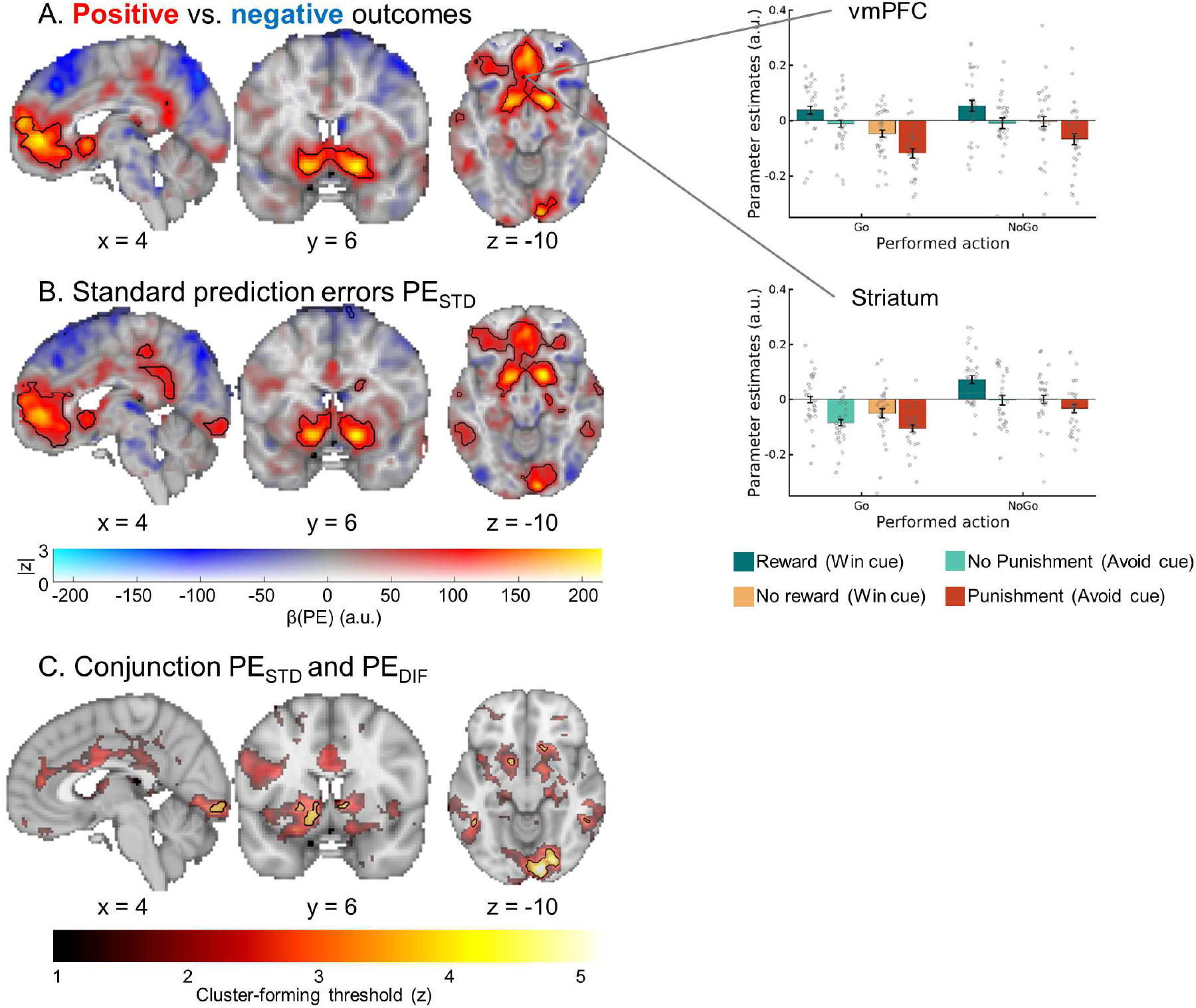
BOLD signal reflecting outcome processing in the subgroup of 29 participants included in the fMRI-inspired EEG analyses. **A.** BOLD signal was higher for positive outcomes (rewards, no punishments) compared with negative outcomes (no rewards, punishments) in a range of regions including bilateral ventral striatum and vmPFC. BOLD effects displayed using a dual-coding data visualization approach with color indicating the parameter estimates and opacity the associated z-statistics. Significant clusters are surrounded by black edges. Bar plots show parameter estimates per action x outcome condition (±SEM across participants) **B.** When using the trial-by-trial PEs participants experienced as model-based regressors in our GLM, positive PE correlations occurred in several regions including importantly the ventral striatum, vmPFC, dACC, and PCC. **C.** Left panel: Regions encoding both the standard PE term and the difference term to biased PEs (conjunction) at different cluster-forming thresholds (color). Clusters significant at a threshold of z > 3.1 are surrounded by black edges. In bilateral striatum, pgACC, bilateral ITG, and primary visual cortex, BOLD was significantly better explained by biased learning than by standard learning. Clusters in dACC, left motor cortex, and PCC were not significant any more.

**Supplementary Figure 3.**
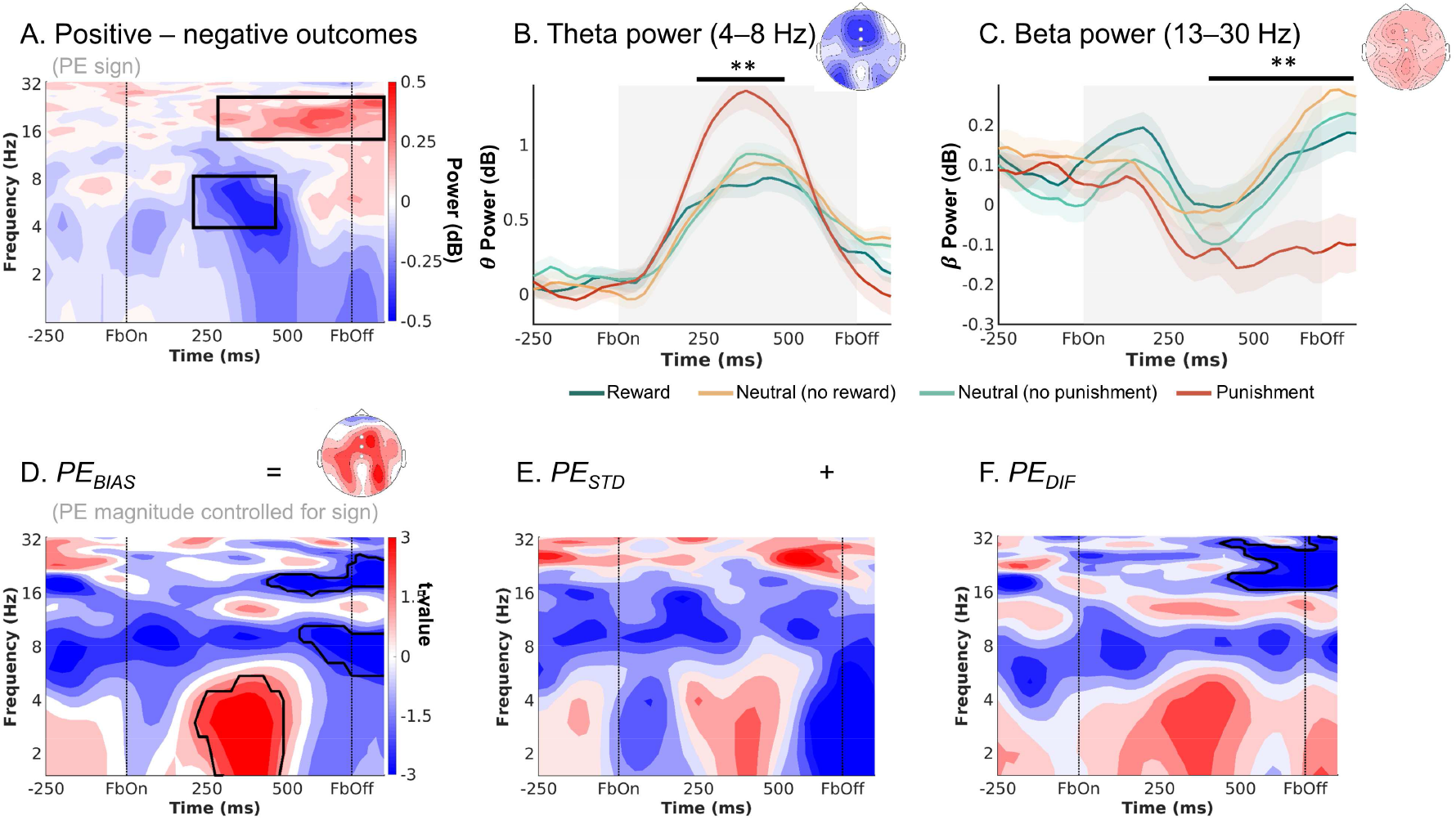
EEG time-frequency power midfrontal electrodes (Fz/ FCz/ Cz) reflecting outcomes processing in the subgroup of 29 participants included in the fMRI-inspired EEG analyses. **A.** Time-frequency plot (logarithmic y-axis) displaying high theta (4–8 Hz) power for negative outcomes and higher beta power (16–32 Hz) for positive outcomes. **B.** Theta power transiently increases for any outcome, but more so for negative outcomes (especially punishments) around 225–475 ms after feedback onset. **C.** Beta was higher for positive than negative outcomes (especially punishments) over a long time period around 300–1,250 ms after feedback onset. **D-F.** Correlations between midfrontal EEG power and trial-by-trial PEs. Solid black lines indicate clusters above threshold. Biased PEs were significantly positively correlated with midfrontal theta power, but also negatively correlated with later alpha and beta power (**D**). The correlations of theta with the standard PEs (**E**) and the difference term to biased PEs (**F**) were also positive, though not significant. Beta power only encoded the difference term to biased PEs (**F**). ** p < 0.01.** *p* < 0.01.

**Supplementary Figure 4.**
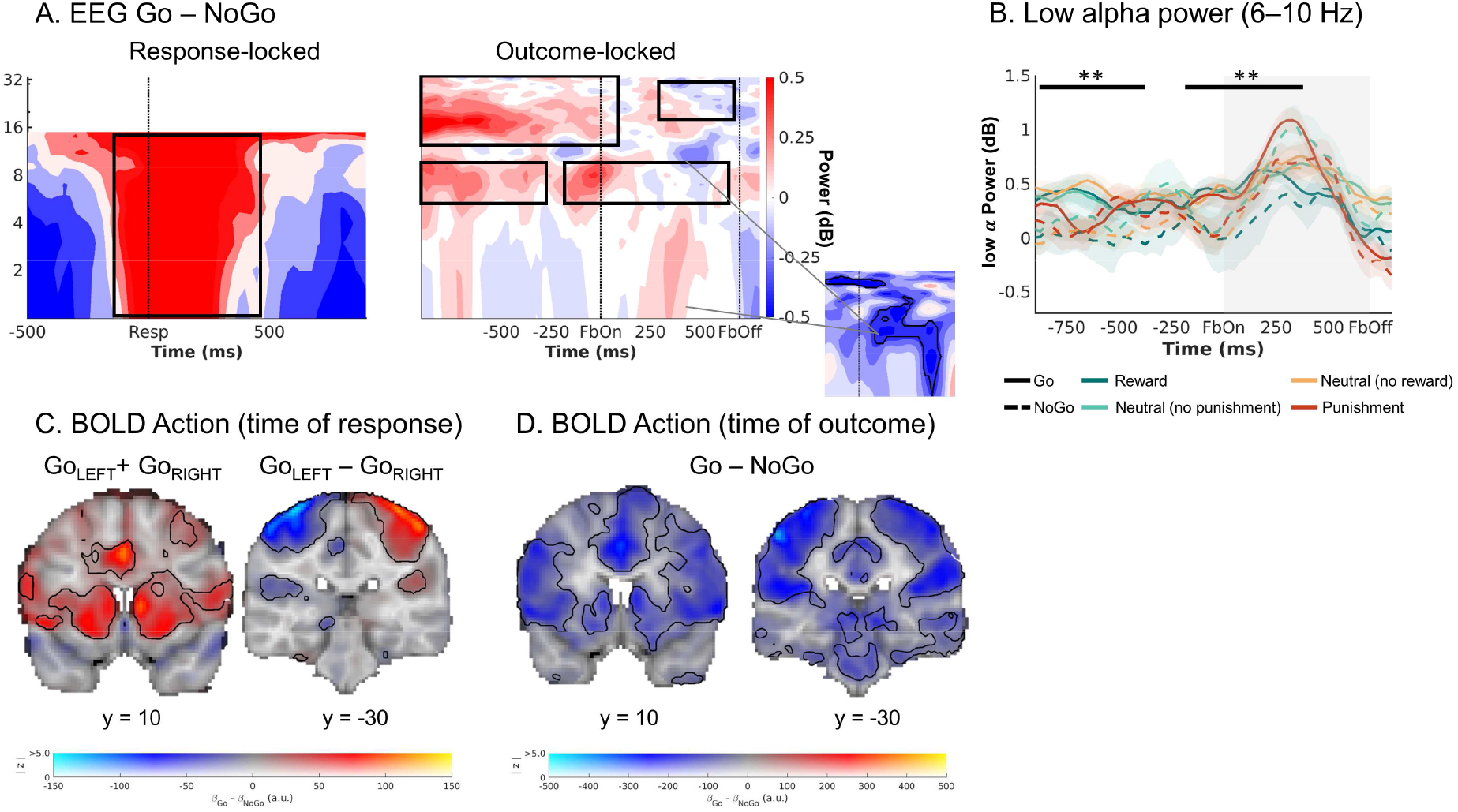
Exploratory follow-up analyses on dACC BOLD signal and midfrontal low-alpha power in the subgroup of 29 participants included in the fMRI-inspired EEG analyses. **A.** Midfrontal time-frequency response-locked (left panel) and outcome-locked (right panel). Before and shortly after outcome onset, power in the lower alpha band was higher on trials with Go actions than on trials with NoGo actions. The shape of this difference resembles the shape of dACC BOLD-EEG TF correlations (small plot; note that this plot depicts BOLD-EEG correlations, which were negative). Note that differences between Go and NoGo trials occurred already before outcome onset in the alpha and beta range, reminiscent of delay activity; but were not fully sustained since the actual response. **B.** Midfrontal power in the lower alpha band per action x outcome condition. Lower alpha band power was consistently higher on trials with Go actions than on trials with NoGo actions, starting already before outcome onset. **C.** BOLD signal differences between Go and NoGo actions (activation by either left or right Go actions compared to the implicit baseline in the GLM, which contains the NoGo actions; left panel) and left vs. right hand responses (right panel) at the time or responses. Response-locked dACC BOLD was significantly higher for Go than NoGo actions. **D.** BOLD signal differences between Go and NoGo actions at the time of outcomes. Outcome-locked dACC BOLD signal (and BOLD signal in other parts of cortex) was significantly lower on trials with Go than on trials with NoGo actions.

**Supplementary Figure 5.**
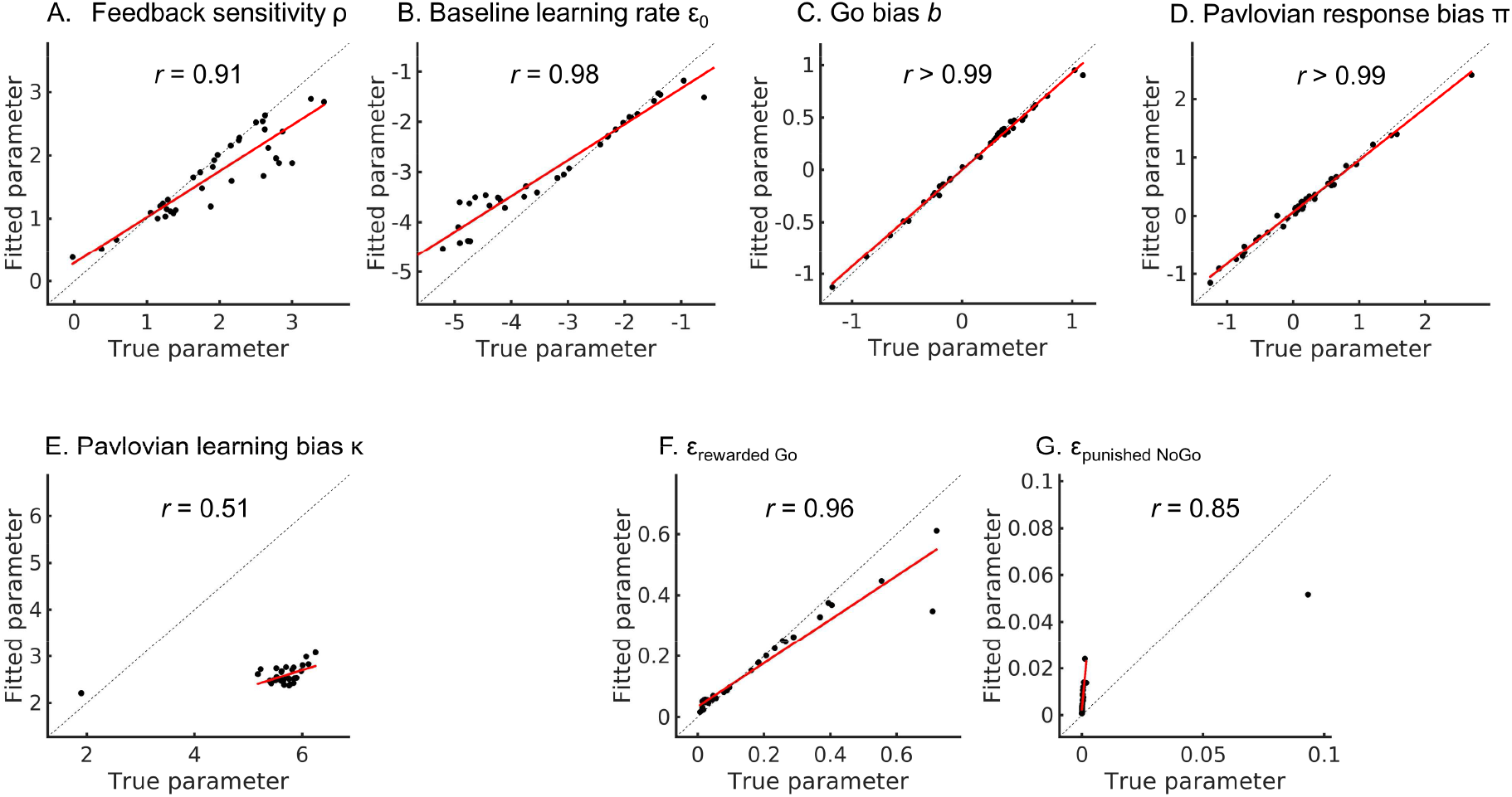
Parameter recovery results for the asymmetric pathways (M5) model. The feedback sensitivity parameter ρ **(A**), the baseline learning rate ε_0_ (**B**), the Go bias *b* (**C**), and the Pavlovian response bias π (**D**) all showed excellent parameter recovery, i.e., between-participants correlations of ground-truth and fitted parameters all exceeded *r* > 0.90. Parameters ρ and ε_0_ are still in sampling space and thus untransformed (which means they can be negative). Dashed lines represent the identity line; red solid lines represent a linear regression line of fitted parameters regressed onto true parameters. Only recovery of the learning bias parameter κ (**E**) was not quite as good, though the correlation between ground-truth and fitted parameters was still strongly positive (r > 0.50). Note an outlier at the bottom left of κ values; the regression line was fitted without this data point. When combining the baseline learning rate ε_0_ with the learning bias κ to compute the biased learning rates for rewarded Go actions ε_rewarded Go_ (**F**) and punished NoGo actions ε_punished NoGo_ (**G**), correlations between ground-truth and fitted parameter values were considerably higher (*r*’s > 0.86). Note again an outlier at the top right of for ε_punished NoGo_ values; the regression line was fitted without this data point.

**Supplementary Figure 6.**
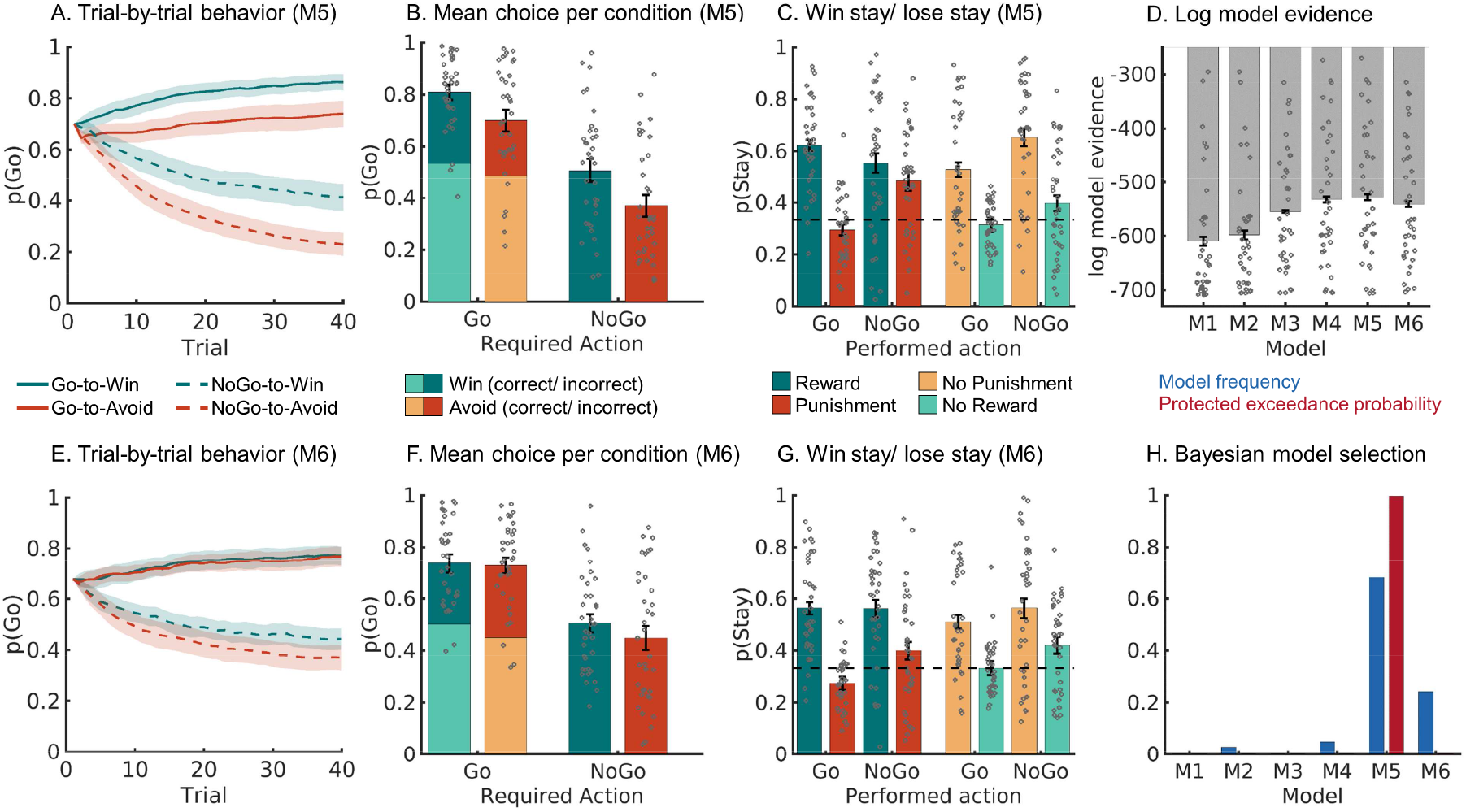
Model comparison and validation of asymmetric pathways (M5) and action priming (M6) model. **(A-C**) One-step-ahead predictions using parameters (hierarchical Bayesian inference) of the winning model asymmetric pathways model (M5). **A.** Trial-by-trial proportion of Go responses (±SEM across participants) for Go cues (solid lines) and NoGo cues (dashed lines); **B.** Mean (±SEM across participants) proportion Go responses per cue condition (points are individual participants’ means); **C.** Probability of repeating a response (“stay”) on the next encounter of the same cue as a function of action and outcome. The asymmetric pathways model was well able to capture core characteristics of the empirical data (see Fig. 2 in the main text). **D.** Log-model evidence favors the asymmetric pathways model (M5), even over the action priming model (M6). **E-G.** Trial-by-trial proportion of Go responses, mean proportion Go responses, and probability of for the action priming model (M6). This model did not reproduce motivational biases (i.e., the difference between green and red lines and bars) well. **H.** Model frequency and protected exceedance probability indicate best fit for model M5 (asymmetric pathways model), in line with log model evidence.

**Supplementary Figure 7.**
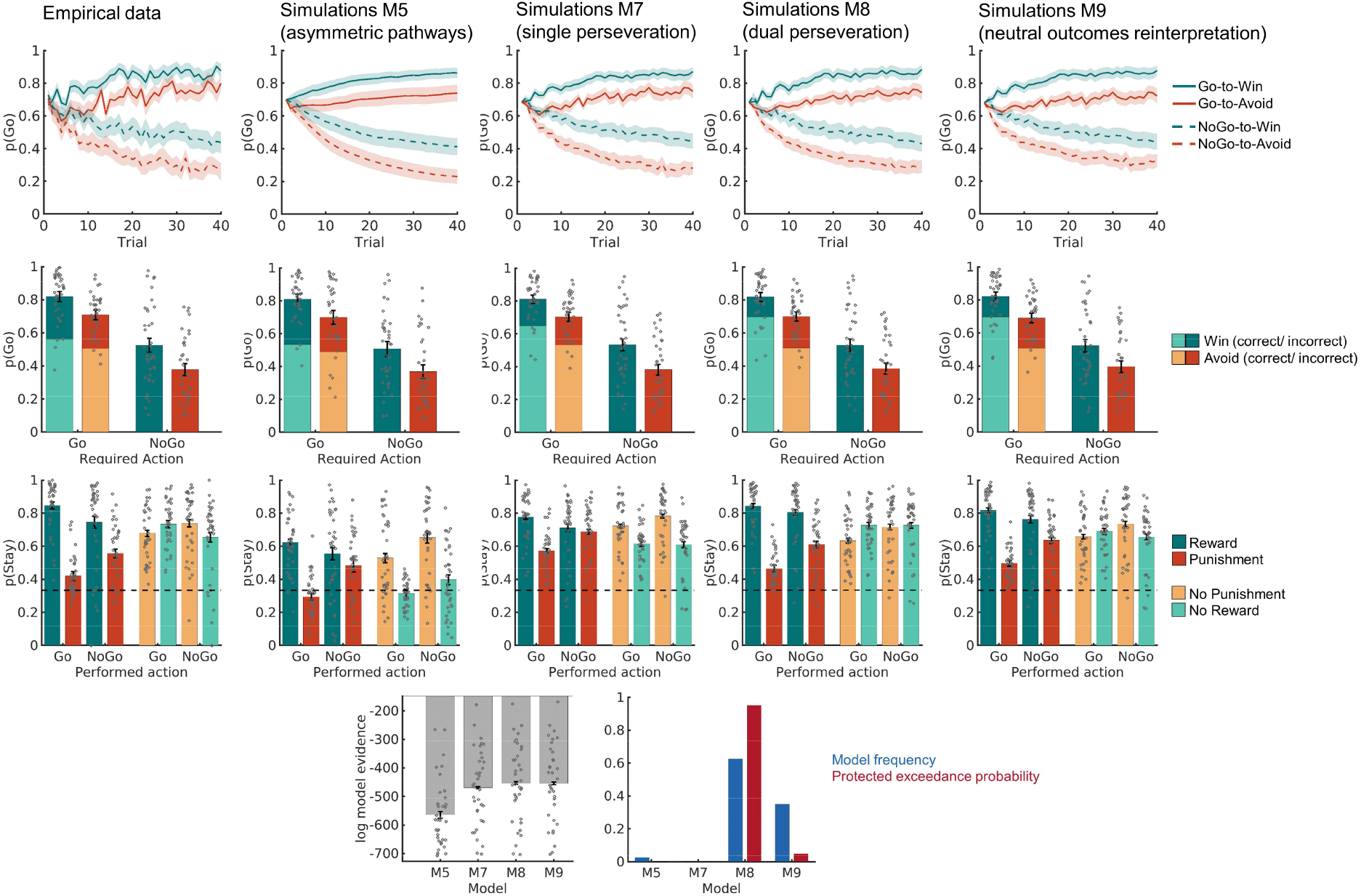
Model comparison and validation of the single perseveration (M7), dual perseveration (M8) and cue valence-based outcome reinterpretation models. **First row.** Trial-by-trial proportion of Go responses (±SEM across participants) for Go cues (solid lines) and NoGo cues (dashed lines). **Second row.** Mean (±SEM across participants) proportion Go responses per cue condition (points are individual participants’ means). **Third row.** Probability to repeat a response (“stay”) on the next encounter of the same cue as a function of action and outcome. **Fourth row.** Log-model evidence, model frequency, and protected exceedance probability all favored the dual perseveration model (M8) over the other models. In sum, the additional models M7-9 provided a better quantitative fit to the data compared to the asymmetric pathways model M5 reported in the main text. They also predicted the propensity of staying overall more accurately than M5. However, these additional models all overestimated the proportion of incorrect Go responses. Furthermore, although the predicted patterns of the propensity of staying mimicked the data more closely than M5, these predicted patterns still mismatched some aspects of the empirical data. Taken together, these models could capture certain qualitative patterns in the data, but not others, which was expectable given the data reduction that comes with fitting a learning model with few parameters only.

**Supplementary Figure 8.**
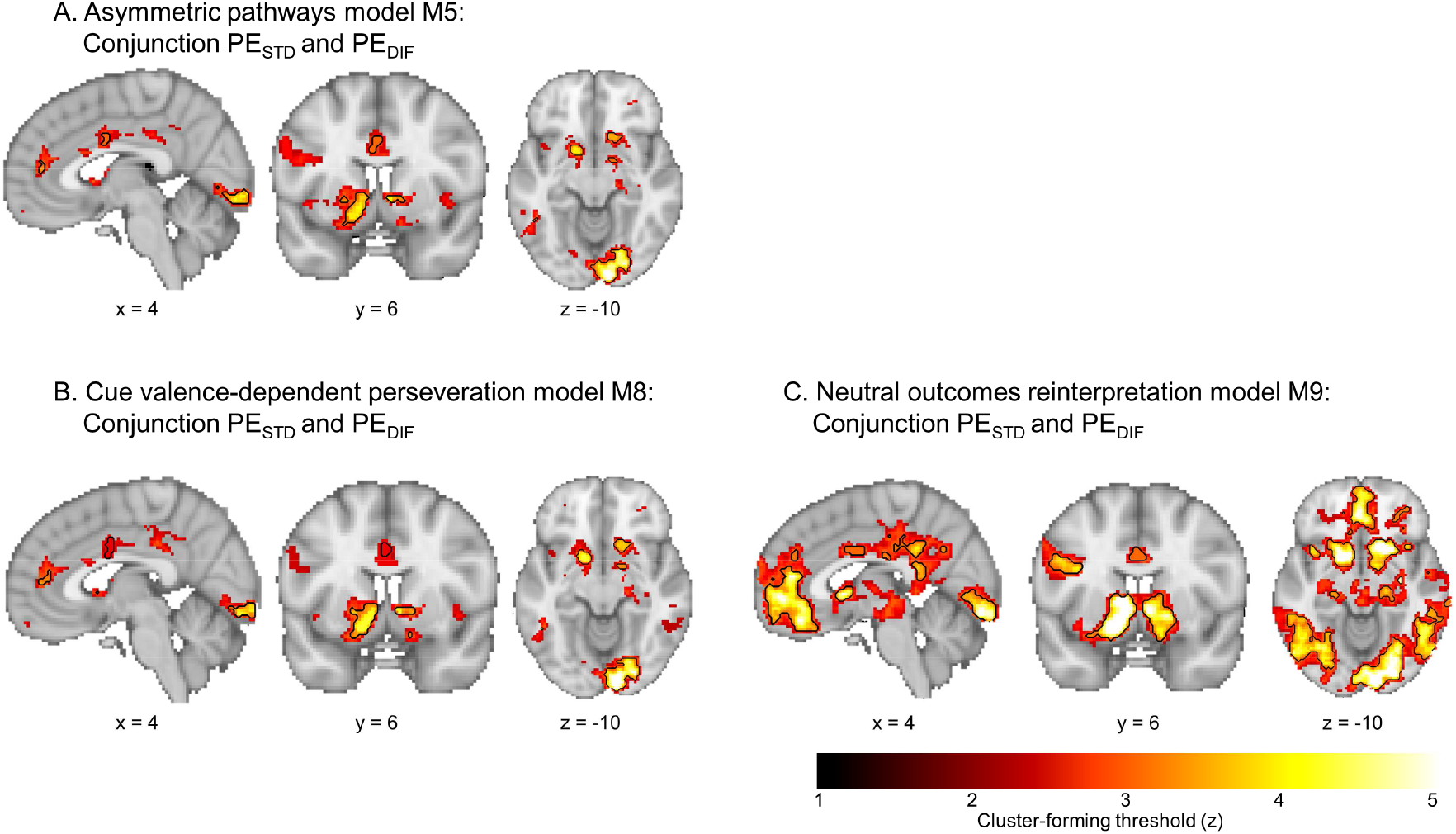
BOLD correlates of biased prediction errors as predicted by the asymmetric pathways model (M5), the cue valence-dependent perseveration model (M8) and the neutral outcomes reinterpretation model (M9). (A) Regions encoding both the standard PE term and the difference term to biased PEs (conjunction) as predicted from the asymmetric pathways model (M5) at different cluster-forming thresholds (1 < z < 5, color coding; opacity constant; replotted from Fig. 3C main text). Clusters significant at a threshold of z > 3.1 are surrounded by black edges. This is a version of Fig. 3C reprinted with a color scheme consistent with the other two panels. (B) Regions encoding both the standard PE term and the difference term to biased PEs (conjunction) as predicted from the cue valence-dependent perseveration model (M8) at different cluster-forming thresholds (1 < z < 5, color coding; opacity constant). Clusters significant at a threshold of z > 3.1 are surrounded by black edges. In line with correlates of biased PEs as predicted by M5, BOLD signal in bilateral striatum, dACC (small-volume corrected), pgACC, PCC, left motor cortex, left inferior temporal gyrus, and primary visual cortex was significantly better explained by biased learning than by standard learning. This finding was not surprising given that adding perseveration to the model did not change the learning mechanism, but only led to slightly different best fitting parameter values. (B) Regions encoding both the standard PE term and the difference term to biased PEs (conjunction) as predicted from the neutral outcomes reinterpretation model (M9). In addition to the regions in which BOLD signal was significantly better explained by biased than standard PEs as derived from M5 and M8, biased PEs derived from M9 also explained BOLD signal in vmPFC (larger cluster than M5), PCC (larger cluster than M5), left inferior frontal gyrus and multiple clusters in superior and inferior lateral occipital cortex significantly better than standard PEs. These results tentatively suggested that vmPFC, PCC, and these other occipital regions might implement an additional mechanism besides biased learning which encodes the cue valence also at the time of the outcome, biasing the processing of neutral outcomes.

**Supplementary Figure 9.**
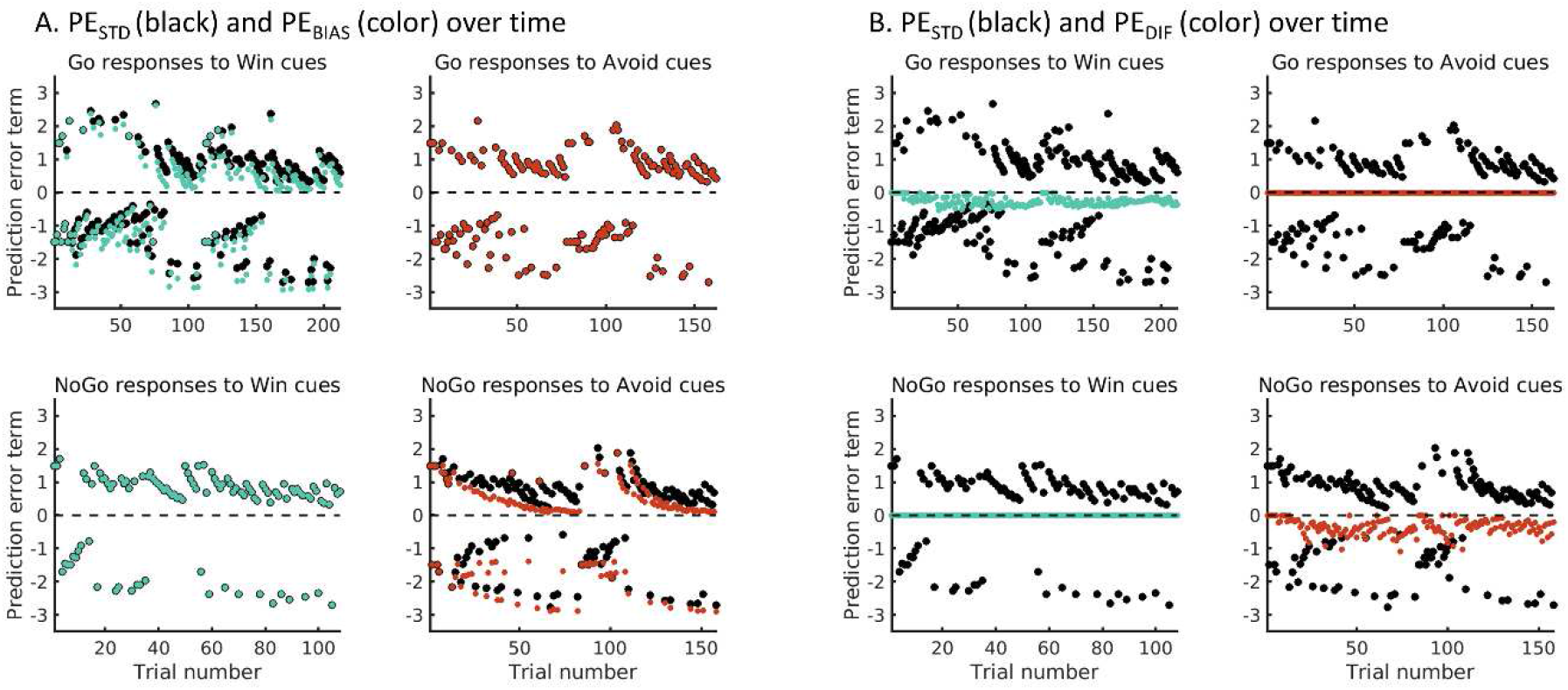
Illustration of biased and standard learning for a representative example participant. **(A)** Prediction errors according to the standard Q-learning model M1 (*PE_STD_*; black dots) and according to the winning model M5 implementing biased learning (*PE_BIAS_*; colored dots). In M5, motivational biases partially arise through biased learning: Participants learn more readily that an action has caused a reward, and are reluctant to learn that inaction has led to a punishment. For each cue, the values of each of the three possible actions (Go_LEFT_, Go_RIGHT_, NoGo) are learnt independently, and prediction errors are calculated relative to the value of the chosen action. The learning bias acts such that the effective learning rate is increased when a reward follows any Go response, and decreased when a punishment follows a NoGo response (see equation 5 in the main manuscript). Hence, for Win cues, action values for Go responses (but not NoGo responses) will be affected by the learning bias and approach the positive asymptote more quickly compared to standard learning, leading to faster decay of positive prediction errors. At the same time, negative outcomes will remain surprising and elicit larger prediction errors compared to standard learning. Hence, model predictions diverge for prediction errors after Go responses to Win cues, but not after NoGo responses to Win cues (colored dots are on top of black dots). Vice versa, for Avoid cues, action values for NoGo responses (but not Go responses) are affected by the learning bias and approach the negative asymptote more slowly compared to standard learning (with negative prediction errors remaining high) as participants are reluctant to take punishments after NoGo responses into account. At the same time, ignoring punishments leads to a faster approach of positive action values to the positive asymptote (and a faster decay of positive prediction errors) compared to standard learning. Model predictions diverge for prediction errors after NoGo responses to Avoid cues, but not after Go responses to Avoid cues (colored dots are on top of black dots). (B) To assess evidence for biased learning despite this high multicollinearity, we decomposed *PE_BIAS_* into *PE_STD_* (black dots) plus a difference term *PE_DIF_ = PE_BIAS_ – PE_STD_* (colored dots). Note that *PE_DIF_* is always zero after NoGo responses to Win cues and Go responses to Avoid cues as both M1 and M5 make identical predictions for these action values. In contrast, for Go responses to Win cues and NoGo responses to Avoid cues, the *PE_DIF_* term is always negative because, in both cases, positive action values approach the positive asymptote more quickly (such that positive prediction errors decay more quickly) compared to standard learning, and negative action values approach the negatively asymptote more slowly (and thus negative prediction errors remain high) compared to standard learning.

**Supplementary Figure 10.**
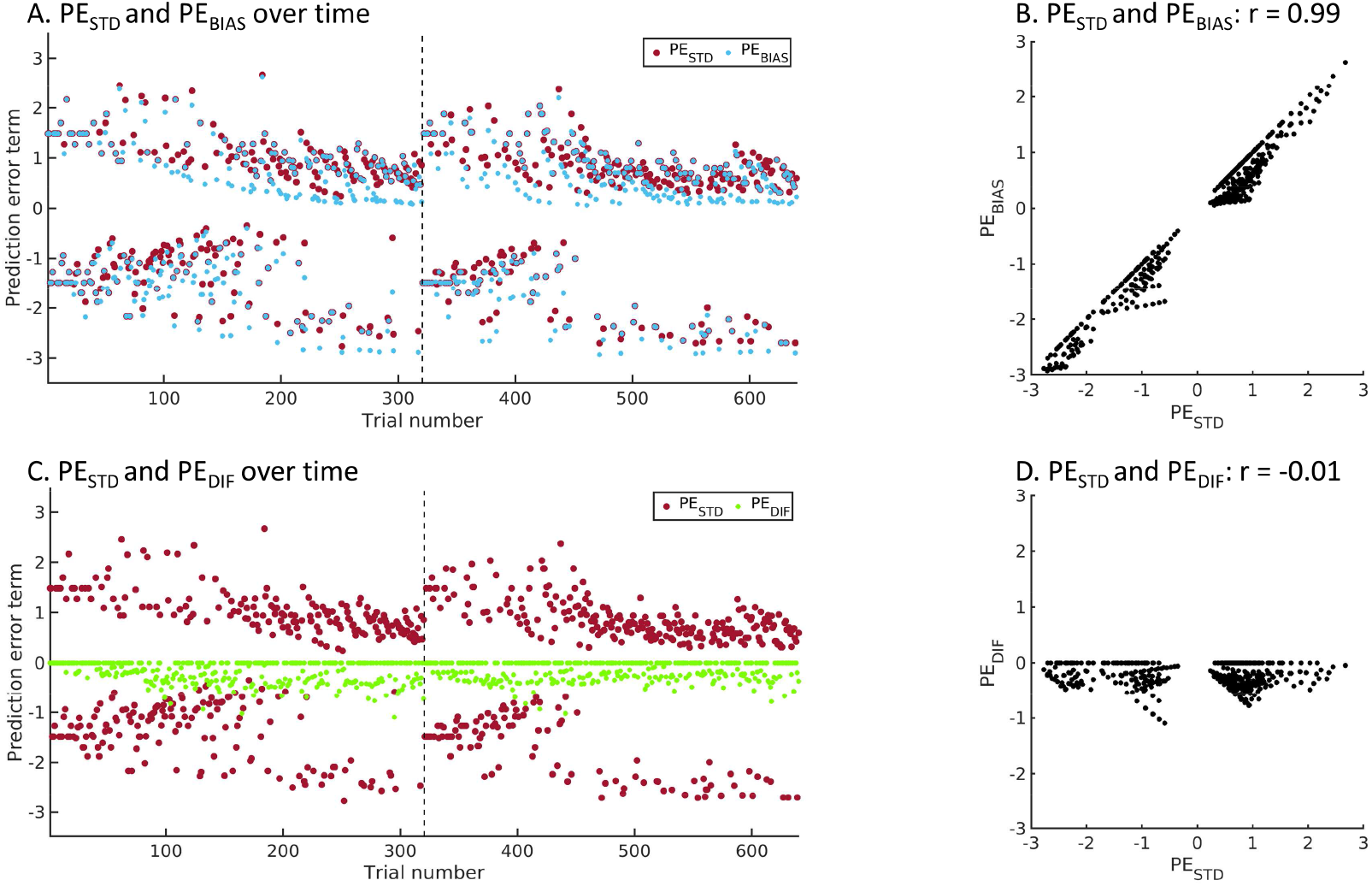
Illustration of prediction error regressor decomposition for a representative example participant. **(A)** Prediction errors according to the standard Q-learning model M1 (*PE_STD_*; larger red dots) and according to the winning model M5 implementing biased learning with more learning from rewarded Go responses and less learning from punished NoGo responses (*PE_BIAS_*; smaller blue dots; blue dots with a red edge reflect trials on which both models make identical predictions). Both prediction error types have a highly similar profile. The key difference between them is an overall downwards shift of *PE_BIAS_* compared to *PE_STD_*, with positive *PE_BIAS_* approaching zero more quickly than positive *PE_STD_*, while negative *PE_BIAS_* remain more negative compared to negative *PE_STD_*. Note that, after trial 320, session 2 starts (vertical dashed line), featuring new cues. **(B)** The prediction errors from both models are highly correlated (mean across participants: *r* = 0.99, range 0.96–0.99), implicating that, when entered together into a multiple linear regression, both regressors would share most of their variance, which would be attributed to neither of them. **(C)** To assess evidence for biased learning despite this high multicollinearity, we decomposed *PE_BIAS_* into *PE_STD_* plus a difference term *PE_DIF_ = PE_BIAS_ – PE_STD_*. *PE_STD_* and *PE_DIF_* show markedly different profiles, with *PE_DIF_* being zero for trials on which both *PE_STD_* and *PE_BIAS_* make identical predictions, and being negative otherwise (reflecting the relatively faster decay of positive PE_BIAS_ and slower decay of negative *PE_BIAS_*). **(D)** Both *PE_STD_* and *PE_DIF_* are much less correlated (mean across participants: *r* = −0.02, range −0.07–0.09), making it possible to enter them in the same multiple linear regression and test whether PE_DIF_ predicts variance in BOLD signal above and beyond *PE_STD_*.

**Supplementary Figure 11.**
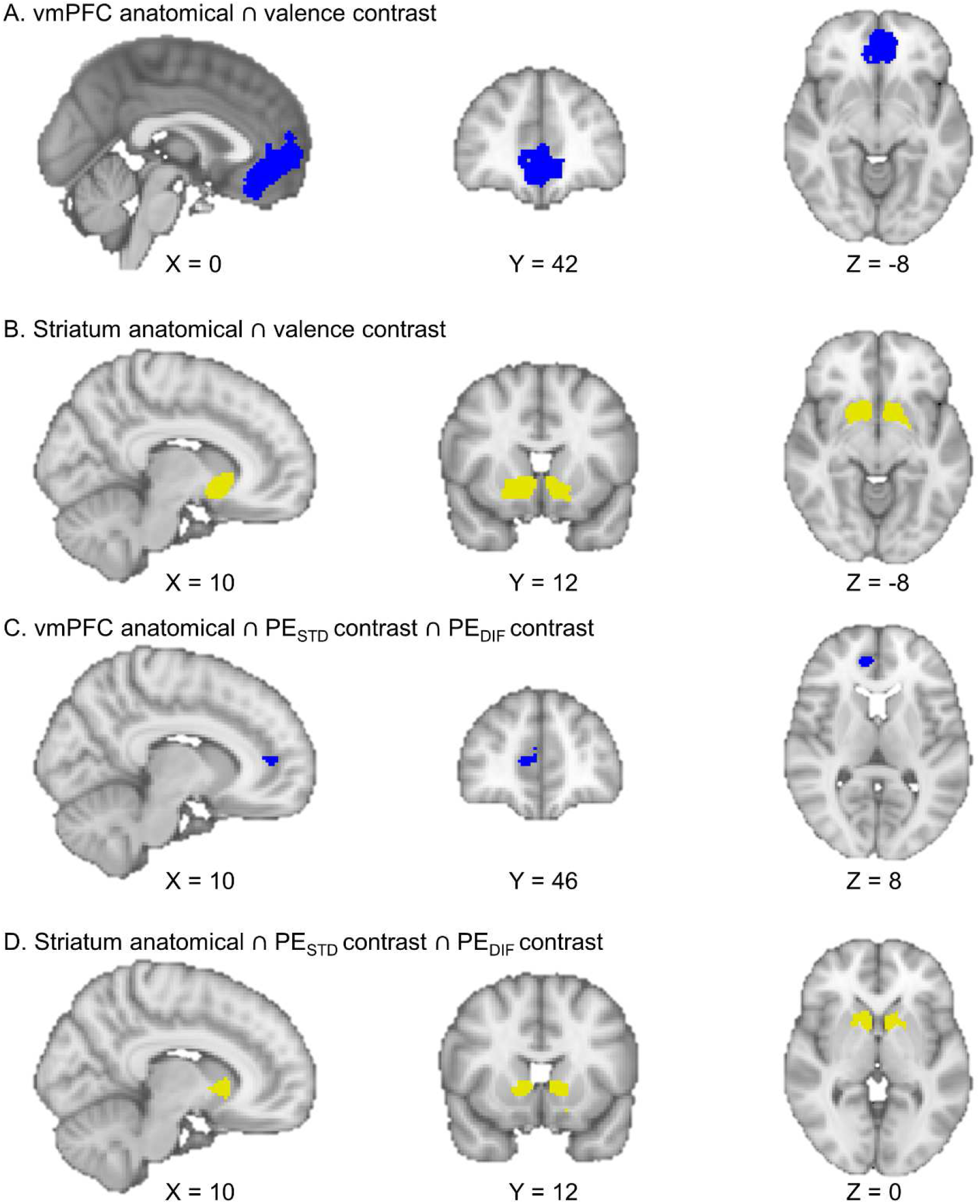
Conjunctions of anatomical masks with functional contrasts from fMRI GLM analyses used for fMRI-informed EEG analyses. Anatomical masks were based on the Harvard-Oxford Atlas. Functional contrasts involve outcome valence and conjunction of PE_STD_ and PE_DIF_. **A**. vmPFC outcome valence contrast (dark blue, conjunction of frontal pole, frontal medial cortex, and paracingulate gyrus). **B.** striatum outcome valence contrast (yellow, conjunction of bilateral nucleus accumbens, caudate, and putamen). **C.** vmPFC PE_STD_ ∩ PE_DIF_ contrast (dark blue, results in a cluster in pgACC). **D.** striatum PE_STD_ *∩* PE_DIF_ contrast (yellow). All anatomical masks were extracted from the probabilistic Harvard-Oxford Atlas, thresholded at 10%. Note that images are in radiological orientation (i.e., left brain hemisphere presented on the right and vice versa).

**Supplementary Figure 12.**
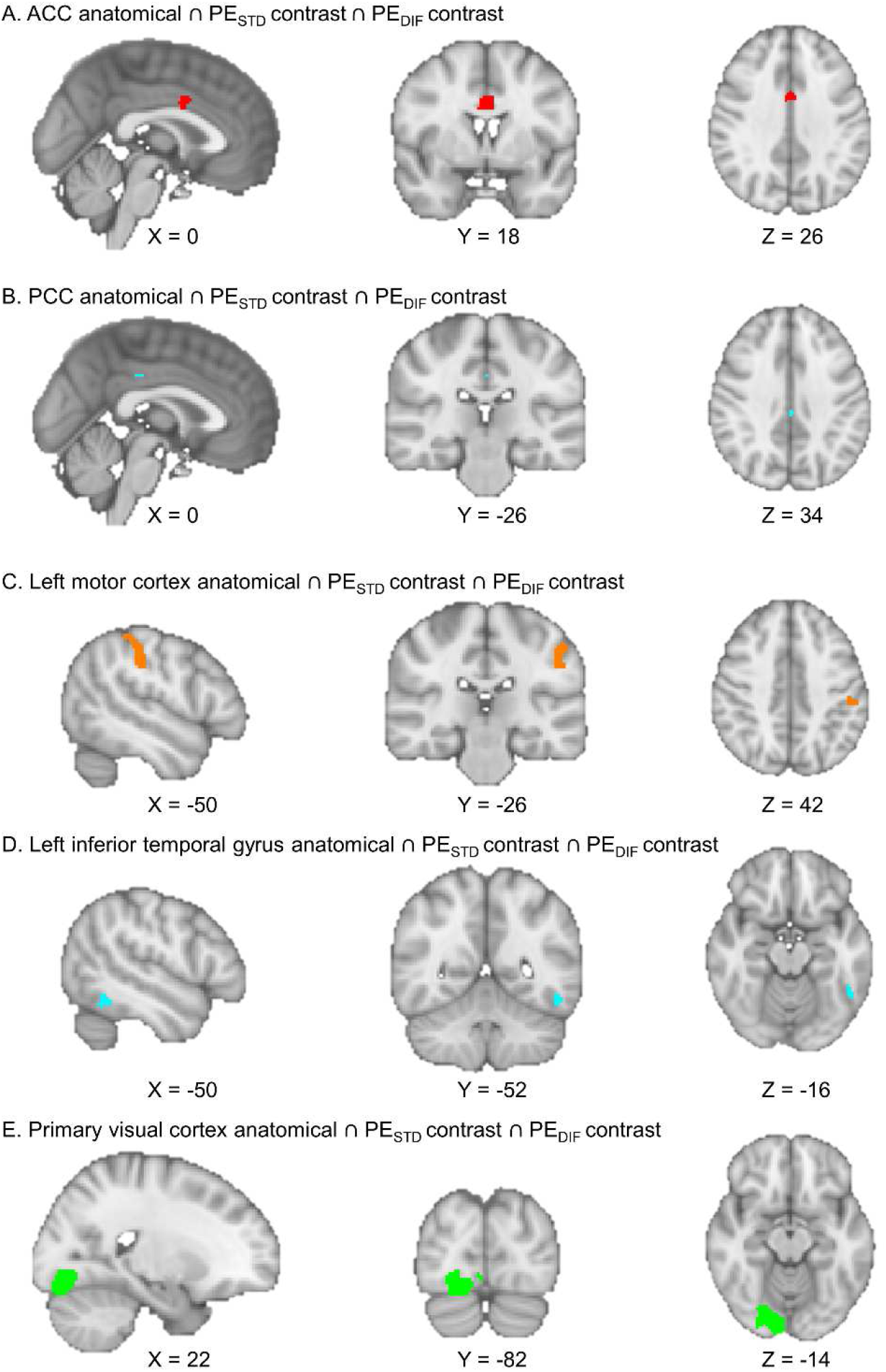
Conjunctions of anatomical masks with functional contrasts from fMRI GLM analyses used for fMRI-informed EEG analyses: **A.** AAC PE_STD_ *∩* PE_DIF_ contrast (red, cingulate gyrus, anterior division, resulting in a cluster in dACC; **B.** PCC PE_STD_ *∩* PE_DIF_ contrast (light blue, cingulate gyrus, posterior division); **C.** Left motor cortex PE_STD_ *∩* PE_DIF_ contrast (orange, conjunction of precentral and postcentral gyrus). **D.** Left inferior temporal gyrus PE_STD_ *∩* PE_DIF_ contrast (turquoise, conjunction of inferior temporal gyrus, posterior division, and inferior temporal gyrus, temporooccipital part). **E.** Primary visual cortex PE_STD_ *∩* PE_DIF_ contrast (green, conjunction of lingual gyrus, occipital fusiform gyrus, occipital pole). All anatomical masks were extracted from the probabilistic Harvard-Oxford Atlas, thresholded at 10%. Note that images are in radiological orientation (i.e., left brain hemisphere presented on the right and vice versa).

**Supplementary Figure 13.**
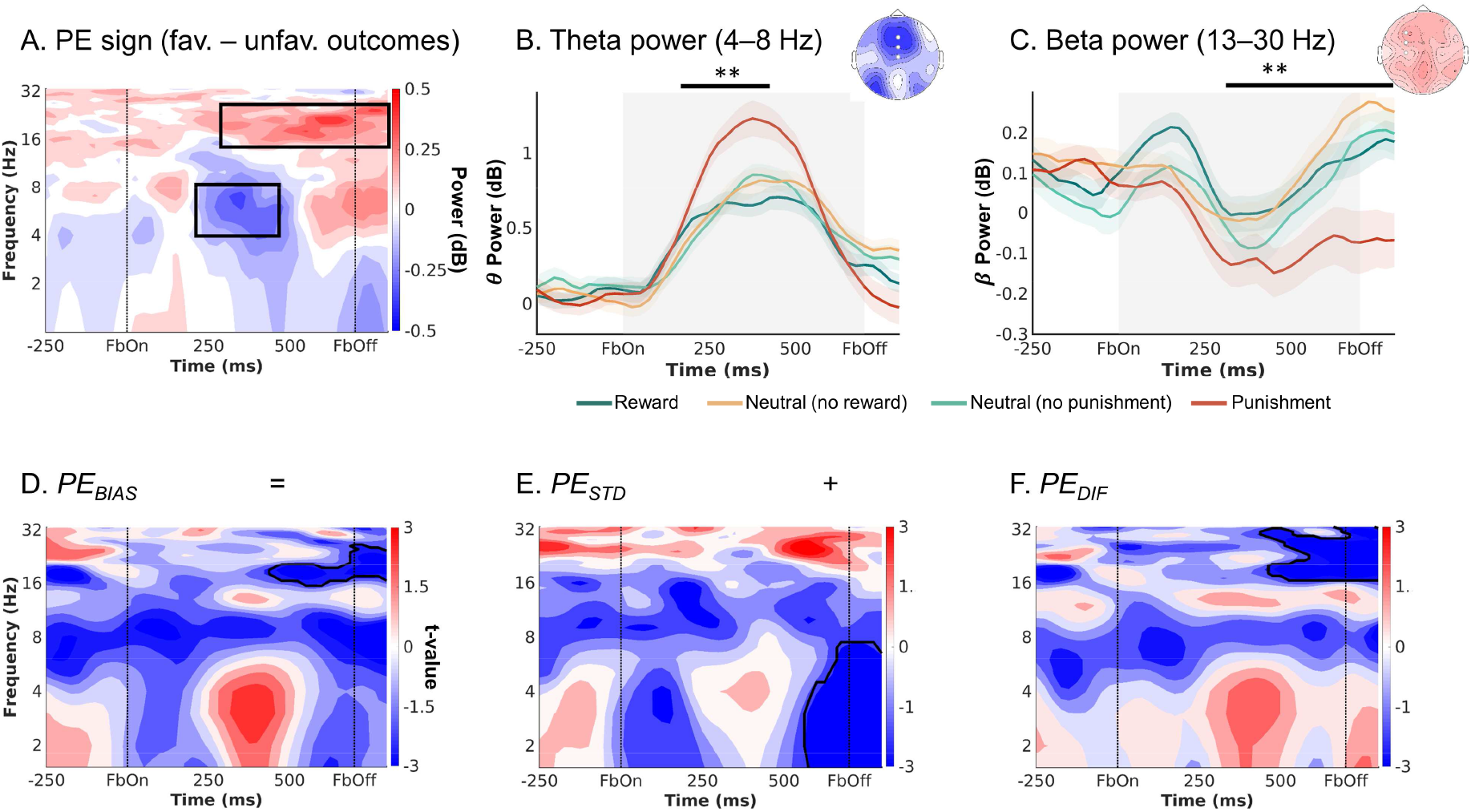
EEG time-frequency power over midfrontal electrodes (Fz/ FCz/ Cz) after the (action x outcome) condition-wise ERPs has been removed. **A.** Time-frequency plot (logarithmic y-axis) displaying high theta (4–8 Hz) power for negative outcomes and higher beta power (16–32 Hz) for positive outcomes. **B.** Theta power transiently increases for any outcome, but more so for negative outcomes (especially punishments) around 225–475 ms after feedback onset. **C.** Beta was higher for positive than negative outcomes (especially punishments) over a long time period around 300–1,250 ms after feedback onset. **D-F.** Correlations between midfrontal EEG power and trial-by-trial PEs. Solid black lines indicate clusters above threshold. There still was a visible positive correlation between biased PEs and midfrontal delta power, but this correlation was not significant (**D**). The correlation of delta with the standard PEs (**E**) was also positive, though not significant; in fact, at a later time point around stimulus offset, delta power correlated significantly negatively with standard PEs. The difference term to biased PEs (**F**) also correlated positively, though not significantly with delta power. Beta power encoded the difference term and biased PEs themselves (F). ** p < 0.01.

**Supplementary Figure 14.**
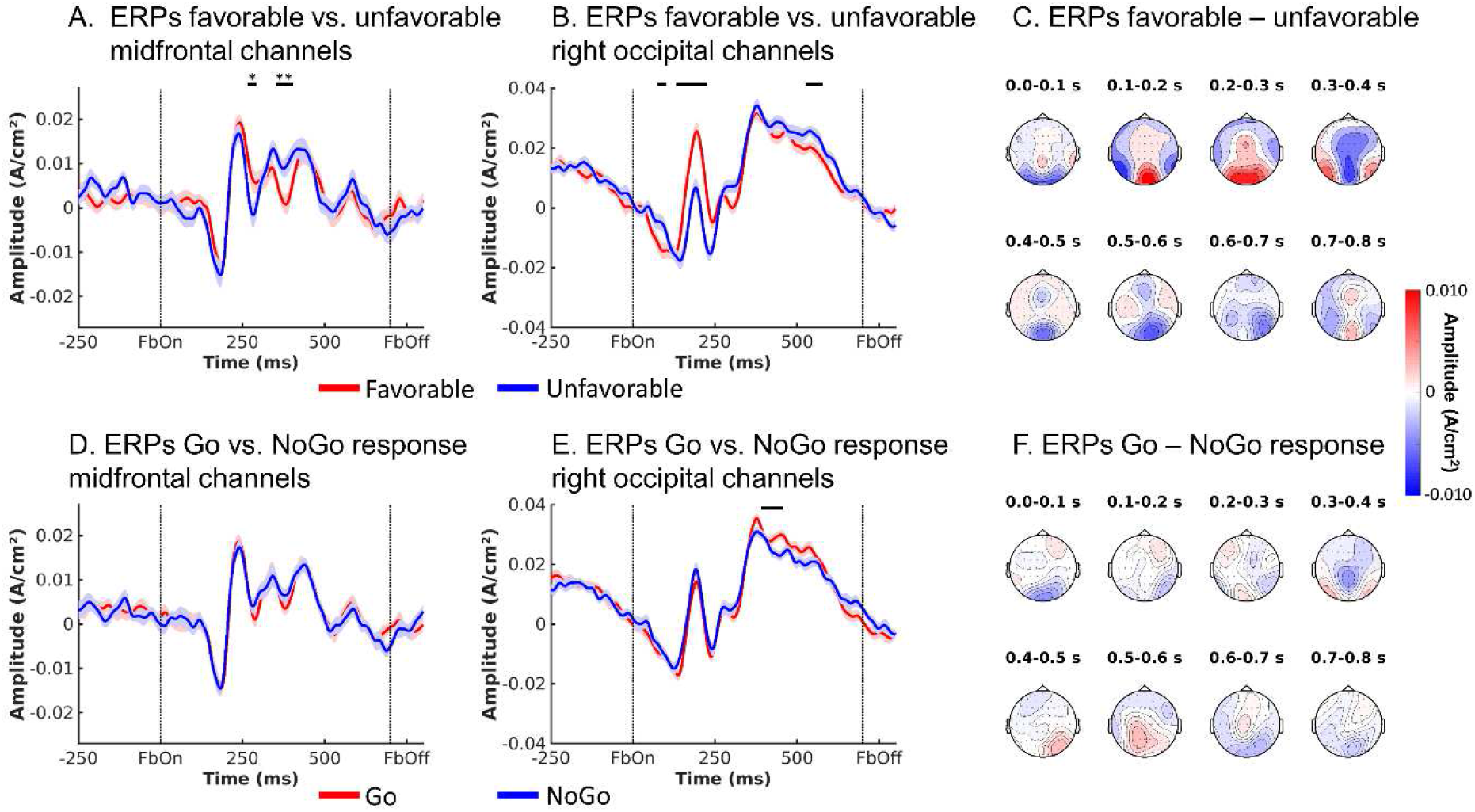
ERPs reflecting outcome valence and performed action. **A.** Voltage (±SEM) over midfrontal electrodes (Fz/FCz/Cz) was lower for negative than positive outcomes around 246–294 ms (stronger N2, FRN) and higher for positive than negative outcomes around 344 – 414 ms (stronger P3/ RewP). **B.** Over right occipital electrodes, the P3 was slightly bigger for positive than negative outcomes. ** *p* < 0.01. * *p* < .05 **C.** Topoplots of difference in voltage between trials with positive and negative outcomes over selected time windows. **D.** There was no difference in voltage over midfrontal electrodes between trials with Go and NoGo responses. **E.** Over right occipital electrodes, the P3 was slightly stronger after Go than NoGo actions (no *p*-value because ROI selected based on visual inspection). **F.** Topoplots of difference in voltage between trials with Go and NoGo actions over selected time windows.

**Supplementary Figure 15.**
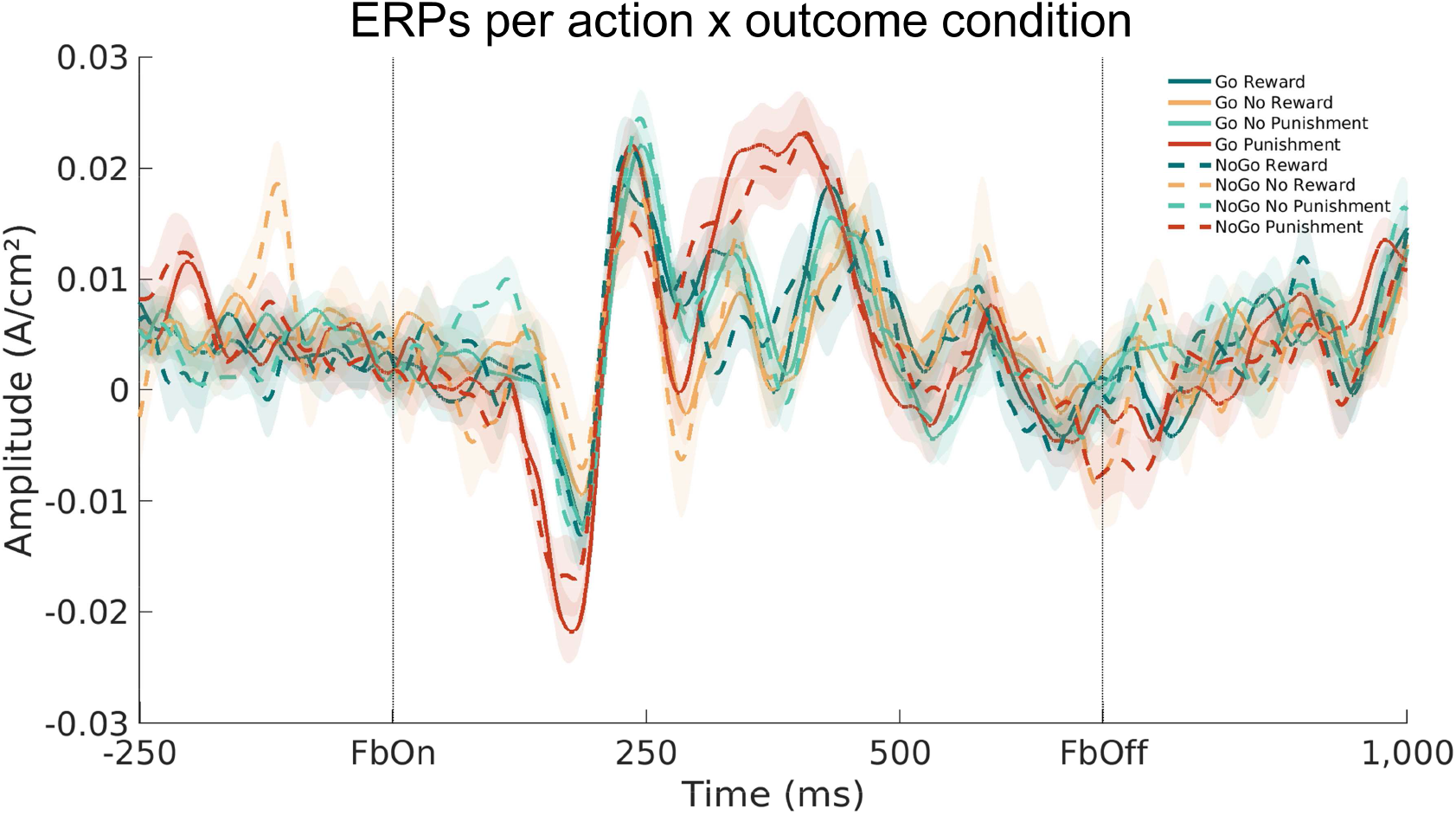
ERPs per action x outcome condition. Biggest differences occurred around the time of the N2 (FRN) and P3 (RewP). N2 and P3 exhibited larger amplitudes on trials with punishments. There was no apparent modulation by the previous action (Go/ NoGo).

**Supplementary Figure 16.**
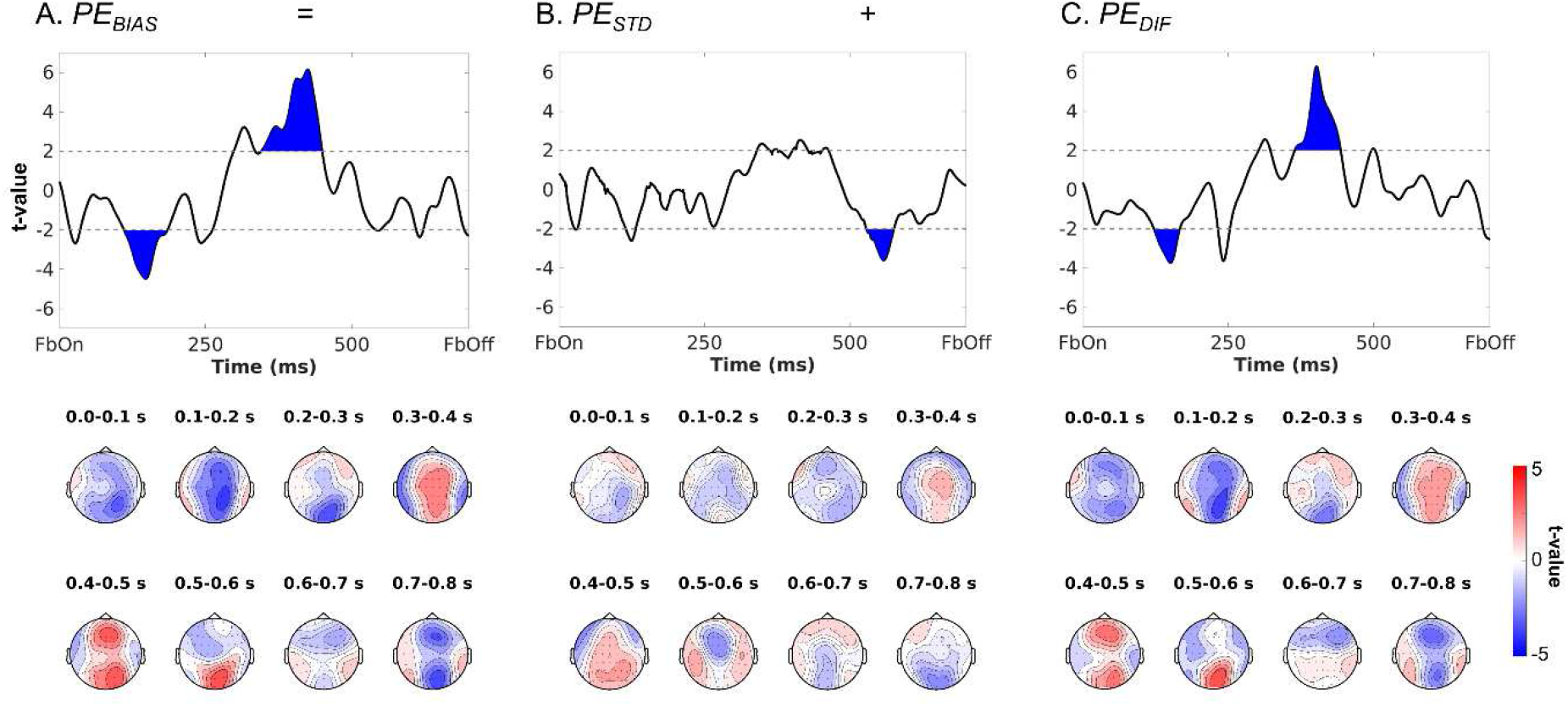
Modulation of EEG voltage by biased PEs and decomposition into the standard PE term and the difference term to biased PEs. **A.** Mean EEG voltage over midfrontal electrodes (Fz, FCz, Cz) was significantly modulated by biased PEs around 111–184 (negatively) and 353–414 ms (positively) after outcome onset. **B.** Correlations with the standard PE term only emerged around 529 – 575 ms (negatively). **C.** Correlations with the difference term to biased PEs were similar to correlations for the biased PE term itself, i.e., around 123–166 (negatively) and 365–443 ms (positively). Bottom row: Topoplots displaying *t*-values of beta-weights for the respective regressor over the entire scalp in steps of 100 ms from 0 to 800 ms.

**Supplementary Figure 17.**
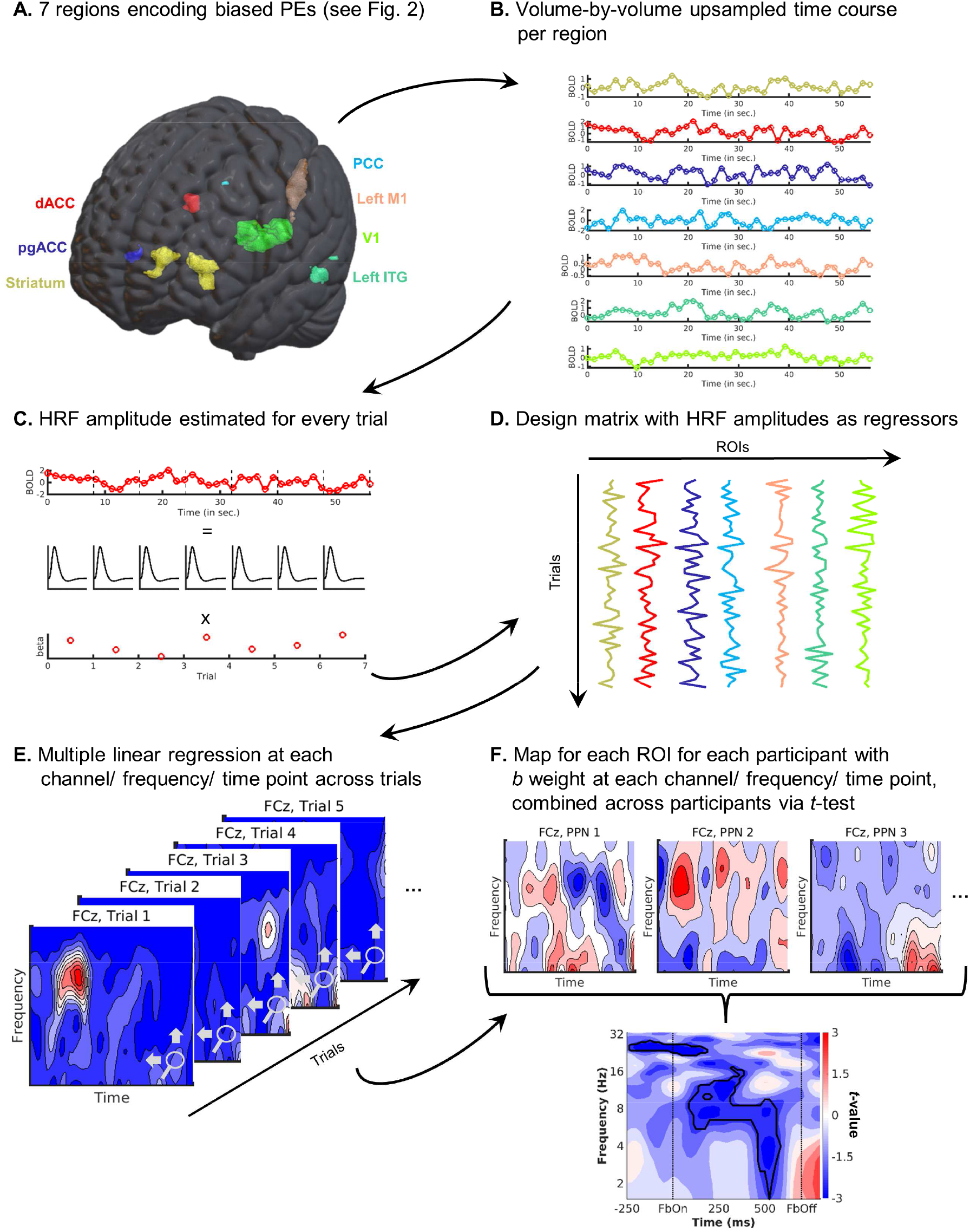
Graphical illustration of the fMRI-informed EEG analysis approach. **A.** Regions are identified to encode biased PEs via a model-based GLM on BOLD data (see Fig. 2 in the main text). **B.** The volume-by-volume time-series of the signal in each ROI is extracted and upsampled. **C.** Time series are epoched into trials and the HRF amplitude is estimated for every trial. **D.** HRF amplitudes in every ROI for every trial are combined into a design matrix. **E.** The design matrix is applied in a multiple linear regression for each participant at each channel, frequency, and time point across trials. **F.** Regressions yield a sensor-frequency-time map of *b* regression weights for each ROI for each participant. Maps are combined across participants using a one-sample *t*-test.

**Supplementary Figure 18.**
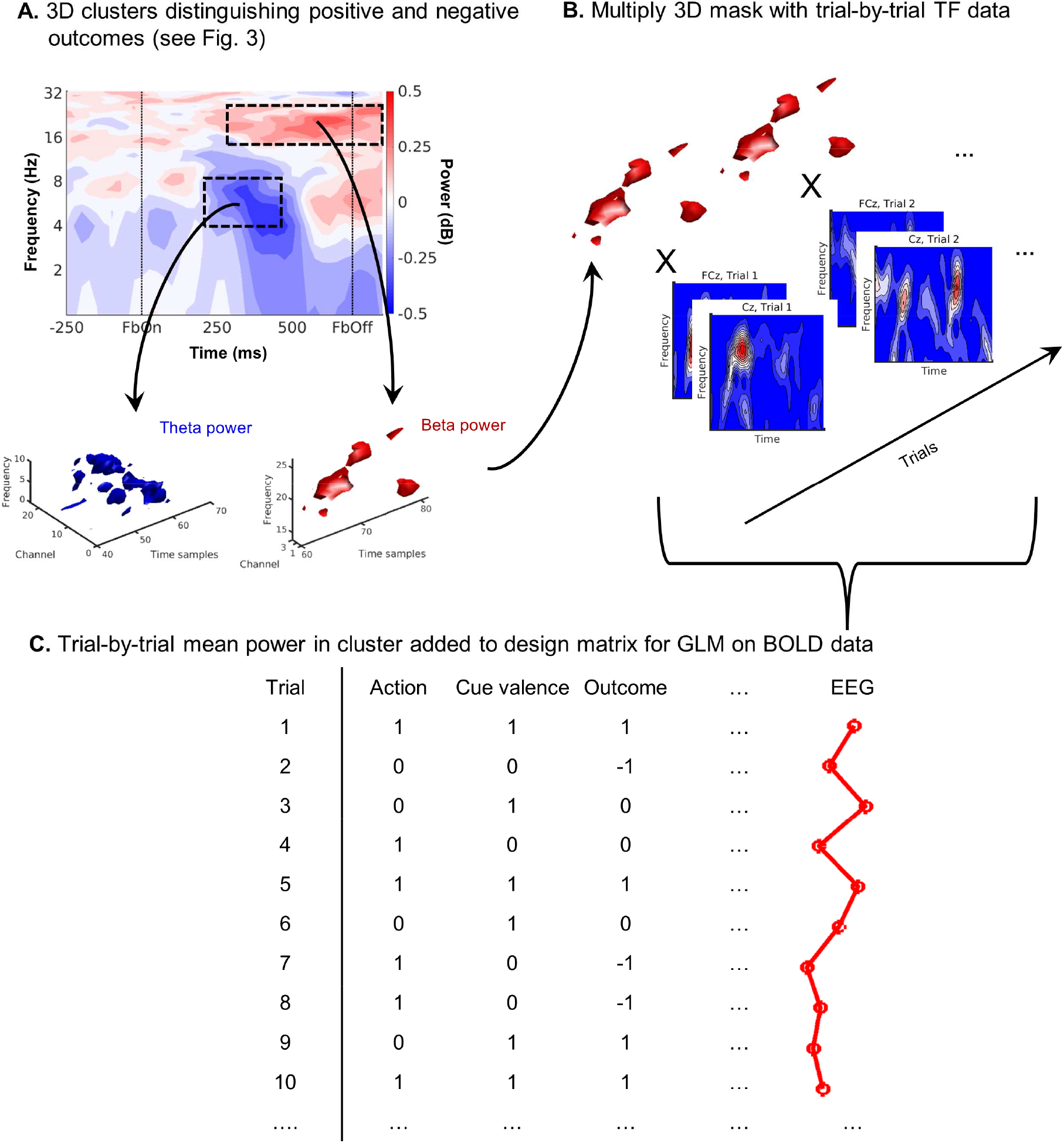
Graphical illustration of the EEG-informed fMRI analysis approach. **A.** 3D clusters of channel-frequency-time points where power significantly distinguishes trials with positive from trials with negative outcomes are identified via a cluster-based permutation test (see Fig. 3A in the main text). The *t*-values above a threshold |2| are retained, weights at all other grid points are set to zero. **B.** The 3D t-value cluster is multiplied with the trial-by-trial channel-frequency-time data, yielding a single average value of power in the cluster at each trial. **C.** Trial-by-trial average power in the cluster is added as a parametric regressor in the GLM on BOLD-data and fitted with FSL.

**Supplementary Figure 19.**
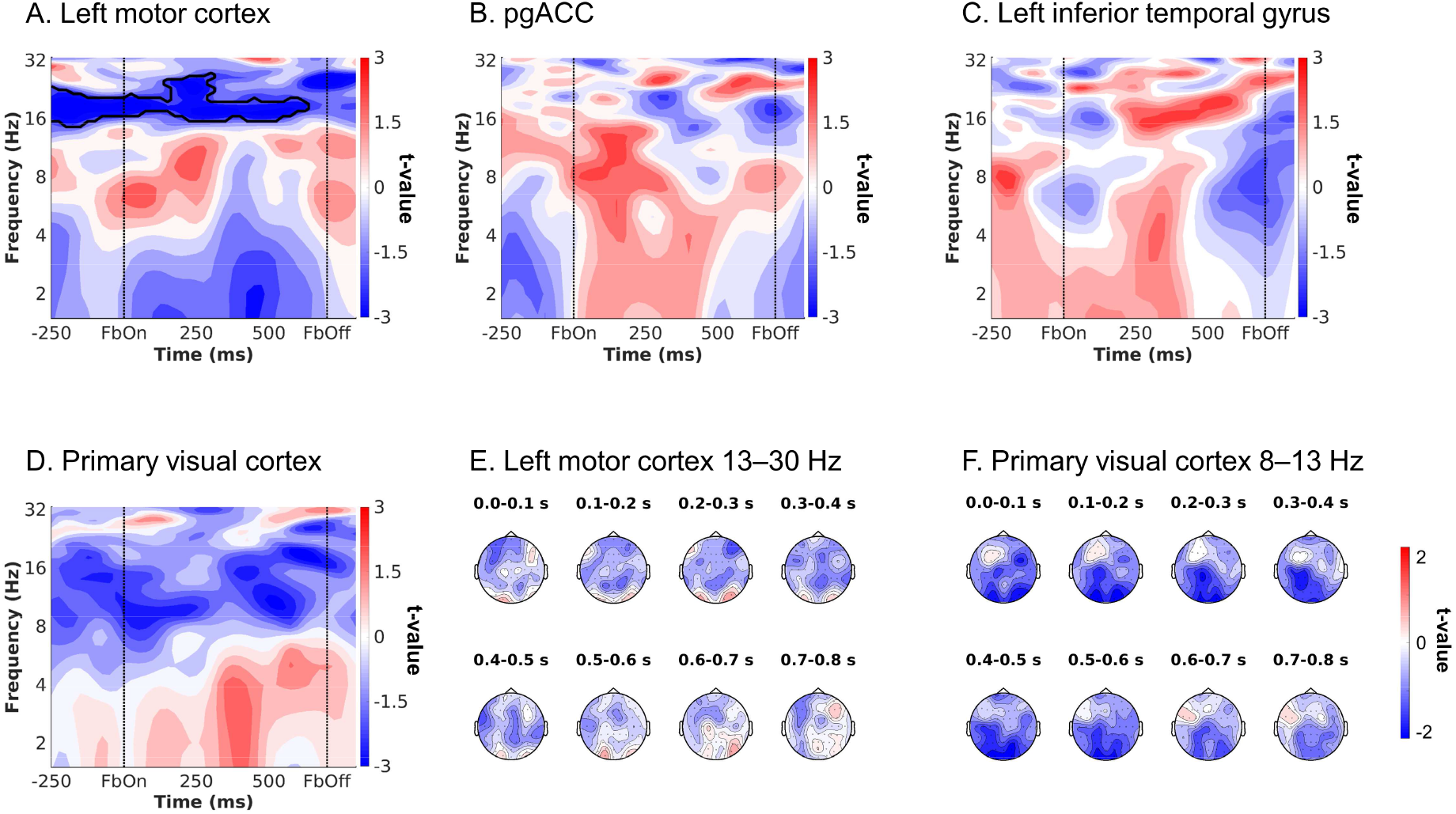
Supplementary fMRI-informed EEG results in the time-frequency domain. Unique temporal contributions of BOLD signal in (**A**) left motor cortex, (**B**) pgACC, (**C**) left ITG and (**D**) primary visual cortex to midfrontal EEG power. Group-level *t*-maps display the modulation of the EEG power over midfrontal electrodes (Fz/ FCz/ Cz) by trial-by-trial BOLD signal in the selected ROIs. There significant correlations between midfrontal EEG TF power in the beta range and left motor cortex BOLD signal (*p* = .002), but no significant midfrontal EEG correlates for BOLD signal from other ROIs. **E.** Topoplots displaying *t*-values of left motor cortex BOLD over the entire scalp between 13 and 30 Hz (beta band) in steps of 100 ms from 0 to 800 ms. There were significant negatively correlates over central electrodes, especially round 300–500 ms. **F.** Topoplot displaying *t*-values of primary visual cortex BOLD over the entire scalp between 8 and 13 Hz (alpha band) in steps of 100 ms from 0 to 800 ms. There were significantly negatively correlations over occipital electrodes throughout outcome presentation.

**Supplementary Figure 20.**
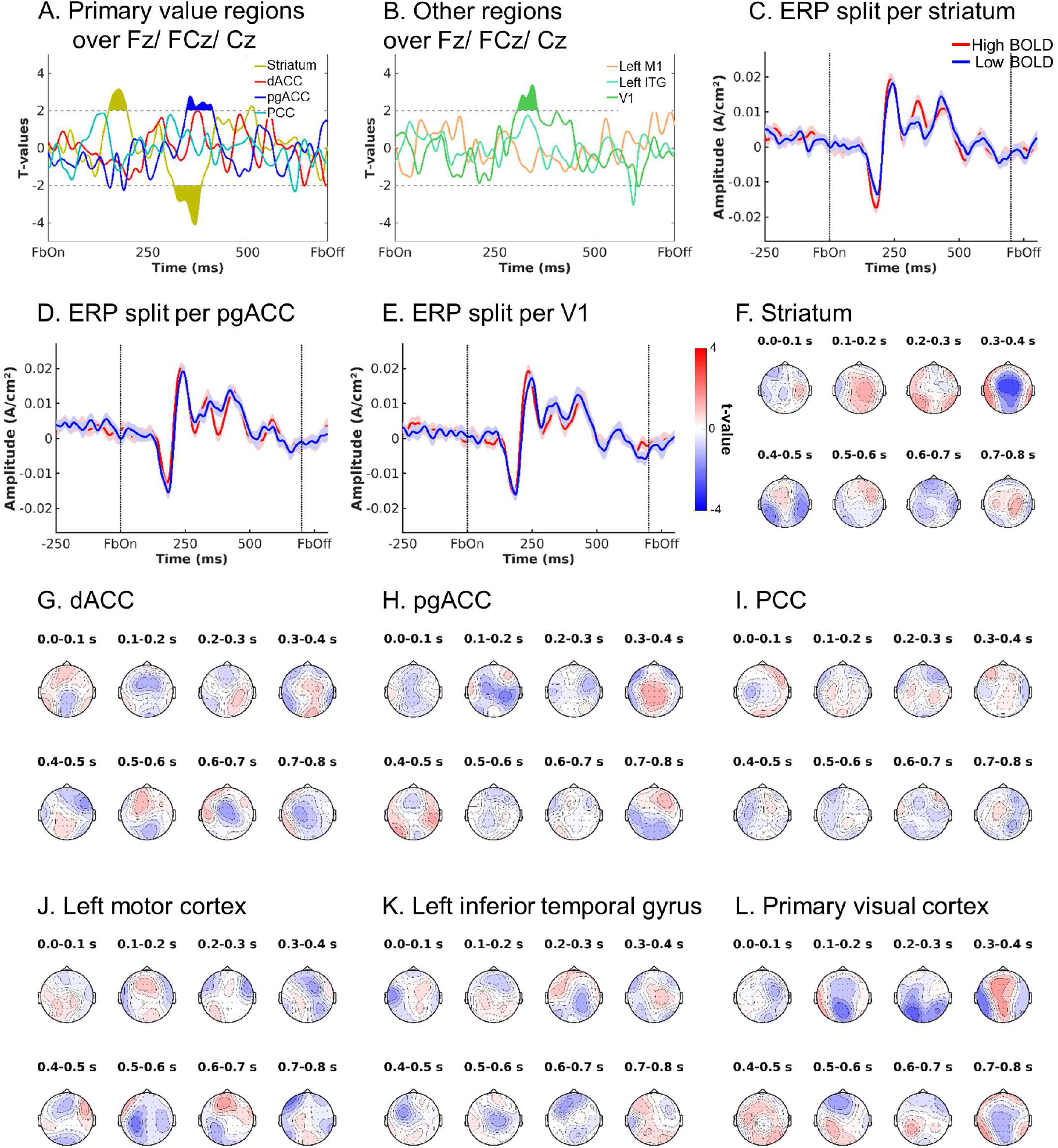
fMRI-informed EEG analyses in the time-domain. Group-level *t*-value time courses display the modulation of the EEG voltage over midfrontal electrodes (Fz/ FCz/ Cz) by trial-by-trial BOLD signal in the selected ROIs. **A.** Correlations between midfrontal voltage and trial-by-trial BOLD signal from core value regions, i.e., striatum, dACC, pgACC, and PCC. Striatal BOLD modulates the amplitude of the N1 and P3, while the P3 amplitude was also modulated by pgACC BOLD. **B.** Correlations between midfrontal voltage and trial-by-trial BOLD signal from other regions, i.e., left motor cortex, left inferior temporal gyrus, and primary visual cortex. Visual cortex BOLD modulates the amplitude of the P3, as well. **C-E.** Midfrontal voltage split up for high vs. low BOLD signal (median split) from regions significantly modulating voltage. Striatal BOLD modulated N1 and P2 amplitude, while pgACC BOLD and visual cortex BOLD modulated N2 (FRN) amplitude. **F-L.** Topoplots displaying *t*-values of correlations between midfrontal voltage and trial-by-trial BOLD for all regions in steps of 100 ms from 0 to 800 ms.

**Supplementary Figure 21.**
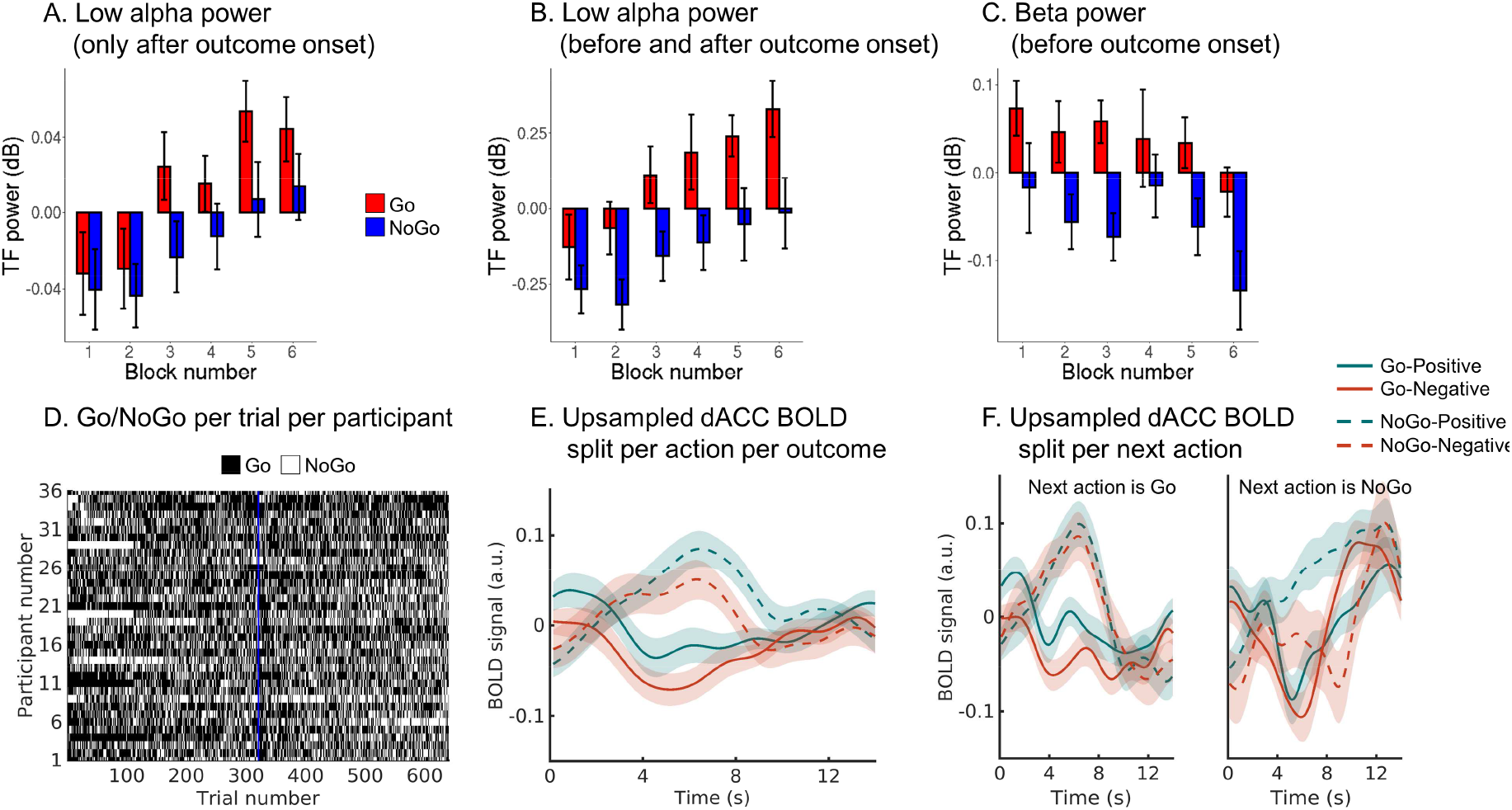
Control analyses excluding temporal confounds in midfrontal lower alpha band power and dACC BOLD. **A.** Mean midfrontal low alpha power (±SEM across participants) after outcome onset, (**B**) before and after outcome onset, and (**C**) beta power before outcome onset as a function of the performed action and block number (i.e., time on task). While low alpha power increases and beta power decreases over the time course of the task, power was always consistently higher for trials with Go than trials with NoGo responses, suggesting that action effects were not reducible to time on task. **D.** Response for each participant (rows) on each trial (columns). There was no noticeable change in the overall ratio of Go to NoGo responses over time. The vertical blue line indicates the start of the second session featuring new stimuli. **E.** Mean upsampled dACC BOLD signal (±SEM across participants) at the time of the outcome, split per performed action (Go/NoGo) and outcome valence (positive/negative). BOLD signal was higher after NoGo than Go responses. **F.** Same plot as (E), but split based on whether the next action was a Go (left panel) or an NoGo (right panel) response. Even if the next response was NoGo, BOLD signal was higher for trials with NoGo responses (on the current trial) than trials Go responses.

**Supplementary Figure 22.**
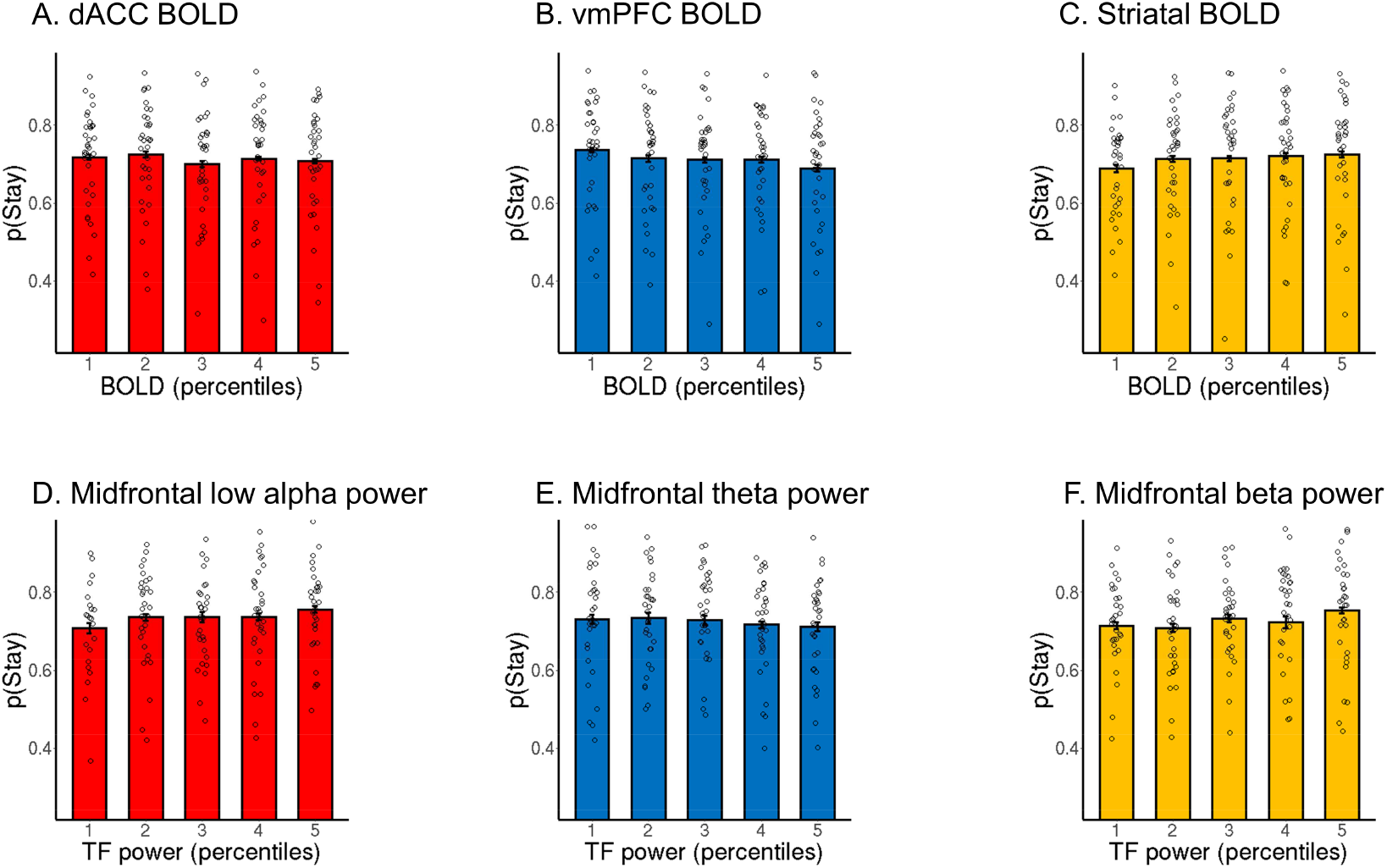
Probability of repeating the same response (“stay”) on the next cue encounter as a function of outcome-related BOLD and EEG signal. **A-C.** Probability of repeating the same action (“staying”) as a function of BOLD signal from (**A**) dACC, (**B**) vmPFC (cluster correlating with theta power in Fig. 5F), and (**C**) striatum (split into 5 bins). While dACC BOLD was not significantly linked to the probability to stay, high BOLD signal in vmPFC predicted a higher chance to switch to another action, while high BOLD signal in striatum predicted a higher probability of staying with the same action. **D-E.** Probability of staying as a function of midfrontal time-frequency power in the (**D**) low alpha, (**E**) theta/delta, and (**F**) beta range. Higher low alpha power and higher beta power predict a higher probability of staying with the same action, while higher theta power predicts a higher chance to switch to another action. Grey circles represent individual per condition-per-participant means. Error bars were very narrow (and thus hardly visible) and computed based on the Cousineau-Morey methods based on per-condition-per-participant means.

**Supplementary Table 1.** Full report of model of stay behavior. Mixed-effects logistic regression of stay vs. switch behavior (i.e., repeating vs. changing an action on the next occurrence of the same cue) as a function of performed action (Go vs. NoGo), outcome salience (salient: reward or punishment vs. neutral: no reward or no punishment), and outcome valence (positive: reward or no punishment vs. negative: no reward or punishment). Follow-up analyses were performed on trials with salient vs. neutral outcomes separately, and then separately based on Go vs. NoGo actions and salient vs. neutral outcomes. *P*-values were computed using likelihood ratio tests using the *mixed*-function (option “LRT”) from package *afex*.

| Effect | $\chi^2$ | Df | p-value |
| --- | --- | --- | --- |
| Action | 0.01 | 1 | .924 |
| Salience | 5.15 | 1 | .021 |
| Valence | 45.59 | 1 | < .001 |
| Action x Salience | 0.12 | 1 | .728 |
| Action x Valence | 3.24 | 1 | .067 |
| Salience x Valence | 30.95 | 1 | < .001 |
| Action x Valence x Salience | 19.73 | 1 | < .001 |
| *Salient outcomes only:* |  |  |  |
| Action | 0.01 | 1 | .960 |
| Valence | 46.36 | 1 | < .001 |
| Action x Valence | 17.80 | 1 | < .001 |
| *Neutral outcomes only:* |  |  |  |
| Action | .102 | 1 | .750 |
| Valence | .830 | 1 | .362 |
| Action x Valence | 12.32 | 1 | < .001 |
| *Go with salient outcomes only:* |  |  |  |
| Valence | 53.93 | 1 | < .001 |
| *NoGo with salient outcomes only:* |  |  |  |
| Valence | 18.23 | 1 | < .001 |
| *Go with neutral outcomes only:* |  |  |  |
| Valence | 0.13 | 1 | .050 |
| *NoGo with neutral outcomes only:* |  |  |  |
| Valence | 7.21 | 1 | .007 |

**Supplementary Table 2.** Model parameters for fitted models. Mean [minimum – maximum] of participant-level parameter estimates in model space, fitted with hierarchical Bayesian inference (only the respective model included in the fitting process). Model frequency and protected exceedance probability were based on a model comparison that involves models M1-M6. Note that Fig. 2 in the main text does not include M6.

|  | M1 | M2 | M3 | M4 | M5<br>(Asymmetric<br>pathways) | M6<br>(Action<br>priming) |
| --- | --- | --- | --- | --- | --- | --- |
| Mean log model evidence | -609.30 | -597.95 | -554.46 | -532.40 | -528.13 | -540.84 |
| Model frequency | 0 | 0.0278 | 0 | 0.0488 | 0.6815 | 0.2419 |
| Protected exceedance probability | 0 | 0 | 0 | 0 | .9970 | .0030 |
| $\rho$ | 7.75<br>[0.53 – 38.68] | 6.81<br>[0.48 – 37.74] | 6.38<br>[0.49 – 35.71] | 10.05<br>[1.26 – 40.60] | 9.41<br>[0.98 – 31.22] | 6.64<br>[0.71 – 22.83] |
| $\varepsilon_0$ | 0.17<br>[0.002 – 0.77] | 0.20<br>[0.003 – 0.82] | 0.21<br>[0.003 – 0.85] | 0.09<br>[0.003 – 0.38] | 0.08<br>[0.003 – 0.41] | 0.039<br>[0.003 – 0.11] |
| $b$ | | -0.05<br>[-1.23 – 0.82] | -0.01<br>[-1.23 – 1.09] | 0.13<br>[-1.16 – 1.03] | 0.14<br>[-1.18 – 1.10] | 0.16<br>[-1.22 – 1.40] |
| $\pi$ | | | 0.77<br>[-0.78 – 3.73] | | 0.17<br>[-1.25 – 2.70] | -1.11<br>[-3.29 – 1.23] |
| $\varepsilon_{\text{rewarded Go}} (\varepsilon_0 + \kappa)$ | | | | 0.749<br>[0.29 – 0.99] | 0.833<br>[0.43 – 0.99] | |
| $\varepsilon_{\text{punished NoGo}} (\varepsilon_0 - \kappa)$ | | | | 0.001<br>[0.001 – 0.02] | 0.003<br>[0.001 – 0.09] | |
| $\varepsilon_{\text{salient Go}}$ | | | | | | 0.49<br>[0.05 – 0.90] |

**Supplementary Table 3.**
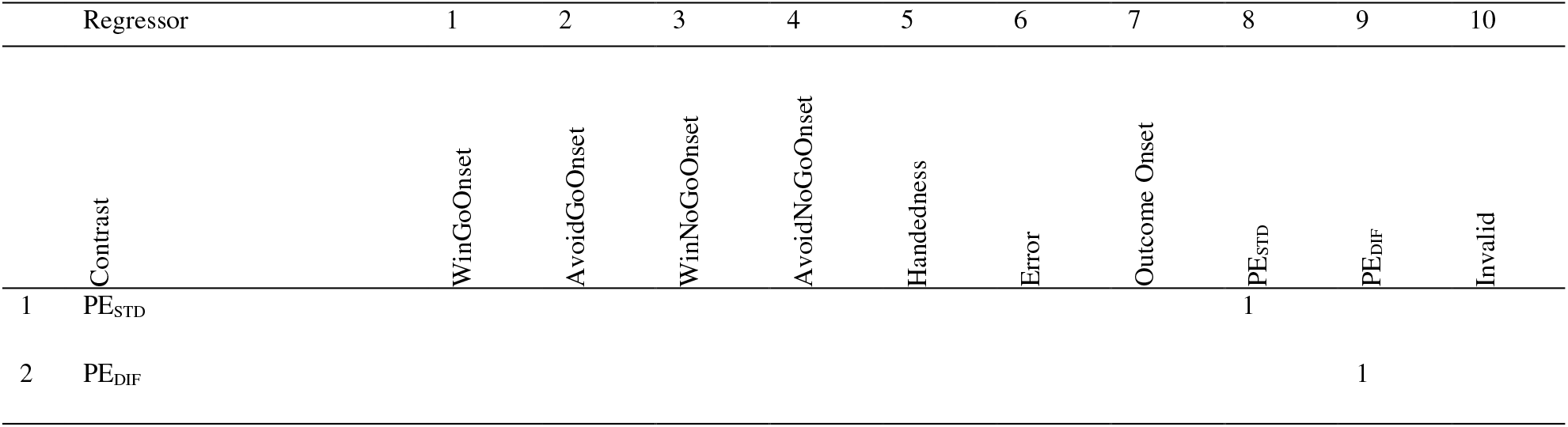
BOLD-GLM with parametric modulation by standard and biased prediction errors. Explanation of regressors: WinGoOnset: for every trial with Win cue and Go action, at cue onset, duration 1, value +1. AvoidGoOnset: for every trial with Avoid cue and Go action, at cue onset, duration 1, value +1. WinNoGoOnset: for every trial with Win cue and NoGo action, at cue onset, duration 1, value +1. AvoidNoGoOnset: for every trial with Avoid cue and NoGo action, at cue onset, duration 1, value +1. Handedness: for every trial, at cue onset, duration 1, value +1 for left hand response, 0 for NoGo 10 response, −1 for right hand response. Error: for every trial, at cue onset, duration 1, value +1 for incorrect response, 0 for correct response. OutcomeOnset: for every trial, at outcome onset, duration 1, value +1 for every trial. PE_STD_: for every trial, at outcome onset, duration 1, value is the demeaned PE times learning rate for model M1. PE_DIF_: for every trial, at outcome onset, duration 1, value is the demeaned difference between (PE times learning rate) for model M1 and (PE times learning rate) for model M5. Invalid: for trials where uninstructed button was pressed, at outcome onset, duration 1, value 1.

**Supplementary Table 4.** BOLD-GLM with response-locked and outcome-locked categorical regressors. Explanation of regressors: GoReward: for every trial with Go action and reward obtained, at outcome onset, duration 1, value +1. GoNoReward: for every trial with Go action and no reward obtained, at outcome onset, duration 1, value +1. GoNoPunishment: for every trial with Go action and no punishment obtained, at outcome onset, duration 1, value +1. GoPunishment: for every trial with Go action and punishment obtained, at outcome onset, duration 1, value +1. NoGoReward: for every trial with NoGo action and reward obtained, at outcome onset, duration 1, value +1. NoGoNoR eward: for every trial with NoGo action and no reward obtained, at outcome onset, duration 1, value +1. NoGoNoPunishment: for every trial with NoGo action and no punishment obtained, at outcome onset, duration 1, value +1. NoGoPunishment: for every trial with NoGo action and punishment obtained, at outcome onset, duration 1, value +1. LeftHand: for very trial with left hand response, at response onset, duration 1, value + 1. RightHand: for very trial with right hand response, at response onset, duration 1, value + 1. Error: for every trial, at cue onset, duration 1, value +1 for incorrect response, 0 for correct response. OutcomeOnset: for every trial, at outcome onset, duration 1, value +1 for every trial. Invalid: for trials where uninstructed button was pressed, at outcome onset, duration 1, value 1.

| Regressors |  | 1 | 2 | 3 | 4 | 5 | 6 | 7 | 8 | 9 | 10 | 11 | 12 | 13 |
| --- | --- | --- | --- | --- | --- | --- | --- | --- | --- | --- | --- | --- | --- | --- |
| Contrast |  | GoReward | GoNoReward | GoNoPunishment | GoPunishment | NoGoReward | NoGoNoReward | NoGoNoPunishment | NoGoPunishment | LeftHand | RightHand | Error | OutcomeOnset | Invalid |
| 1 | Valence | 1 | -1 | 1 | -1 | 1 | -1 | 1 | -1 |  |  |  |  |  |
| 2 | Action | 1 | 1 | 1 | 1 | -1 | -1 | -1 | -1 |  |  |  |  |  |
| 3 | Hand Sum |  |  |  |  |  |  |  |  | 1 | 1 |  |  |  |
| 4 | Hand Dif |  |  |  |  |  |  |  |  | 1 | -1 |  |  |  |

**Supplementary Table 5:**
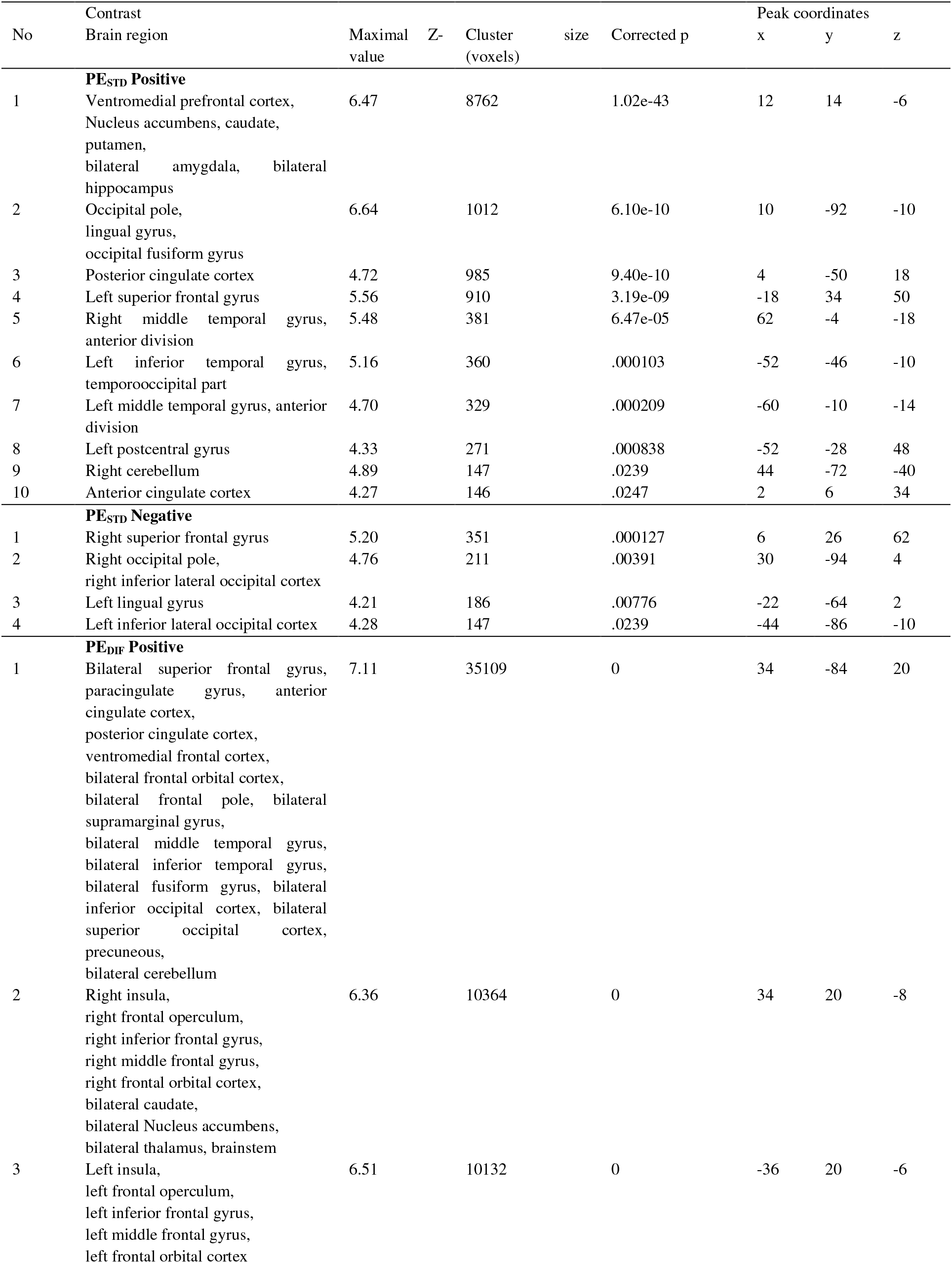
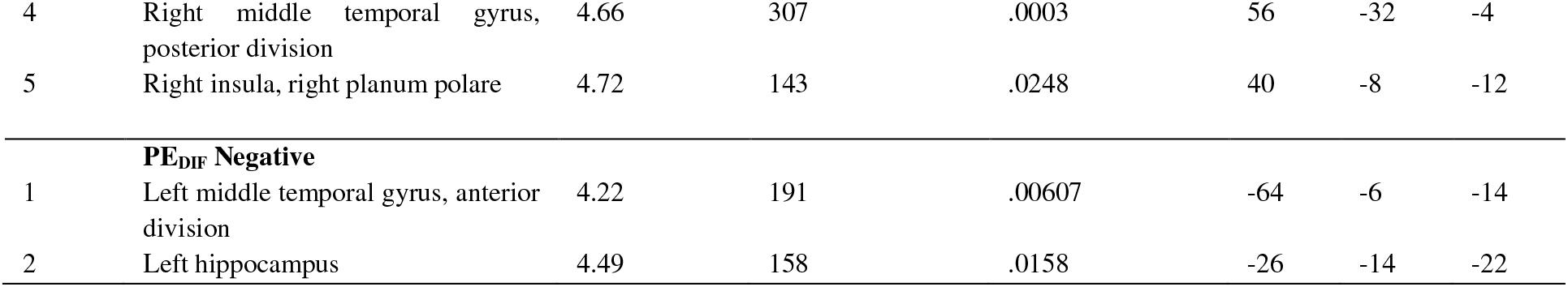
Significant clusters in BOLD-GLM with parametric modulation by standard and biased prediction errors.

**Supplementary Table 6:**
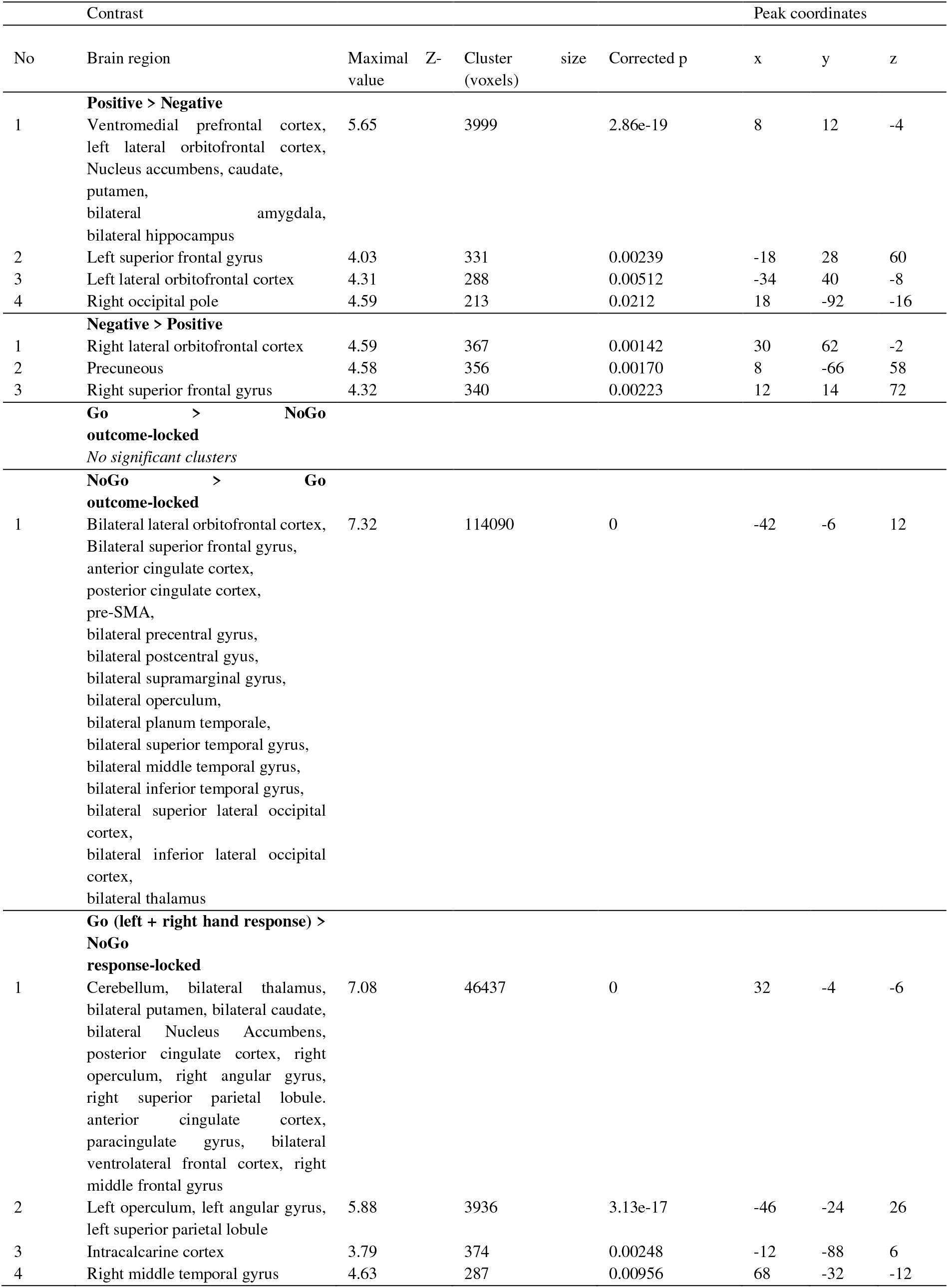
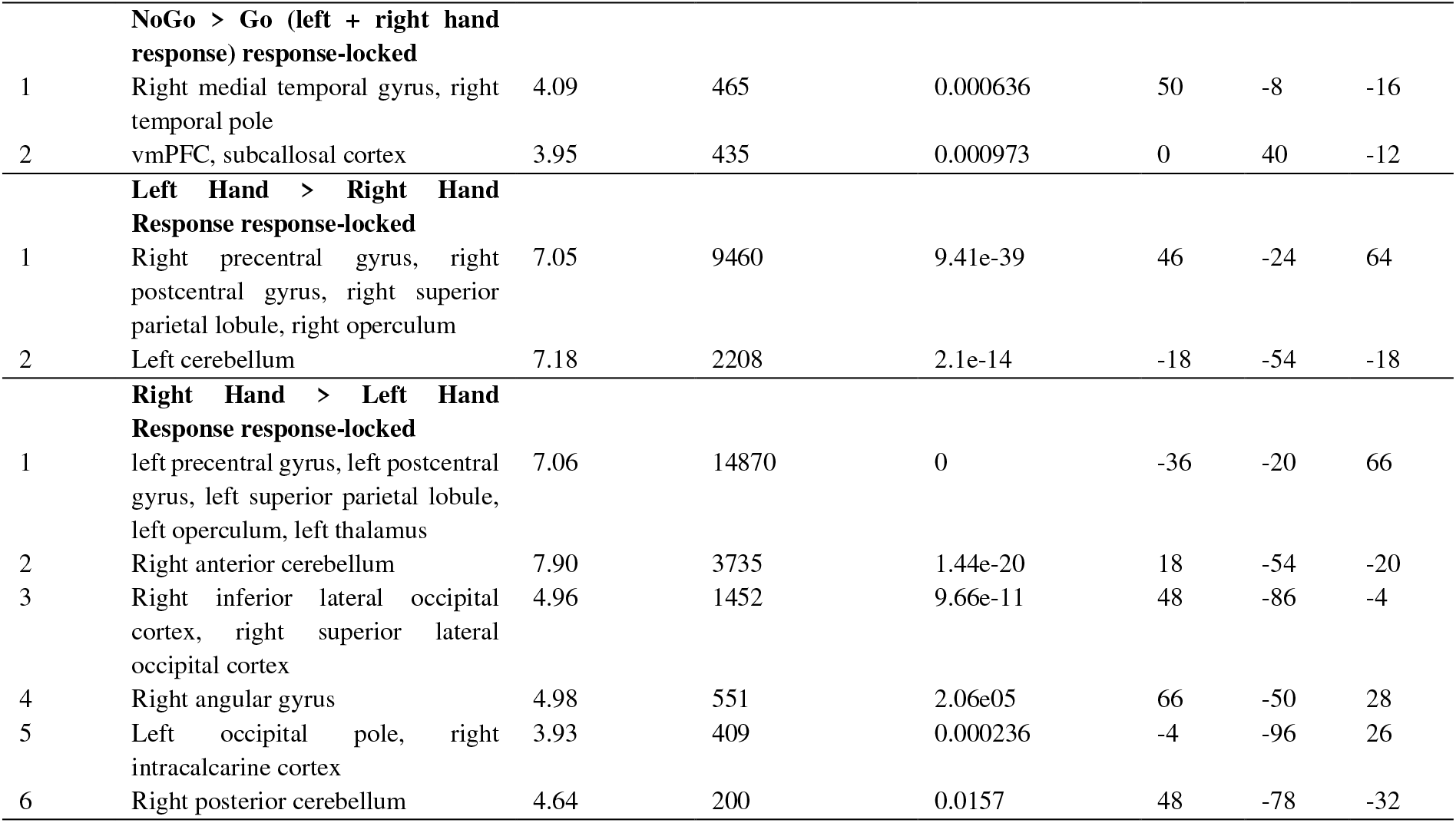
Significant clusters in BOLD-GLM with response-locked and outcome-locked categorical regressors.

**Supplementary Table 7:**
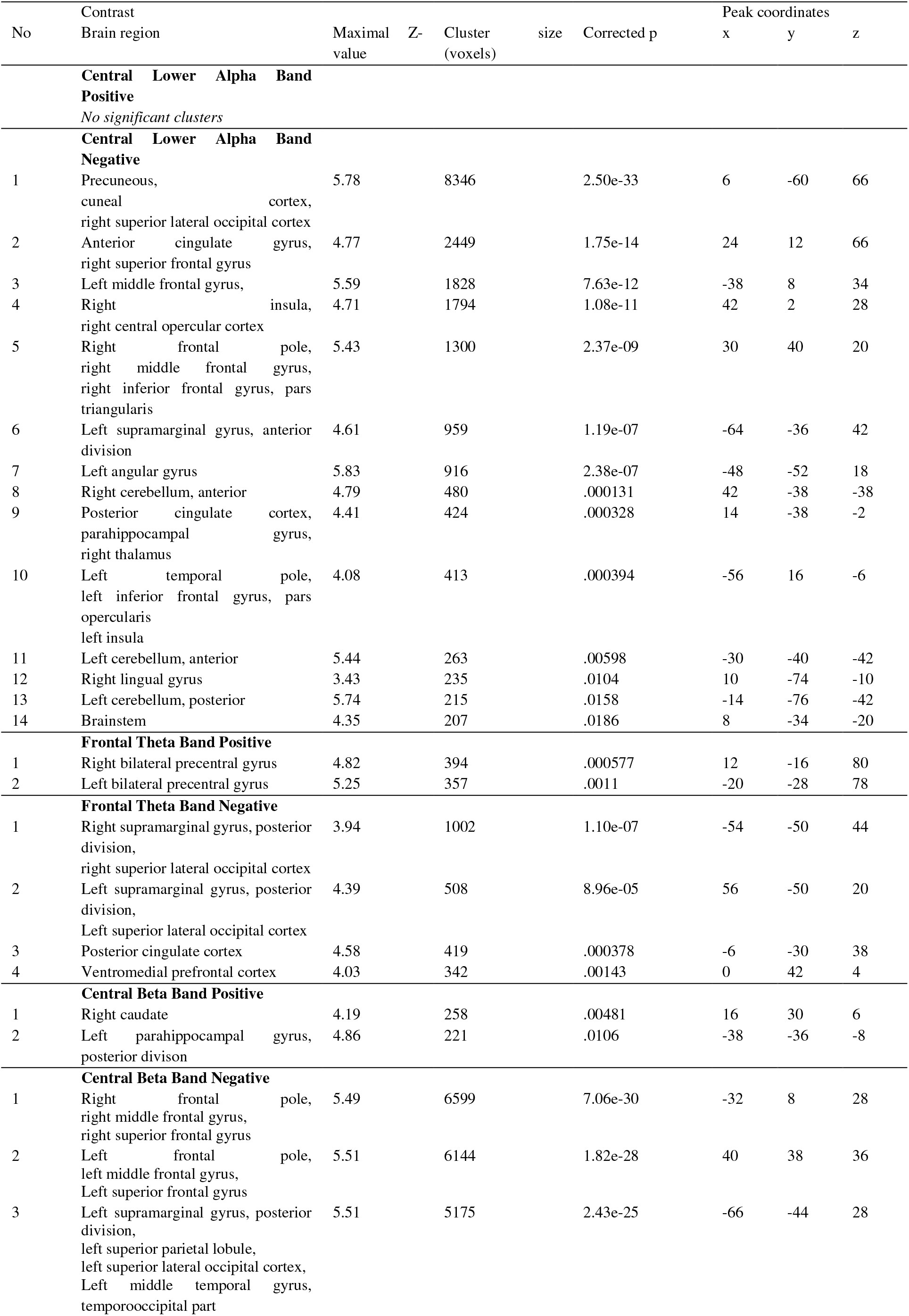
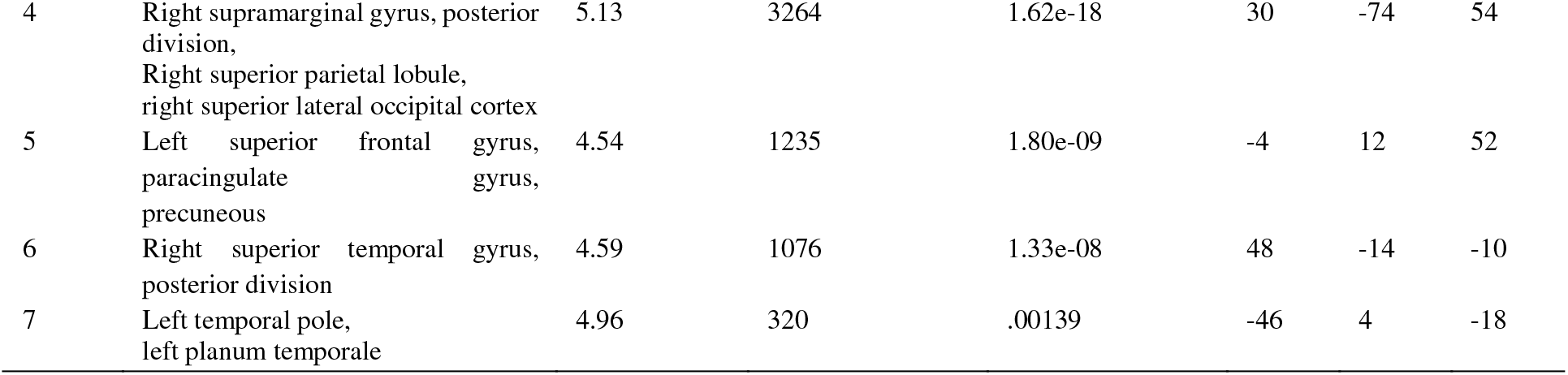
Significant clusters in BOLD-GLM with EEG regressors.

| No | Contrast<br>Brain region |  |  |  | Maximal<br>value | Z- | Cluster<br>(voxels) | size | Corrected p | Peak coordinates<br>x y z |  |  |
| --- | --- | --- | --- | --- | --- | --- | --- | --- | --- | --- | --- | --- |
|  | **Central Lower Alpha Band**<br>**Positive** |  |  |  |  |  |  |  |  |  |  |  |
|  | *No significant clusters* |  |  |  |  |  |  |  |  |  |  |  |
|  | **Central Lower Alpha Band**<br>**Negative** |  |  |  |  |  |  |  |  |  |  |  |
| 1 | Precuneous, cuneal cortex, right superior lateral occipital cortex |  |  |  | 5.78 |  | 8346 |  | 2.50e-33 | 6 | -60 | 66 |
| 2 | Anterior cingulate gyrus, right superior frontal gyrus |  |  |  | 4.77 |  | 2449 |  | 1.75e-14 | 24 | 12 | 66 |
| 3 | Left middle frontal gyrus, |  |  |  | 5.59 |  | 1828 |  | 7.63e-12 | -38 | 8 | 34 |
| 4 | Right insula, right central opercular cortex |  |  |  | 4.71 |  | 1794 |  | 1.08e-11 | 42 | 2 | 28 |
| 5 | Right frontal pole, right middle frontal gyrus, right inferior frontal gyrus, pars triangularis |  |  |  | 5.43 |  | 1300 |  | 2.37e-09 | 30 | 40 | 20 |
| 6 | Left supramarginal gyrus, anterior division |  |  |  | 4.61 |  | 959 |  | 1.19e-07 | -64 | -36 | 42 |
| 7 | Left angular gyrus |  |  |  | 5.83 |  | 916 |  | 2.38e-07 | -48 | -52 | 18 |
| 8 | Right cerebellum, anterior |  |  |  | 4.79 |  | 480 |  | .000131 | 42 | -38 | -38 |
| 9 | Posterior cingulate cortex, parahippocampal gyrus, right thalamus |  |  |  | 4.41 |  | 424 |  | .000328 | 14 | -38 | -2 |
| 10 | Left temporal pole, left inferior frontal gyrus, pars opercularis left insula |  |  |  | 4.08 |  | 413 |  | .000394 | -56 | 16 | -6 |
| 11 | Left cerebellum, anterior |  |  |  | 5.44 |  | 263 |  | .00598 | -30 | -40 | -42 |
| 12 | Right lingual gyrus |  |  |  | 3.43 |  | 235 |  | .0104 | 10 | -74 | -10 |
| 13 | Left cerebellum, posterior |  |  |  | 5.74 |  | 215 |  | .0158 | -14 | -76 | -42 |
| 14 | Brainstem |  |  |  | 4.35 |  | 207 |  | .0186 | 8 | -34 | -20 |
|  | **Frontal Theta Band Positive** |  |  |  |  |  |  |  |  |  |  |  |
| 1 | Right bilateral precentral gyrus |  |  |  | 4.82 |  | 394 |  | .000577 | 12 | -16 | 80 |
| 2 | Left bilateral precentral gyrus |  |  |  | 5.25 |  | 357 |  | .0011 | -20 | -28 | 78 |
|  | **Frontal Theta Band Negative** |  |  |  |  |  |  |  |  |  |  |  |
| 1 | Right supramarginal gyrus, posterior division, right superior lateral occipital cortex |  |  |  | 3.94 |  | 1002 |  | 1.10e-07 | -54 | -50 | 44 |
| 2 | Left supramarginal gyrus, posterior division, Left superior lateral occipital cortex |  |  |  | 4.39 |  | 508 |  | 8.96e-05 | 56 | -50 | 20 |
| 3 | Posterior cingulate cortex |  |  |  | 4.58 |  | 419 |  | .000378 | -6 | -30 | 38 |
| 4 | Ventromedial prefrontal cortex |  |  |  | 4.03 |  | 342 |  | .00143 | 0 | 42 | 4 |
|  | **Central Beta Band Positive** |  |  |  |  |  |  |  |  |  |  |  |
| 1 | Right caudate |  |  |  | 4.19 |  | 258 |  | .00481 | 16 | 30 | 6 |
| 2 | Left parahippocampal gyrus, posterior division |  |  |  | 4.86 |  | 221 |  | .0106 | -38 | -36 | -8 |
|  | **Central Beta Band Negative** |  |  |  |  |  |  |  |  |  |  |  |
| 1 | Right frontal pole, right middle frontal gyrus, right superior frontal gyrus |  |  |  | 5.49 |  | 6599 |  | 7.06e-30 | -32 | 8 | 28 |
| 2 | Left frontal pole, left middle frontal gyrus, Left superior frontal gyrus |  |  |  | 5.51 |  | 6144 |  | 1.82e-28 | 40 | 38 | 36 |
| 3 | Left supramarginal gyrus, posterior division, left superior parietal lobule, left superior lateral occipital cortex, Left middle temporal gyrus, temporooccipital part |  |  |  | 5.51 |  | 5175 |  | 2.43e-25 | -66 | -44 | 28 |

|  |  |  |  |  |  |  |  |
| --- | --- | --- | --- | --- | --- | --- | --- |
| 4 | Right supramarginal gyrus, posterior division,<br>Right superior parietal lobule,<br>right superior lateral occipital cortex | 5.13 | 3264 | 1.62e-18 | 30 | -74 | 54 |
| 5 | Left superior frontal gyrus,<br>paracingulate gyrus,<br>precuneous | 4.54 | 1235 | 1.80e-09 | -4 | 12 | 52 |
| 6 | Right superior temporal gyrus,<br>posterior division | 4.59 | 1076 | 1.33e-08 | 48 | -14 | -10 |
| 7 | Left temporal pole,<br>left planum temporale | 4.96 | 320 | .00139 | -46 | 4 | -18 |

